# The changing mouse embryo transcriptome at whole tissue and single-cell resolution

**DOI:** 10.1101/2020.06.14.150599

**Authors:** Peng He, Brian A. Williams, Diane Trout, Georgi K. Marinov, Henry Amrhein, Libera Berghella, Say-Tar Goh, Ingrid Plajzer-Frick, Veena Afzal, Len A. Pennacchio, Diane E. Dickel, Axel Visel, Bing Ren, Ross C. Hardison, Yu Zhang, Barbara J. Wold

## Abstract

In mammalian embryogenesis differential gene expression gradually builds the identity and complexity of each tissue and organ system. We systematically quantified mouse polyA-RNA from embryo day E10.5 to birth, sampling 17 whole tissues, enhanced with single-cell measurements for the developing limb. The resulting developmental transcriptome is globally structured by dynamic cytodifferentiation, body-axis and cell-proliferation gene sets, characterized by their promoters’ transcription factor (TF) motif codes. We decomposed the tissue-level transcriptome using scRNA-seq and found that neurogenesis and haematopoiesis dominate at both the gene and cellular levels, jointly accounting for 1/3 of differential gene expression and over 40% of identified cell types. Integrating promoter sequence motifs with companion ENCODE epigenomic profiles identified a promoter de-repression mechanism unique to neuronal expression clusters and attributable to known and novel repressors. Focusing on the developing limb, scRNA-seq identified 25 known and candidate novel cell types, including progenitor and differentiating states with computationally inferred lineage relationships. We extracted cell type TF networks and complementary sets of candidate enhancer elements by de-convolving whole-tissue IDEAS epigenome chromatin state models. These ENCODE reference data, computed network components and IDEAS chromatin segmentations, are companion resources to the matching epigenomic developmental matrix, available for researchers to further mine and integrate.

## Introduction

Hierarchical transcription programs regulate mammalian histogenesis, a spatiotemporally coordinated process of changing cell identities, numbers and locations ^1^. Contemporary RNA-seq time-courses can quantify expression trajectories comprehensively, including those of genes encoding the transcriptional regulators that drive patterning, cell-type specification and differentiation. Here we report a systematic mapping of the mouse polyadenylated RNA transcriptome, tracking 12 major tissues from E10.5 to birth (P0) (Fig. 1a, b and Extended Data Fig. 1l) and covering much of organogenesis and histogenesis. Crucial for regulatory genomics analysis and modeling, these data are part of the ENCODE Consortium mouse embryo project, which provides companion microRNA-seq, DNA methylation, histone mark ChIP-seq, and chromatin accessibility datasets for the same sample matrix ^2^. To better interpret the core sample set, we added 5 additional organs at P0, sampling 17 tissues in all. As these whole-tissue data are intended for community use, including integration with high resolution single-cell transcriptomes, we applied a widely used RNA-seq method (SMART-seq) that is robust at both bulk sample and single-cell scales ^3^ and has been used for other single-cell RNA-seq (scRNA-seq) experiments in ENCODE ^4^ (https://www.encodeproject.org/) and elsewhere (e.g. Tabula Muris) ^5^.

Single-cell RNA-seq data are increasingly the means for discovering and defining constituent cell-types and states that comprise complex tissues such as those in our bulk mRNA seq matrix ^6–9^. For embryogenesis and regenerating systems in particular, scRNA-seq further promises to address longstanding questions about the nature and number of intermediate cell types in a developmental lineage and the regulatory mechanisms that govern transitions between them. Finally, scRNA-seq data offer an important input for gene network modeling by unambiguously assigning to an individual cell (or cell group) its transcription factor repertoire. Different contemporary scRNA-seq methods have complementary strengths, with some (e.g. Fluidigm SMART-seq) assaying relatively modest numbers of cells with high transcript detection efficiency and RNA isoform discriminating coverage while others (e.g. 10x Genomics) capture larger cell numbers at lower transcript detection efficiency and without isoform or promoter use information ^5,10–12^. We present here an ENCODE scRNA-seq resource containing both data-types for the developing forelimb, a tissue series not represented in the Tabula Muris project ^5^. We then identify limb cell lineages and stages within them, and extract their corresponding cell-type marker gene sets, TF networks, and promoter and distal candidate regulatory elements with their TF binding motifs. The higher sensitivity Fluidigm data-type additionally uncovered developmentally precocious low-level transcription of lineage specific regulators that further support computed lineage inference models.

An emerging goal for developmental genomics is to comprehensively chart the cis- and trans-acting regulatory codes of embryogenesis with single-cell resolution. In this direction, we use the limb scRNA-seq data to deconvolve integrative *cis*-element enhancer state (IDEAS) models ^13,14^ based on whole-tissue ENCODE epigenomic data. The resulting collection of candidate active and poised enhancer elements, parsed for cell type and stage, complements matching trans-acting TF networks. All primary RNA-seq data and processed quantifications for tissue-level and single-cell datasets are available from the ENCODE portal (https://www.encodeproject.org).

## Results and Discussion

The developmental timespan from mid-gestation (day E10.5) to birth (P0) encompasses much of histogenesis and organogenesis in the mouse (Fig. 1a; Ext. Fig.1l). The timecourse transcriptomes clustered according to their respective tissue identities and, within tissues, by developmental time, as shown by principal component analysis (PCA) (Fig. 1b; Supplementary Data 2), *t*-distributed stochastic neighbor embedding (*t*-SNE) (Extended Data Fig. 4a), and hierarchical clusterings (Fig. 1c and Extended Data Fig. 4b). Overall, this polyA RNA transcriptome encompasses 84% of all protein coding genes and 44% of lncRNAs, with 15,644 genes differing by ≥10-fold across the matrix and 9,085 more uniformly expressed genes that include housekeeping activities and structures (Extended Data Figs. 1a; 3a). Relative to the FANTOM5 mouse resource ^10^ (http://fantom.gsc.riken.jp/) which covers many of the same tissues and stages and is based on CAGE promoter data, we detected 97% of its 13,999 protein coding genes, plus 5,035 that are novel in our data (Extended Data Fig. 1m).

### Haematopoiesis and neurogenesis polarize the developmental transcriptome

Neurogenesis and haematopoiesis dominate the global data structure, with transcriptomes from these systems occupying opposite ends of the first two principal components (PCs) (Fig. 1 b,c). Nearly 1/5 of the expressed transcriptome (∼5000 genes) unambiguously defines this differential axis, which was robust to the choice of quantification units (FPKM or TPM Extended Data Fig. 4f,g) and to tissue representation (Extended Data Fig. 4d,e). Because whole-tissue data sum over all constituent cell types, their transcriptomes obscure underlying cell identities and relative cell proportions that are fundamental in histogenesis (Fig. 1a). We therefore projected cell type marker genes and cell identities from a recent single-cell mouse whole embryo survey ^11^ into our transcriptome structure (Fig. 1d). This showed that the high-complexity CNS and haematopoetic gene profiles correspond to high cellular diversity defined by the single cell decomposition, with 40% of cell types mapping to CNS and haematopoetic gene clusters. Focusing in, the single-cell projection further identifies tissue level expression of numerous gene clusters or sub-clusters attributable to specific cell type contributions (e.g. ependymal cells; neural progenitor cells, cardiomyocytes) (Fig. 1d, black boxes).

### Temporal drivers

Developmental changes were expected at the tissue level, but we did not know in advance what genes and functions would most prominently define the temporal axis, nor how they would distribute in tissue/organ/cell space. Analysis across all tissues found three classes of temporal drivers:

#### 1) Universal

PC3 captured a strong global time component (Fig. 1b, *z*-axis) explained at the gene level by widespread diminution in cell proliferation machinery and early erythroid markers (Extended data Fig. 3c). The top 100 PC3 positive-loading genes are highly enriched for mitotic cell cycle components (GO *p* = 3e-13) that map to expression Cluster 21 (Fig. 1c and Extended Data Fig. 2–21) which, in turn, maps to the stromal and early erythroid cell types of Cao et al (Fig. 1d, red boxes). Furthermore, their stromal cell marker set is itself enriched in cell cycle genes (*p* =1.8e-13 “Cell cycle”) and the reverse is also true. Thus the universal transcriptome time axis of PC3 can be explained, at least in part, by a gradual system-wide disappearance of circulating primitive erythrocytes and a decrease in the relative proportion of proliferating stromal cells across many tissues and organs.

**Figure 1:**
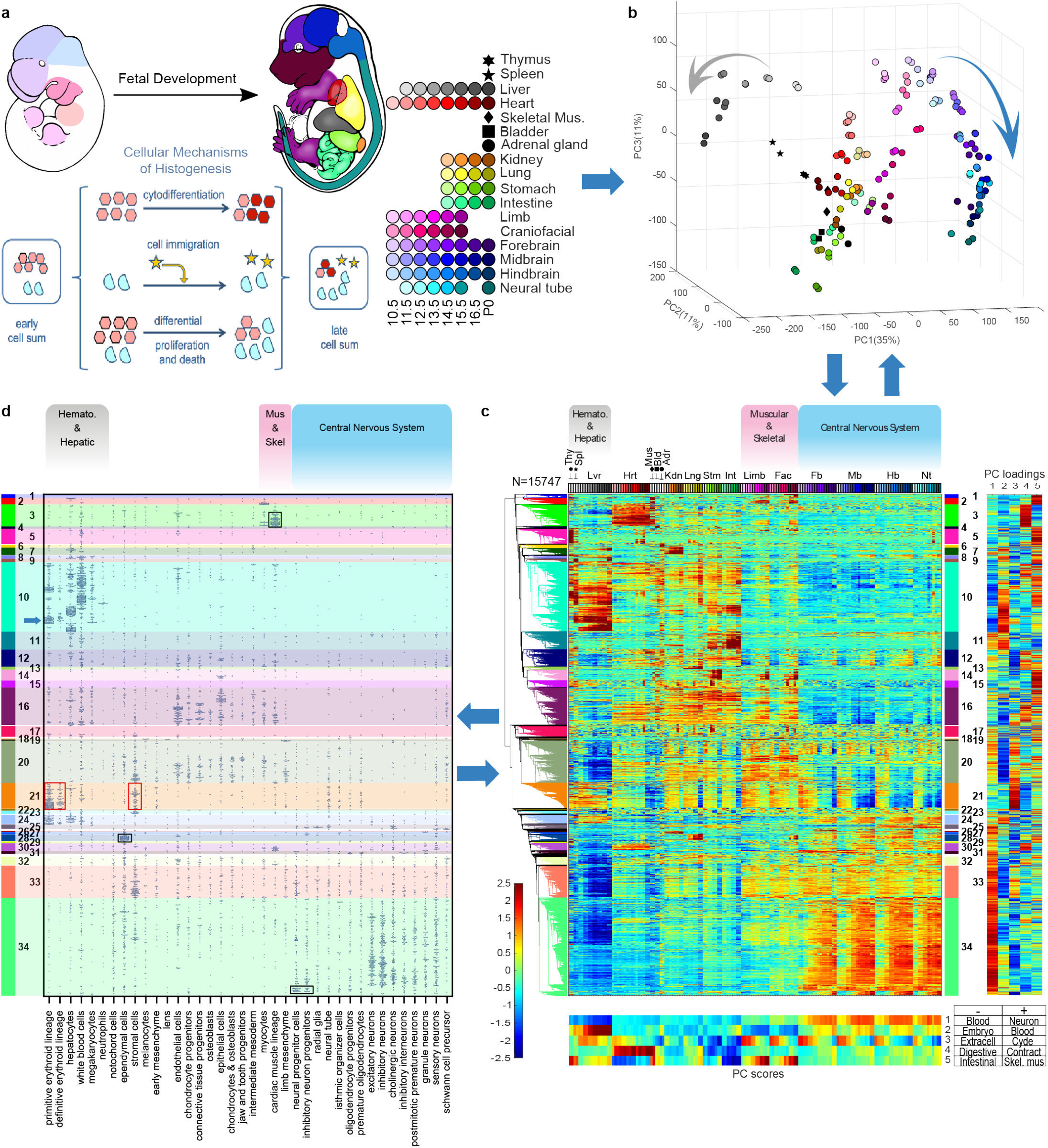
Whole-tissue polyA RNA transcriptome structure with cell type decomposition. (a) Schematic of E10.5 and E15.5 embryos shows the color key for organ identity and developmental stage across the timespan of the study with the complete key adjacent and the major cellular mechanisms of histogenesis below (b) Whole-tissue transcriptome top 3 principal components (PCs); color code from (a) (viewable in 3D, Supplementary Video 1). *n* = 156 bio replicates. (c) Hierarchical clustering of differentially expressed genes, heat map (bottom) for normalized log_2_(FPKM) values; 2 bio-replicates per tissue. Thy, thymus; Spl, spleen; Lvr, liver; Hrt, heart; Mus, skeletal muscle; Bld, bladder; Adr, adrenal gland; Kdn, kidney; Lng, lung; Stm, stomach; Int, intestine; Lmb, limb; Fac, craniofacial prominence; Fb, forebrain; Mb, midbrain; Hb, hindbrain; Nt, Neural tube. Right panel: Normalized loadings of each gene for the top 5 PCs. Bottom panel: normalized scores of the top 5 PCs (same sample order as clustergram). Gene ontology (GO) terms for the top 100 positive-loading and top 100 negative-loading genes abbreviated as key words (bottom right). (d) Integrating single-cell organogenesis data from whole mouse embryos (Cao et al. 2019) with the whole-tissue transcriptome clustering (c). *Y*-axis, genes are ordered as in (c); *x*-axis, 38 cell types from Cao et al. (2019). A point in the diagram indicates expression of a Cao et al. marker gene with horizontal jittering. Boxes highlight specific cell types and gene clusters of interest (see text).

#### 2) Specification and differentiation

The most numerous and diverse temporal drivers reflect cell differentiation pathways. For example, PC5 is prominent in differentiating skeletal muscle systems of limbs and face (*p* = 3e-12), with the high-PC5-loading Cluster 2 containing genes turned on as myogenesis progresses (Fig. 1c and Extended Data Fig. 2–2). Neuronal and glial differentiation in CNS tissues are highlighted in PC1 (*p* =2e-22), prominently marking genes of Cluster 34 (Extended Data Fig. 2–34), that are further parsed from single cell marker distributions by cell sub-type (Fig. 1d).

#### 3) Inter-tissue cell migration

Migratory cell populations, either invading or exiting, are well known to be important for the development of many tissues, as detailed further below using scRNA-seq data of the limb. At whole-tissue resolution, examples include a blood component (e.g. PC2 *p* =3e-35) that emerges prominently in the haematopoietic tissue of origin (liver) and then in other tissues (Fig. 1c and Cluster 10 in Extended Data Fig. 2-10), while genes marking maturing B-cells ^15–18^ in Cluster 10 appear in liver, and then in tissues with developing lymphatics (Extended Data Fig. 3b).

### Additional data structure

Much additional dynamic and biological structure is summarized schematically at the major cluster level and is annotated further for individual clusters and sub-clusters (Extended Data Fig. 2). The anterior/posterior (AP) spatial axis was evident from its enrichment in six of the top twenty PCs of different *Hox* cluster members expressed according to their known positional codes (Supplementary Data 1 and 2, and expression clusters 19 and 25 in Extended Data Fig. 2–19 and 2–25). Reanalyzing specific gene groups of interest, such as transcription factors (Extended Data Fig. 5a-e), or applying specialty algorithms can provide additional insights such as anti-correlations of miRNAs with predicted polyA RNA targets ^19^. To evaluate additional meta-data feature effects on transcriptome structure, we applied canonical correlation analysis ^20,21^ (CCA, see Methods), which identified dissection-based batch effects and sex-specific expression that may be pertinent to some future data uses (e.g. differential amounts of maternal blood; thymic contamination of some lung and heart samples; sex-biased samples from embryos of different sex. Extended Data Fig. 1l and 6, Supplementary Data 3).

### Global transcription factor DNA motif topology

RNA co-expression patterns revealed by clustering (Fig.1c; Extended Data Fig. 2) are caused in part by transcriptional co-regulation. Elevated frequencies of TF recognition sequence motifs in promoters of co-expressed genes can computationally link specific TFs or TF families to their likely target genes and regulatory elements. We tested the proximal promoters (500 bp) of all genes in each expression cluster, (numbered according to the expression cluster origin in Fig. 1c) for enrichment of all known consensus TF binding motifs (718 motifs) (see Methods). A bipartite graph was constructed to identify local and global relationships between the resulting combinatoric motif-codes and their source expression clusters (Fig. 2). First, the resulting 307 significantly enriched motifs displayed expected local relationships: fetal liver Cluster 10 is characterized by haematopoetic (GATA1/2, Runx1, Bcl11a) and hepatic (SMAD1, PPARG, NR1H2) markers, the highly specific Rfx factor family marks its cilium cluster (Cluster 28); and the E2F family is prominent in the previously discussed cell cycle-themed Cluster 21 (Extended Data Fig. 2; Supplementary Data 5).

The graph topology also shows binary and higher-degree motif code-sharing (gray shaded nodes) that selectively connect specific expression cluster promoter nodes from Fig. 1c with each other, suggesting that they jointly use identical or paralogous TFs. At a high level, the prominent separation of neurogenesis (Cluster 34) from haematopoiesis (Cluster 10) first observed in the transcriptome emerged independently for the motif codes, with only two shared motifs between them, whereas many other clusters share numerous motifs with each of them and with each other. The ubiquitous expression cluster had the strongest and most numerous motif enrichments in the entire transcriptome, with extensive Ets and Cre family representation (Fig. 2b and Extended Data Fig. 8e), whose enrichment and occupancy have been previously associated with human housekeeping genes ^22,23^. Finally, the most extensive code-sharing among expression clusters was with CNS neuronal Cluster 34 which connects with many other clusters of diverse tissue origins and functional themes (Figs. 1c; 2b). A plausible explanation for this CNS-centric sharing pattern is that many involved TFs (and/or their paralogs) were recruited during evolution to new uses that support increasing mammalian neuronal diversity.

**Figure 2:**
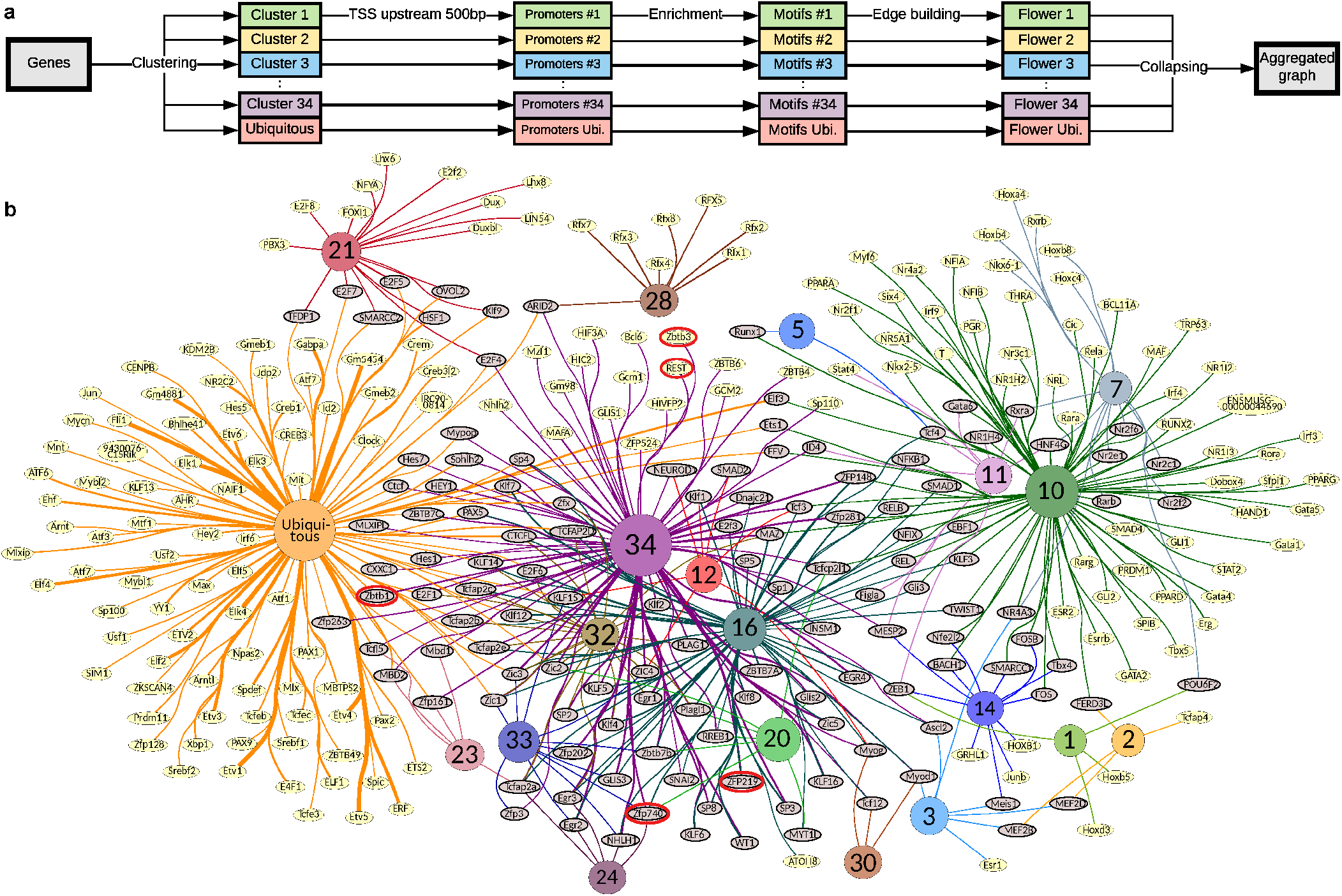
Promoter motif codes for dynamic expression clusters of Figure 1. (a) Flowchart for motif enrichment analysis. (b) A computed graph summary of unique and shared TF recognition sequence motifs. TF motifs and their respective source gene cluster identities (colored and numbered per clusters in Fig. 1c) are shown as graph nodes. Edges connect the motif node and the expression cluster node(s) in which it is enriched, with edge thickness indicating significance (−*log*_10_(*p*-value)). Motifs enriched in more than one cluster (gray) versus yellow for unique enrichment. The size of each source expression cluster node is proportionate to the scaled number of genes in the corresponding cluster.

### Cluster-specific regulatory mechanisms

The transcriptome structure and corresponding promoter motif resource provide entry points to identify cluster-specific regulatory mechanisms. For example, integrating our transcriptome and global epigenomic maps across matched samples showed that the up-regulated brain Cluster 34 has strong repressive histone mark density (H3K27me3) at early developmental times that declines with rising RNA expression (Extended Data Fig. 7e,f). Subsequent global quantification of developmental differentials in H3K27me3 promoter signal relative to RNA output across all clusters found that brain clusters 30, 32 and 34 stand out as candidates for a H3K27me3-mediated de-repression mechanism, despite similarly rising RNA trajectories in many other clusters (Extended Data Fig. 7a,d). Our prior motif enrichment analysis showed that the neuronal repressor REST/NRSF’s motif is specifically and strongly enriched in cluster 34 promoters (Fig. 2b). The putative targets of REST/NRSF, inferred from an independent ChIP-seq study ^24^, are also specifically enriched in Cluster 34 (Extended Data Fig. 7b); the RNA expression of REST/NRSF decreases in brain tissues over time (Extended Data Fig. 7c); and REST/NRSF-occupied promoters ^24^ show even greater H3K27me3 signal enrichment at early times (Extended Data Fig. 7f), all of which is consistent with a significant role in CNS-focused de-repression. This in vivo brain result agrees with a prior cell culture-based neural progenitor study ^25^, but it contrasts with an embryonic stem cell study reporting no H3K27me3 enrichment at REST/NRSF locations ^26^. Beyond REST/NRSF, other candidate repressors whose motifs are enriched in clusters 34 and/or 32, also exhibit expression trajectories that diminish as development progresses (e.g. Zfp219, Zbtb1, Zbtb3, Zfp740; red oval outlines, Fig. 2b) while additional presumptive C2H2 zinc finger transcriptional repressors whose recognition motifs are unknown, are concentrated in the CNS-enriched expression Cluster 33 (Extended Data Fig. 5e) with overall downward expression trajectories (Extended Data Fig. 2–33). Our working model is that they provide additional targeting diversity and specificity for the pervasive H3K27me3-mediated repression/depression process of the developing brain. This will become testable as their individual binding targets and derived motifs are determined. In a separate analysis, we examined the large ubiquitous cluster and found evidence suggesting that a post-transcriptional mechanism is playing a substantial role in setting divergent levels of expression within the ubiquitous cluster (Extended Data Fig. 8).

### Histogenesis at single cell resolution

From E10 to E15.5 the developing forelimb progresses from a simple limb bud composed mainly of undifferentiated mesoderm to a highly patterned structure with distinct skeletal, muscular, vascular, haematopoietic and dermal tissue systems (Fig. 3a). Two scRNA-seq datatypes were collected; each spans the same time points as the parent bulk tissue study (Fig. 3): 1) 920 cells from the C1 platform, sequenced to relatively high depth (∼1M reads/cell), which achieved sensitive RNA detection rates, and full-length transcript coverage comparability with the bulk data (Extended Data Fig. 1c–k); and 2) ∼90,000 cells from the 10x Genomics 3’ end-tag platform which expanded cell type discovery (Extended Data Fig. 1c-e). We detected in the high-resolution data 15,931 protein coding and 938 lncRNAs, of which 91% and 71% respectively overlapped with the limb whole tissue time-course (Extended Data Fig. 1b), while the 10x data captured 81% and 36%. Comparing these data with published whole embryo scRNA-seq ^11^, showed overlap of cell type relationships (Extended Data Fig. 9b) and genes, with 15,314 protein coding genes in common plus 2,230 and 637 genes novel to the whole embryo and the forelimb, respectively. This is consistent with greater cellular breadth in the whole embryo study versus deeper cellular and molecular coverage in the more focused forelimb study (Extended data Fig. 1 c-e).

**Figure 3:**
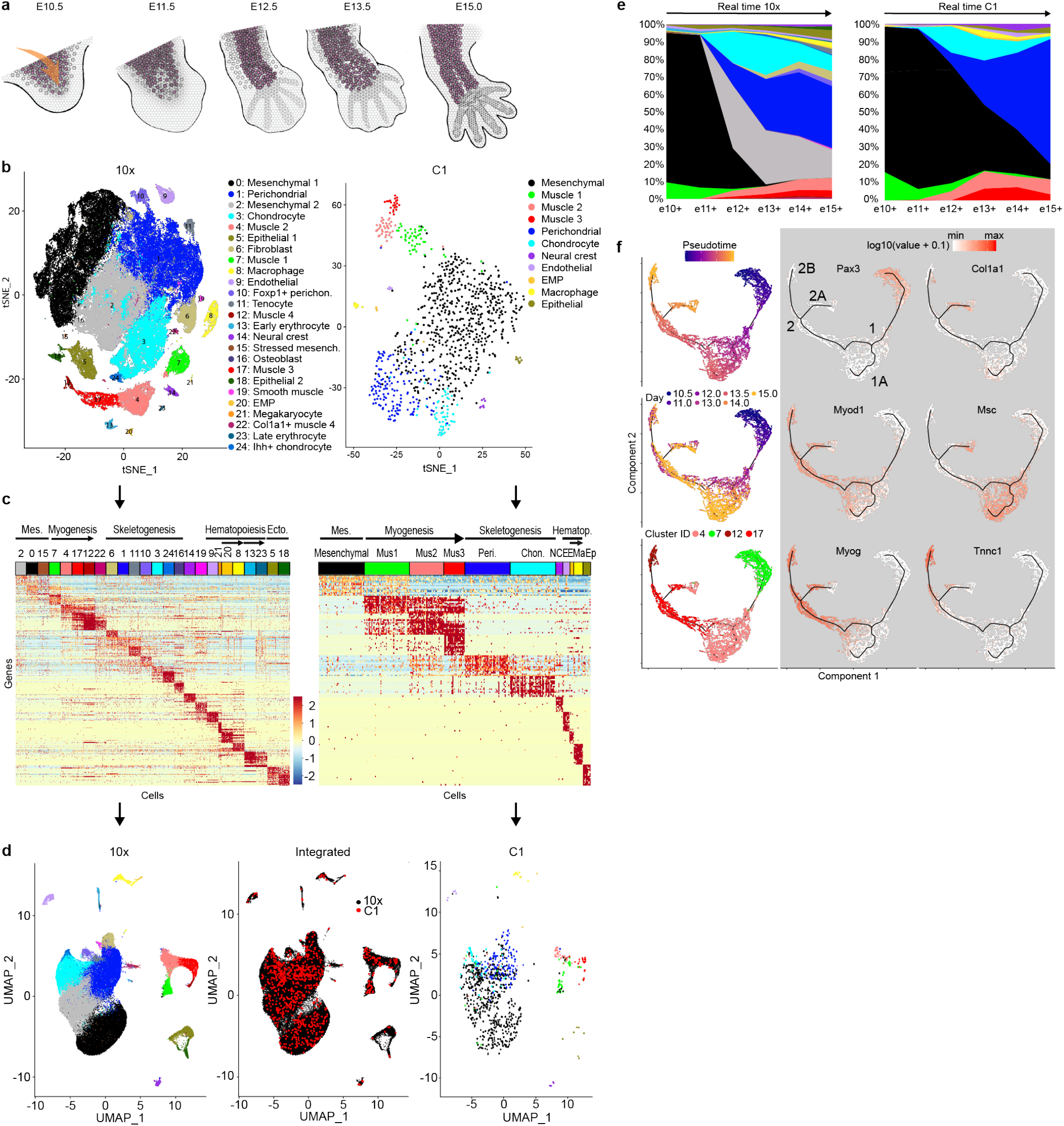
Single cell analyses of forelimb histogenesis. (a) Limb development schematic. Arrow indicates immigrating lineages. (b) 2D *t*-SNE of cell clusters, of 10x (left, *n* = 90,637 cells) and Cl data (right, *n* = 920 cells). Colors indicate provisional cell identities per Supplementary Note 1. (c) Cell cluster marker genes (top 15 per cluster), down-sampled for display to 100 cells per cluster for 10x and 30 cells for Cl. (d) Integrated visualization of 10x (left) and Cl (right) single cells on a 2D UMAP plane, separately or jointly projected (center panel) (see text and Methods). (e) Cell type composition plotted as a time series. The color code corresponds to cell clusters in (b). (f) Monocle lineage inference model for skeletal myogenesis. Pseudotime, developmental time and cell type (left); informative marker gene expression mapped on the right; *n* = 7,668 muscle cells.

### Resident and immigrating cell types

Clustering the most differential genes across all cells identified major progenitor and differentiating cell types and showed similarity relationships between them (Fig. 3b-d; Extended Data Fig. 9; Materials and Methods). Provisional cell identity assignments were based on GO enrichment analysis together with support from the developmental literature for previously reported “marker” genes (Supplementary Note 1, Supplementary Tables 1 and 2; references and discussion of marker gene limitations therein; Fig. 3b,c). Major cell types in both studies included resident limb-bud mesenchyme and its chondrogenic and osteogenic derivatives, plus independently immigrating lineages that give rise to myogenic, monocyte/macrophage, endothelial or neural crest derivatives. 10x data additionally provided evidence for 14 more cell types or states. When projected into the whole-tissue transcriptome and compared with similarly projected whole-embryo scRNA-seq, this deeper and more focused limb sampling showed lineage subdivisions and sharpening of some types compared with the whole embryo (e.g. myocytes, connective progenitors, limb mesenchyme; Extended Data Fig. 9b).

### Developmental progression and lineage inference

Whole transcriptome *t*-SNE and UMAP and phylogenetic clustering analyses segregated cell types (Fig. 3b-d; Extended Data Fig. 9c-e) whose trajectories through time were then mapped (Fig. 3e). The extent of under-representation of large multinucleated myotubes, together with other possible disaggregation, differential cell capture and survival, and stochastic sampling artifacts, were assessed relative to unperturbed whole-limb RNA data using CIBERSORT ^27^ to produce an adjusted tissue proportion model (Extended Data Fig. 9f,g).

Computed UMAP and Monocle lineage models (Fig. 3d,f and Extended Data Fig. 9c) were consistent with classical and modern tracing studies and inferences from genetic knockouts, while also identifying new relationships and associated regulators. In the myogenic system early progenitors require the Pax3 TF to migrate into the limb bud from adjacent axial somites ^28–30^, and *Pax3* is indeed the strongest differential gene defining the Muscle1 cell cluster (Wilcoxon rank sum test: 3.7-fold enrichment in 10x data and 16.7-fold in C1 from both data-types), which mapped to the earliest Monocle pseudo-time group (Fig. 3f). The stages in the progression and inferred relationships among stages are defined by overall correlation patterns among differential genes (Extended Data Fig. 9a,b), while specific marker genes from the myogenesis literature provided biological interpretation and hypothesis generation (Fig. 3f; Extended Data Fig. 9d).

The Monocle myogenic lineage model showed two branch points, with the first in both real time and pseudotime producing a branch 1A consistent with an important known population of muscle stem cells that later give rise to the regenerative cells of adult muscle. They are marked by the genetically essential *Pax7* regulator (Extended Data Fig. 9d), and its direct target *Msc* (Fig. 3f), which represses myocyte differentiation ^31,32^. Branch point 2 leads, on one hand, to expected mature myocytes marked by *Tnnc* expression (branch 2B), while branch 2A was unexpected. It models a cell population expressing signatures of interstitial muscle fibroblasts (IMFs) ^33^, such as *Col1a1* and *Osr1/2*, in addition to classic myogenic markers including *MyoD1* (Fig. 3f; Extended Data Fig. 9h). We confirmed that individual cells in the developing forelimb co-immunostain for muscle and IMF marker proteins (Extended Data Fig. 9i). This phenotype resembles the small and somewhat mysterious 10x Cluster 22, and a second Monocle model incorporating Cluster 22 cells supports that interpretation (Extended Data Fig 9h). These observations and models, considered in light of prior studies showing that adult tissue IMFs have latent myogenic capacity ^34–36^, raise questions about the developmental origin (from resident mesenchyme or *Pax3*^+^ precursors); adult fate (whether to become an adult IMF and whether to maintain myogenic potential); and the biological significance of these cells. More broadly, we confirmed and extended prior microarray results on populations of FACS enriched muscle precursor cells ^37,38^ and recent scRNA-seq of *Pax3*-GFP selected cells ^39^. Our Monocle models share some basic characteristics with the one originally constructed by Trapnell and colleagues ^40^, although the systems and models also reflect substantial differences between adult human muscle regeneration in vitro and fetal murine myogenesis in vivo.

In haematopoiesis, we identified both EMP and macrophage at early stages, aided by their exceptionally robust sets of marker genes (Supplemental Note 1; Extended Data Fig. 9e), which is consistent with limb macrophage developing from limb-resident EMPs (Extended Data Fig. 9c) in situ. Finally, the skeletogenic system and its resident progenitors are the largest limb component throughout the time course. Condensation, expansion and differentiation into cartilage and bone is the primary fate of the resident limb mesenchyme, ^41–43^ represented here by UMAP and Monocle models (Extended Data Fig. 9c) that focus on putative chondrocytes and fibroblast/perichondrial cells that form two dominant branches from the mesenchyme. The structure detected is much less clearly partitioned and ordered than myogenesis, and a more refined single-cell resolved model of skeletogenesis will likely require deeper and more focused cell sampling coupled with spatial genomics methods to capture additional anatomical clues. ^44–47^

### *Trans*-acting cell-type TF networks and their *cis*-acting candidate enhancers

Each cell type cluster has a substantial set of differentially expressed transcription factors (Supplementary Data 4). In the myogenic lineage, these differential TFs were expressed in three modes with different regulatory and lineage inference implications (Fig. 4a,b; Extended Data Fig. 9d,j): 1) sharply stage-restricted Boolean patterns separate cell stages from each other, including the well-known causal transcription regulators *Pax3*, *Pax7*, *Msc*, and *Myog*; plus newly added ones (e.g. *Sp5* and *Sox8*); 2) lineage-restricted uniformly expressed regulators that define the lineage (*Pitx2* and *Six1*); and 3) multi-stage TFs with graded expression levels like *MyoD1* and *Pitx3*, whose expression joins two or more stages together, while nevertheless discriminating stages quantitatively (Fig. 4b). Some regulators, including TFs widely understood to function only at later stages in the lineage, were detectably and precociously expressed at low levels, but only in the more sensitive C1 data (Fig. 4a,b). For example, low level *MyoD1* in *Pax3*-expressing cells is detected ahead of MyoD1’s well-known myoblast- and myocyte-stage functions ^48^. This suggests that the locus is already open, and companion ENCODE DHS and browser inspection of histone mark data at E10.5 show distal and promoter proximal sites that support this idea (Extended Data Fig 11f).

Known protein and genetic interactions were used to organize all cell-type differential TFs into their respective interaction networks (myogenic lineage Fig. 4c; all other cell type clusters Extended Data Fig. 10), showing that pan-lineage and graded factors extensively switch interacting partners across stages of the myogenic lineage progression. The inference leverage provided by the low-level graded-pattern genes was platform sensitive, with the higher sensitivity of the C1 data detecting anticipatory (and also trailing) expression in sequential stages that had escaped detection in our 10x data (Fig. 4a).

### *Cis*-acting cell-type regulatory elements: Decomposing whole-tissue epigenomics

The companion ENCODE whole tissue histone modification, chromatin accessibility and DNA methylation data provide rich biochemical signatures from which candidate regulatory elements can be computationally inferred at the whole-tissue level ^2,13,14^, although they lack cell type resolution. To parse genomic elements that are selectively active in a given cell type or state (Fig. 4d), we first defined the boundaries of biochemically active sequence elements using the companion limb DNase peak calls. We then applied IDEAS ^13,14^ to learn and summarize epigenomic features over fixed genomic segment bins, and extracted those DNase peaks that overlap with active and bivalent IDEAs bins (the bivalents include both active signals from minor cell types diluted by cells with alternative signatures, as well as poised elements) (http://personal.psu.edu/yzz2/IDEAS/). We assigned these active elements to cell types based on the cell type specificity of their associated genes from scRNA-seq. Summing the active and bivalent signatures, among 2,208 cell-type and lineage-specific genes, 2,018 (91.4%) had at least one affiliated active or poised element among the total collection of 22,230 (Supplementary Data 6). Individual loci with multiple candidate elements, plus supporting IDEAS state tracks, developmental DHS and RNA expression patterns are shown for biologically important chondrogenic, myogenic and macrophage examples (Fig. 4e; Extended data Fig. 11b,c). Based on our overall element recovery and on prior limb tissue reconstruction results (Fig. 3e; Extended Data Fig. 9f,g), we estimate that the whole limb epigenomic data have the sensitivity to identify validated cell type enhancers for cells comprising less than 5% of the starting population.

We evaluated all elements in our collection that overlap with the independently derived VISTA transgenic mouse database of regulatory elements. For the overlapping set 63% were validated VISTA enhancers (https://enhancer.lbl.gov/) distributed across our major cell types ^2,49^ (Fig. 4f). We did not expect all IDEAS overlapping elements to have scored positively in the VISTA assay paradigm for reasons summarized in the accompanying paper ^2^ and because of VISTA’s narrower developmental time-window (E11.5– E12.5 versus the IDEAS limb input E10.5–E15). The spatiotemporal domains in VISTA typically included limb *LacZ* transgene staining, but additionally showed staining elsewhere in the embryo. This is expected, as our major cell types are all represented elsewhere in the body and are not restricted to the limb. Conversely, there are some spatially patterned limb elements in VISTA (e.g. Mm1505 and Mm1492; Extended Data Fig. 11b) that do not appear limited to a single cell type, and so are not in our cell-type differential collection. Compared with the mouse FANTOM enhancer and promoter sets, which were computed from CAGE data and covered a wide sampling of fetal tissues and cell types ^10^ (http://fantom.gsc.riken.jp/5/), the limb IDEAS set overlaps with 44% and 30% of all FANTOM promoters and enhancers, respectively. Of these, 14% of each (9,943 promoters and 2,147 enhancers) are specific to our cell-type collection. Another large group of ours (20,119 and 19,384 IDEAS cell-type enhancers and promoters) were not in the FANTOM collection, which is overall a smaller collection (Extended Data Fig. 11e).

**Figure 4.**
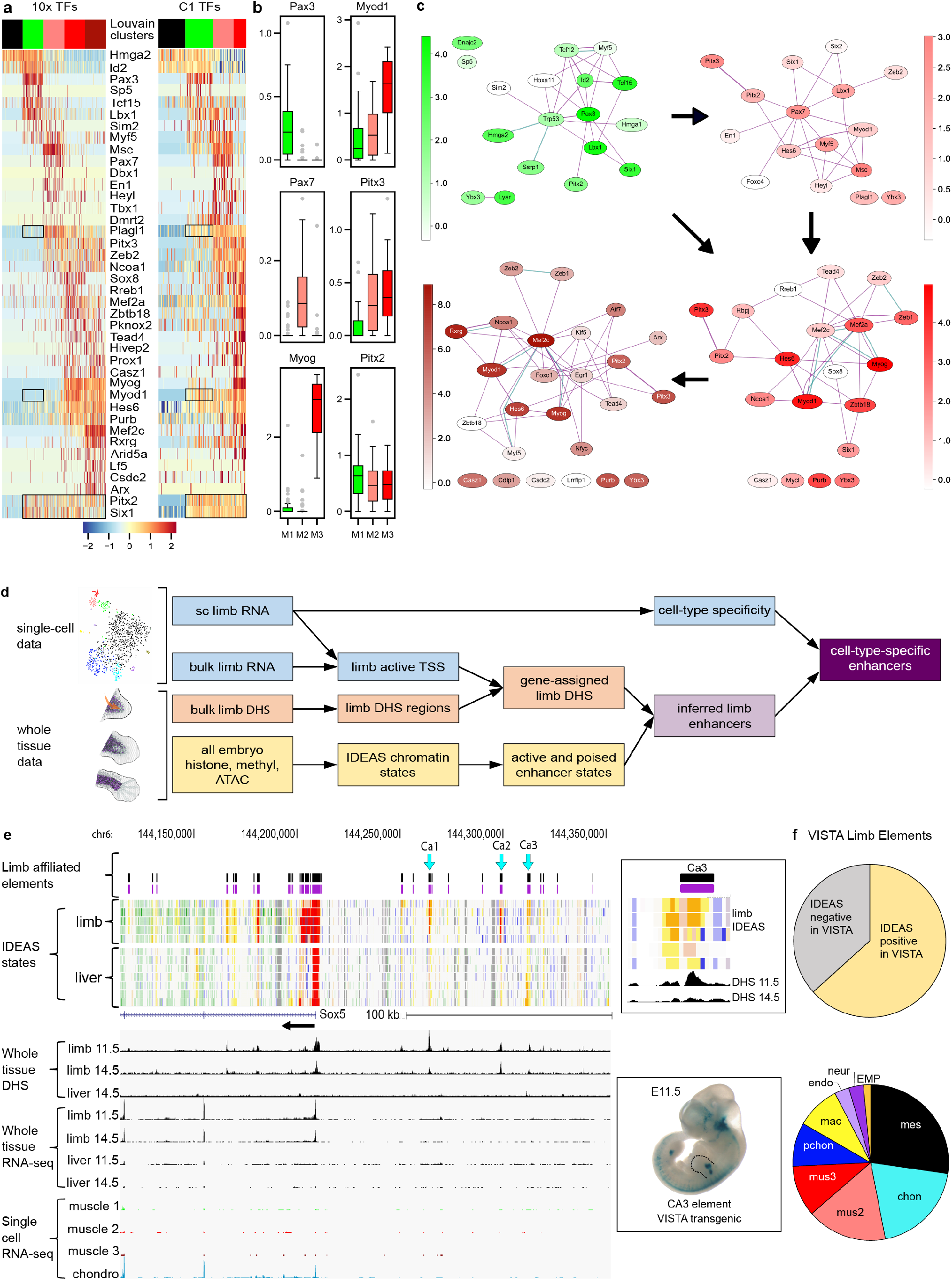
*Trans*-acting and *cis*-acting regulatory networks inferred for specific limb cell types. (a) Transcription factors enriched in limb mesenchyme and skeletal muscle lineage clusters from 10x or C1 data. Cells down-sampled for display per Fig. 3c and color-coded for cell-cluster identity per Fig 3b. Boxes highlight genes (*Myod1*, *Plagl1*) with early stage expression detected in C1 but not 10x data versus pan-lineage markers (*Six1*, *Pitx2*) detected similarly in both. (b) Box plots of Boolean, graded, and pan-lineage pattern TFs; *n* = 23 Muscle 3 cells; *n* = 38 Muscle 2 cells; *n* = 54 Muscle 1 cells. Boxes are 25th–75th percentiles; median-centered; minima and maxima 1.5×interquartile. (c) STRING networks of skeletal muscle lineage for cell-type differential TFs from 10x data (methods); edges are colored by types of STRING evidence (cyan for database and magenta for experimental); nodes colored according to 10x RNA-seq levels; arrows indicate lineage transitions (see text). (d) Schematic for discovering cell-type enhancer and promoter elements using scRNA-seq and IDEAS chromatin state elements defined in whole tissue chromatin assays (see text, methods and Extended Data Fig. 11a). (e) Candidate upstream limb skeletal enhancers (Ca1-3) for *Sox5* with in vivo enhancer data from VISTA for a Ca3-containing segment at right (https://enhancer.lbl.gov/frnt_page_n.shtml). Computed IDEAS limb cell-type elements (purple track); IDEAS epigenomic segmentation tracks below with poised and active enhancer type (orange) and promoter type (red) states below. (f) Summary of IDEAS/scRNA-seq cell-type elements in the VISTA resource. Top: IDEAS limb elements in VISTA (*n* = 235/371, 63%). Bottom: VISTA-positive IDEAS elements by cell-type (*n* = 66 cell-type-specific elements).

Transcription factor binding motifs significantly enriched in cell-type IDEAS elements (Supplementary Data 5) of distal candidate enhancer elements (≥2Kb from the affiliated TSS) or promoters, were organized in computed graphs that show lineage-related cluster nodes joined by motif sharing across stages and related cell types (i.e. muscle clusters 4, 12, 17; haematopoetic clusters 8, 13, 20, 21 in Extended Data Fig. 11d). Neural crest stood out for its large number of distal motifs, including many Hox family members, likely reflecting the use by neural crest derivatives of positional signaling gradients in their specification and migration. We similarly extracted motif codes for genes whose expression is significantly depleted in a cell typespecific manner. Such genes were especially prominent in early haematopoetic cells, and their distal and promoter elements were strikingly enriched in repressor and Hox motifs. Although speculative, this suggests to us a regulatory logic in which cells traversing the entire embryo silence genes which, in other cell states or types, actively respond to positional signaling. Overall, it should be possible to model cell-type/state-preferential enhancers in this and other ways for other ENCODE tissues by adding a corresponding sc-RNA-seq dataset and integrating it with the IDEAS models or another epigenomic state model of choice.

## Conclusions

As developmental biologists might expect, the fetal transcriptome reflects known molecular and cellular mechanisms of histogenesis at multiple levels of organization. Across all tissues, we found a relatively simple and universal temporal RNA signature, attributable to changing stromal and erythroid cell components. And within individual tissues and organs, distinctive signatures emerge from shifting proportions of constituent cell lineages, each progressing toward cytodifferentiation and a final spatial distribution. These sensitive whole-tissue data, uniformly processed and encompassing 17 tissues, define a dynamic transcriptome whose derived co-expression modules and corresponding DNA motif topology comprise an initial regulatory code map for mouse fetal development that can be further interpreted by single-cell data.

An advantage of the ENCODE fetal transcriptome compared to prior conceptually similar efforts is the opportunity to integrate companion epigenome and microRNA resources. ^2,19,50,51^ As an example, single cell RNA-seq decomposed whole-tissue chromatin state models (IDEAS) built from ENCODE epigenomic data, to extract cis-regulatory enhancer/promoter element collections for major limb cell types, together with corresponding trans-acting TF identity networks and their respective motif codes. This could be generalized to other tissues by introducing appropriate scRNA-seq, and further strengthened by integrating scATAC-seq and more sophisticated algorithms ^52–54^.

Both tissue-level and single-cell datasets are available through the ENCODE portal for browser viewing or large-scale computing. They cover a substantial window of mouse fetal development, but the earliest steps for most organs were not included, owing mainly to limitations imposed by available tissue amounts. Recent progress in low-input and single-cell methods for both transcriptome and epigenome mapping open the door to filling these gaps.

## Materials and Methods

### Bulk RNA-seq from mouse embryo tissues

Pulverized pooled mouse embryo tissue replicates from timepoints E10.5, E11.5, E12.5, E13.5 E15.5 and E16.5 were received from the Ren lab which supplied these tissues for the entire mouse development project ^47^. E14.5 and P0 tissues were dissected from single animals at Caltech. Replicate tissue samples were lysed and extracted using the Ambion mirVana protocol (AM1560). Residual genomic DNA was removed using the Ambion Turbo DNA-free kit (AM1907). Total RNA was quantified with Qubit and RIN values were collected with the BioAnalyzer Pico RNA kit (5067–1513). The median RIN value was 9.7 (CV=4.4%). Each cDNA library was built using 10 ng total RNA spiked with ERCC spikes (AM4456740) diluted 1:5,000 in UltraPure H_2_O (InVitrogen 10977023) containing carrier tRNA (AM7119) at 100 ng/*μ*L, RNAse inhibitor (Clontech 2313A) at 1 units/*μ*L and DTT (Promega P1171) at 1 mM. cDNA was reverse-transcribed and amplified according to the protocol in the SMARTer UltraLow RNA kit for Illumina (634935) using Clontech SMARTScribe reverse transcriptase (639536), and TSO, dT priming and amplification primers from the Smart-seq2 protocol ^5^. The first-strand product was cleaned up on Ampure XP beads, and then amplified using the Clontech Advantage 2 PCR kit (639207) with 13 PCR cycles and an extension time of 12 minutes. After a second round of Ampure XP cleanup, the amplified cDNA was quantified on Qubit and the size distribution was checked with the HS DNA BioAnalyzer kit (5067-4626). cDNA libraries were then tagmented using the Illumina/Nextera DNA prep kit (FC 121-1030) with index tags from Illumina (FC 121-1031), cleaned up with Ampure XP beads, quantified on Qubit and sized with the Agilent HS DNA kit. Libraries were sequenced on the Illumina HiSeq 2500 as 100 bp single-end reads to 30M aligned reads depth. Inclusion for ENCODE submission required replicate concordance scores by Spearman correlation of FPKM values ≥ 9.0.

### Single-cell transcriptome measurements using the Fluidigm C1 and 10x Genomics v2

One pair of embryonic forelimbs from a single mouse was used at each timepoint (E10.5, E11.0, E11.5, E12.0 and E13.0, E13.5, E14.0, E15.0. After dissection from the carcass, limbs were incubated in a 50 *μ*L droplet of a 10% collagenase solution (Worthington LS004202) for 5 minutes at 37 °C. The limbs were then visualized under a dissecting scope and the ectoderm was removed manually with a pair of #5 Dumont forceps. The mesenchymal core of the limb bud was then transferred to a 200-*μ*L droplet of Accumax (AM105), and the dish was reincubated for 15 minutes at room temperature. The cells were then manually triturated once with a P200 tip to suspend them, and pipetted into 500 *μ*L of DMEM + 10% FBS. Limb cells were spun at 500 *g* for 5 minutes at 4 °C, resuspended in 500 *μ*Ls fresh DMEM + 10% FBS, and passed over a 20 micron mesh (Miltenyi 130-101-812). They were then counted and diluted in DMEM + 10% FBS to achieve a final concentration of 250,000 cells/mL. 12 *μ*Ls of this suspension was added to 8 *μ*Ls of Fluidigm Cell Suspension Reagent for loading on the Fluidigm IFC (10-17 micron size). Cells were then visually inventoried for doublets and empty chambers, and returned to the C1 for lysis, reverse transcription and amplification using the SMART-Seq v4 protocol. Lysis buffer: 8.6 *μ*L water, 1 *μ*L C1 loading buffer, 2.4 *μ*L Smartseq2 oligo dT primer (10mM), 2.4 *μ*L Clontech 10mM dNTPs, 2 *μ*L ERCC spikes (AM4456740) (diluted 1:40,000 in UltraPure H_2_O (InVitrogen 10977023) containing carrier tRNA (AM7119) at 200 pg/*μ*L, RNAse inhibitor (Clontech 2313A) at 1 units/*μ*L and DTT (Promega P1171) at 1 mM), 0.5 *μ*L 100mM DTT, 2.6 *μ*L Clontech single-cell reaction buffer. Reverse transcription reaction: 5.6 *μ*L Clontech 10x transcription buffer, 0.6 *μ*L C1 loading buffer, 5.6 *μ*L Smart-seq2 TSO (10mM), 0.4. *μ*L Clontech RNAse inhibitor, 2.8 *μ*L Clontech SMARTScribe. PCR reaction: 4.4 *μ*L H_2_O, 4.5 *μ*L C1 loading buffer, 75.2 *μ*L Clontech SeqAmp buffer, 3 *μ*L Smart-seq2 amplification primers (10 mM) and 2.9 *μ*L Clontech SeqAmp polymerase.

Amplified cDNA samples were diluted in 10 *μ*L of C1 DNA dilution reagent, and a 1 *μ*L aliquot of each was quantified on Qubit. 11 samples from the IFC were selected for BioAnalyzer sizing based on yield and chamber occupancy. An aliquot of the cDNA libraries was diluted to 0.1–0.3 ng/*μ*L using C1 Harvest reagent, and the libraries were then tagmented using the Nextera XT DNA sample prep kit (FC 131-1096) and Nextera XT indices (FC 131-1002). After tagmentation and amplification, libraries were pooled, cleaned up twice with Ampure XP beads (0.9× volume), quantified on Qubit and sized on the BioAnalyzer using the HS DNA kit. The libraries were then sequenced as 50-bp single reads to a depth of about 1M aligned reads on the Illumina Hi-Seq 2500.

10x Genomics single-cell libraries were prepared from the single-cell suspensions described above, targeting 10,000 cells per library, exactly as described in the manufacturer’s protocol. They were sequenced as 150 bp paired end libraries, to a depth of 400M reads each on the Illumina Hi-Seq 4000.

### Read mapping and quantification

All the whole-tissue RNA-seq and C1 single-cell RNA-seq data were processed through the standard ENCODE pipeline (https://www.encodeproject.org/pipelines/ENCPL002LSE/), which uses STAR to align raw reads against mm10 genome with spikes and quantifies transcript abundances using RSEM which provides FPKM, TPM and count values. Downstream analyses were mainly done using Matlab scripts (https://github.com/brianpenghe/Matlab-genomics). 10x single-cell RNA-seq data were processed using CellRanger with a compatible GTF annotation and default parameters.

### Whole-tissue RNA-seq PCA, CCA and Hierarchical clustering

tRNA genes and genes covered by fewer than 10 reads in all tissues were removed. Principal Component Analysis (PCA) was performed over the *log*_2_-transformed FPKM values (with 0.1 added as pseudo-counts) to unmask relatively lowly expressed transcripts in order to accommodate high sensitivity of whole-tissue RNA-seq assays. *Z*-scores of eigenvalues from PCA were used to visualize “PC scores”, while eigenvector coefficients from PCA were used to visualize “PC loadings”. Genes with top highest positive values and lowest negative values were used to interpret biological meanings for each PC.

Canonical Components Analysis (CCA) was performed on the top 20 PCs and Boolean variables for tissue identities, stages, gender and dissection metadata. Standardized canonical variables scores were visualized using the heatmap in Extended Data Fig. 6c, while *z*-scores of sample canonical coefficients were visualized using the heatmap in Extended Data Fig. 6b and d. Canonical-correlation gene loading coefficients could be calculated by multiplying the PC-gene loading coefficient matrix (from PCA) and canonical-correlation PC loading coefficients (from CCA). Genes with top highest positive values and lowest negative values could be used to interpret biological meanings for each CC (Supplementary Data 3).

The dynamic genes were defined as those with at least 10-fold difference in FPKM values between the most and least abundant RNA samples; genes with less than 10-fold difference were defined as flat, or ubiquitous. Dynamic genes and ubiquitous genes were categorized into different classes (protein-coding etc.) based on gene types annotated by GENCODE M4. One-way and two-way hierarchical clustering were done using Pearson correlation coefficient and average linkage for the dynamic genes. Clusters were defined by traversing from the root of the tree towards the leaves, and splitting out clades with different dominant tissues and GO terms, recognized manually, until no more major clusters could be split out. Clades with at least 30 nodes were defined as major clusters. In order to test the robustness of the results, we did an independent analysis with the forebrain, hindbrain and neural tube removed to decrease CNS representation, using the same methodology. Another independent analysis was performed using TPM values for all the tissues, using the same methodology. The main conclusions were largely the same.

### Whole-tissue RNA-seq transcription factor analysis

Transcription factor expression vectors were used to generate *t*-SNE and clustering maps using the same settings as the whole-transcriptome analysis. Transcription factor families were compared against cluster identities. The hypergeometric test was performed to assess enrichment.

### Embryo sex inference

For the samples that were made from single embryos, we inferred their sex by comparing gene expression levels of *Xist* (a female marker) and *Ddx3y* (a male marker). Embryos that expressed *Xist* only are female while those that express *Ddx3y* only are male. Mixed embryo pools had both genes detected.

### Ubiquitous gene analysis

Among the genes defined ubiquitous by the whole-tissue RNA-seq analysis, those with *log*_2_(FPKM + 0.1) values no higher than 2 were removed. The 3000 genes with smallest sample variance were equally assigned into high, medium and low groups based on their average FPKM values.

GRO-seq and Bru-seq reads were mapped and quantified using the ENCODE standard pipeline for computational consistency. Average 3’ UTR lengths for each gene were extracted from the GENCODE M4 annotation. The *log*_2_(FPKM + 0.1) values and *log*_2_(3’ UTR length) were used for comparisons and linear regressions.

### Histone modification analysis

Histone modification ChIP-seq data were processed using the ENCODE ChIP-seq pipeline (https://www.encodeproject.org/pipelines/ENCPL220NBH/), and *log*_2_ fold change for ChIP-seq samples over input controls were calculated and plotted using Deeptools2.4.1 (https://github.com/fidelram/deepTools/tree/2.4.1). To summarize the fold decrease of histone modification signals in a specific sample among a specific cluster of genes, a 4-kb window enclosing the TSS at the center is used and average *log*_2_ fold change against input samples were calculated and visualized using a 3D heated barplot. The fold decrease is the difference between the fold changes of the earliest and latest timepoint. Rest target overlap p-value is calculated based on the hypergeometric test using the iQNP Rest ChIP-seq target list from Mukherjee et al, 2016..

### Gene Ontology Analysis

FuncAssociate 3.0 (http://llama.mshri.on.ca/funcassociate/) was used at its default settings for term calling.

### C1 Single-cell RNA-seq clustering and *t*-SNE visualization

Spike and tRNA gene FPKM values were removed to rescale FPKM values. Libraries with no cell or more than one cell in their corresponding C1 chambers spotted by microscope were removed. Libraries from the same C1 Fluidigm chip that had systemic 3’ coverage bias were all removed. Cells with fewer than 100,000 reads mapped to the transcriptome or fewer than 4000 genes above 10 FPKM cutoff were removed. Genes that were expressed in less than 5 cells (0.5%), or at lower than 10 FPKM in all cells, or that were covered by fewer than 100 mapped reads in all cells were filtered out. We then used *log*_2_-transformed (FPKM + 1) values for the following analyses. The genes were ranked based on their dispersion scores (defined by sample variance over sample mean). The top 1,500 genes were selected, from which non-coding genes and mitochondria genes were filtered out, leaving 1,269 genes. *t*–SNE projection was done based on these genes, using the top 30 PCs and 30 as perplexity parameter (default for Laurens van der Matten’s original MATLAB script) ^55^. Two-way hierarchical clustering was then performed on the *log*_2_-transformed FPKM values using complete linkage with Spearman rank correlation coefficient to cluster the cells. Cell types were annotated manually.

### 10x Single-cell RNA-seq clustering and *t*-SNE visualization

UMI counts from CellRanger were filtered first, where cells with fewer than 1000 genes detected and genes detected in less than 0.1% cells were removed. Within each cell, counts were divided by the sum and multiplied by 10,000, added by 1, and *log*-transformed. Top 4,000 high-dispersion genes were identified. To remove noise (https://github.com/brianpenghe/python-genomics), we first performed a hierarchical clustering for these genes and then extracted genes that fell in “tight” clusters (those with more than 2 members after cutting the dendrogram at 0.8 distance), removing a large number of sporadic genes which had high dispersion scores but were barely co-expressed with other genes. These genes were used in place of “highly-variable genes” for the Seurat pipeline. Using the Seurat pipeline, cells with more than 20% mitochondria reads or more than 8000 genes detected were removed. Genes were scaled and regressed against the number of UMI per cell and mitochondria percentage. The resulting matrix, guided by the aforementioned feature genes, was used to perform PCA. Jackstraw was then performed using Seurat’s default settings, resulting in 42 significant PCs. These PCs were in turn used for Louvain cell clustering and tSNE visualization. Clusters 3,4,5,6,8,12 and 13 were further re-clustered using the same method, yielding clusters 17-24.

### Marker gene identification for C1 and 10x single-cell RNA-seq data

Marker genes (Supplementary Table 4) were calculated using Seurat’s `FindMarkers()` for both C1 and 10x singlecell data with `min.pct` = 0.25 and its default Wilcoxon rank sum test with `min.diff.pct` set to be 0.2 or 0.4. For marker visualization, each cell type was down-sampled to at most 100 cells for 10x data and at most 30 cells for C1 data. `min.diff.pct` was set to be 0.2 and top 15 markers for each cell type were visualized.

### Comparing C1 and 10x cell types

Two methods were used to compare cell type annotations for C1 and 10x data. Based on Seurat3’s “Label Transfer” method, transfer anchors were calculated from 10x data and were used to predict cell types for C1 data. Independently, the scaled 10x data matrix was used to train a multinomial logistic regression model using `scikit-learn` package. The trained model was used to predict cell types for C1 data.

### Integrating C1 and 10x data for UMAP visualization

Seurat3 was used to calculate integration anchors and to integrate the two different types of datasets. The joint set was scaled and visualized on UMAP based on an arbitrary top 50 PCs.

### Lineage trajectory analyses

Prior to lineage inference, doublets were removed using a Scrublet-based ^56,57^ subclustering scheme. Monocle3 alpha (2.99.3) was then used for trajectory analysis of the 10x data that contain a large number of cells. The function `plot_pc_variance_explained()` was used to select significant PCs above the knee cutoff. UMAP visualization and SimplePPT method were applied. The root node for each lineage tree was defined as the node that connects to the largest number of the cells from the earliest developmental timepoint (E10.5).

### Differential transcription factor analysis

Transcription factors recorded at TFDB (http://bioinfo.life.hust.edu.cn/AnimalTFDB/) were selected from marker genes derived at 0.2 cutoff (described above), to infer evidence-based interaction networks using STRING ^58^ (https://string-db.org/). A Python interface for STRING was used to query the database directly and render the resulting graph using Graphviz ^59^. Edges of type “database” and “experimental” were used, filtered to meet a confidence value of greater than 0.400. Nodes were colored using normalized values obtained from SCANPY ^60^. The graph was laid-out using layout software included with the Graphviz package. The algorithm used was SFDP. The complete code base as well as Docker and Singularity container recipes can be accessed on the GitHub repository: https://github.com/hamrhein/mouse_embryo.

### IDEAS states

The IDEAS epigenetic states on the ENCODE3 mouse developmental data were generated by the IDEAS software ^13,14^ using 10 epigenetic marks: H3K27ac, H3K27me3, H3K36me3, H3K4me1, H3K4me2, H3K4me3, H3K9ac, H3K9me3, ATAC-Seq and DNAse methylation data. We first converted the raw data in each sample to −*log*_10_*p*-values using a Negative Binomial model. The mean and variance parameters of the model for each sample were calculated using the bottom 99% of the data. We then adjusted the mean parameters at each genomic position from the input data to account for local genomic variations. Specifically, we downloaded the input data for each tissue (see list of data sets), and we calculated rolling means per genomic position using a 20-kb window centered at the position, for both signals and the input. The ratio between the two means at each position is multiplied to the overall mean estimate of the sample, and we normalized the ratios across the genome to have mean 1. We treated the −*log*_10_*p*-value as input data for IDEAS, capped at 16, and we ran the program in its default setting. The output from IDEAS is a set of genome tracks to display in the genome browser, where each epigenetic state is assigned a color as a weighted mixture of colors preassigned by the program to each epigenetic mark. The IDEAS segmentation can be accessed by the Hub link at http://bx.psu.edu/~yuzhang/me66/hub_me66n_org.txt.

### Cell type and lineage-specific marker genes identification and cCRE assignment

Genes exclusively expressed in only one cell type or lineage were regarded as marker genes for this series of analysis. Using the high-resolution C1 Fluidigm data, marker genes at 0.2 or 0.4 cutoff were cross-intersected to derive exclusively expressed markers of cell types or groups of related cell types (Muscle 1+Muscle 2, Muscle 2+Muscle 3, Muscle1-3, Chondrocyte+Perichondrium, EMP+Macrophage etc.). Candidate *cis*-regulatory elements (cCREs) were defined by merging all the DHS peaks called by the ENCODE HOTSPOT2 pipeline. These merged regions were assigned to closest transcription start sites of genes that are expressed (FPKM higher than 0.1 in at least one bulk limb tissue, or detected in more than 4 cells in single-cell limb data). These merged regions were then compared against IDEAS chromatin states generated from ENCODE3 mouse developmental time course data (see below). Only the peaks that overlapped with active (state 14, 19, 20, 21, 23, 24, 25, 27, 28, 30–32), poised (8 and 13) or bivalent (26 and 29) IDEAS states were regarded as “IDEAS active DHS” (cCREs). Finally, these cCREs assigned to the aforementioned marker genes’ TSS’s were regarded as cell type or lineage-specific cCREs. Based on the distance between each cCRE and its assigned gene, cCREs were further divided into three categories: proximal (the distance is no greater than 200bp in any direction), middle (the distance is longer than 200 bp no greater than 2,000 bp in any direction) and distal (the distance is longer than 2,000bp in any direction).

### Motif analysis

For whole-tissue RNA-seq promoter motif analysis, the upstream 500 bp sequences of each co-expression cluster were extracted and pooled. For limb cell type-associated gene promoter analysis, the upstream 500 bp sequences of each cell type’s marker genes (derived from 10x data using Seurat, `min.diff.pct` = 0.4) were extracted and pooled. For limb cell type-associated cCRE analysis, the DNA sequences of proximal, middle, or distal cCREs for each cell types marker genes were extracted and pooled. These sequence pools were used for motif discovery.

A detailed flowchart can be found in Extended Data Figure 11.

The analysis of transcription factor recognition motifs was carried out using version 4.11.2 of the MEME-SUITE ^61^. Motifs annotated in the CIS-BP database ^62^ (http://cisbp.ccbr.utoronto.ca/) were used to evaluate motif enrichment in the sequence pools mentioned above; enrichment was scored by the AME program in the MEME-SUITE ^63^. The analysis was carried out twice based on UCSC mm10 refFlat and GENCODE M4 separately and only motifs with corrected *p*-values smaller than 0.01 in both analyses were called significant.

### Comparing whole-tissue RNA-seq and single-cell RNA-seq

10x single-cell data (without log transformation or Gaussian scaling) and the aforementioned 10x feature genes were used as input for CIBERSORT ^27^ (https://cibersort.stanford.edu/) to compare against whole-limb RNA-seq data (without log transformation or Gaussian scaling). To compare cell type-associated gene signatures against ENCODE whole-tissue RNA-seq clusters, cell type-associated marker genes were acquired from the article by Cao et al. 2019 (Table S4 for gene names and Table S3 for cell type names) and filtered (*p* < 0.05 and *q* < 0.05). These signature genes (noting that CIBERSORT is highly sensitive to the choice of input gene set) were mapped to the ordered heatmap of the bulk-tissue clustergram (Fig. 1d). For better visualization, we jittered individual dots, to create a re-purposed swarm plot to show distribution of the locations (instead of quantities) of signature genes for each cell.

### Immunocytochemical detection in tissue sections

Staged embryos were fixed in 4% PFA in PBS, cryoprotected with 30% sucrose in PBS, and frozen in OCT on dry ice. 10-micron cryosections were blocked using the mouse on mouse blocking reagent from Vector (cat. # MKB-2213), and then stained with antibodies for Osr1 (mouse monoclonal Santa Cruz cat. # 376545 at 1:40) and myogenin (Abcam RabMab cat. # ab124800 at 1:40). Secondary detection was done with InVitrogen donkey anti-rabbit Alexa 594 cat. # A21207, and InVitrogen goat anti-mouse Alexa 488 cat. # A11029, both at 1:300 dilutions. Sections were first screened on a Zeiss Axio Observer Z.1 and then imaged for deconvolution microscopy using a Leica DMI6000, with a 63X oil immersion lens, and Huygens Professional deconvolution software from SVI.

## Supporting information

Supplementary Table 1 - clusters

Supplementary Table 2 - PCA

Supplementary Table 3 - CCA

Supplementary Table 4 - Marker genes

Supplementary Table 5 - Motifs

Supplementary Table 6 - IDEAS-based elements

## Acknowledgments

We wish to thank Mr. Gordon Ace Dan for scientific illustration of limb development; Sean Upchurch and Sreeram Balasubramanian for data handling; Zhiping Weng and Arjan van der Velde for providing consolidated datasets to YZ; Bing Ren for leadership of the ENCODE mouse working group, Sarah A Teichmann, Lior Pachter, Cole Trapnell and Matt Thomson for discussions, and Hongbo Zhang and Krzysztof Polaski for discussion and advice on computing. We also thank Igor Antoshechkin at the Caltech Jacobs Genetics and Genomics Laboratory for sequencing the Illumina libraries, Sisi Chen and Jeff Park of the Single-Cell Profiling and Engineering Center at Caltech for building 10x Genomics libraries, Andres Collazo at the Beckman Institute Imaging Center for IF imaging work, and Eun Hee Shim and Rebekah Loving for supporting immunocytochemistry.

## Author Contributions

P.H. bionformatics, data analysis, figures, wrote the paper; B.A.W. performed all bulk and single-cell RNA-seq experiments, data analysis, wrote the paper; G.K.M. DNA motif analysis, edited paper; D.T. performed sequencing analysis, data submission, figure generation, edited paper; H.A. Network visualization, figure generation; L.B., S.-T.G. IF experiments, imaging and analysis; I.P.-F. V.A. staged and dissected mouse embryos; L.A.P. mouse developmental matrix design, oversight, and VISTA resource; D.E.D. A.V. coordinated and supervised mouse dissection and staging; B.R. Mouse developmental matrix design and oversight of mouse ENCODE effort; R.C.H. IDEAS development and edited the paper; Y.Z. developed and implemented IDEAS; B.J.W. supervised the project, analyzed the data, wrote the paper.

## Correspondence and requests for materials should be addressed to

Prof. Barbara J. Wold

Division of Biology and Bioengineering

MC 156-29

California Institute of Technology

Pasadena, CA 91125

## Data Availability Statement

These data are part of the ENCODE Consortium mouse embryo project, which provides companion microRNA-seq, DNA methylation, histone mark ChIP-seq, and chromatin accessibility datasets for the sample matrix (https://www.encodeproject.org/woldlab).

## Code Availability Statement

Standard ENCODE RNA-seq pipeline (https://www.encodeproject.org/pipelines/ENCPL002LSE/), ENCODE ChIP-seq pipeline (https://www.encodeproject.org/pipelines/ENCPL220NBH/), all Matlab scripts (https://github.com/brianpenghe/Matlab-genomics).

10x single-cell RNA-seq data were processed using Cell-Ranger with a compatible GTF annotation and default parameters.

Deeptools2.4.1 (https://github.com/fidelram/deepTools/tree/2.4.1)

FuncAssociate 3.0 (http://llama.mshri.on.ca/funcassociate/)

TFDB (http://bioinfo.life.hust.edu.cn/AnimalTFDB/)

Motifs annotated in the CIS-BP database (http://cisbp.ccbr.utoronto.ca/) STRING (https://string-db.org/).

The complete code base for STRING interaction graphs, as well as Docker and Singularity container recipes can be accessed on the GitHub repository: https://github.com/hamrhein/mouse_embryo.

The IDEAS segmentation can be accessed by the Hub link at http://bx.psu.edu/~yuzhang/me66/hub_me66n_org.txt

CIBERSORT (https://cibersort.stanford.edu/)

---

**Extended Data Figure 1:**
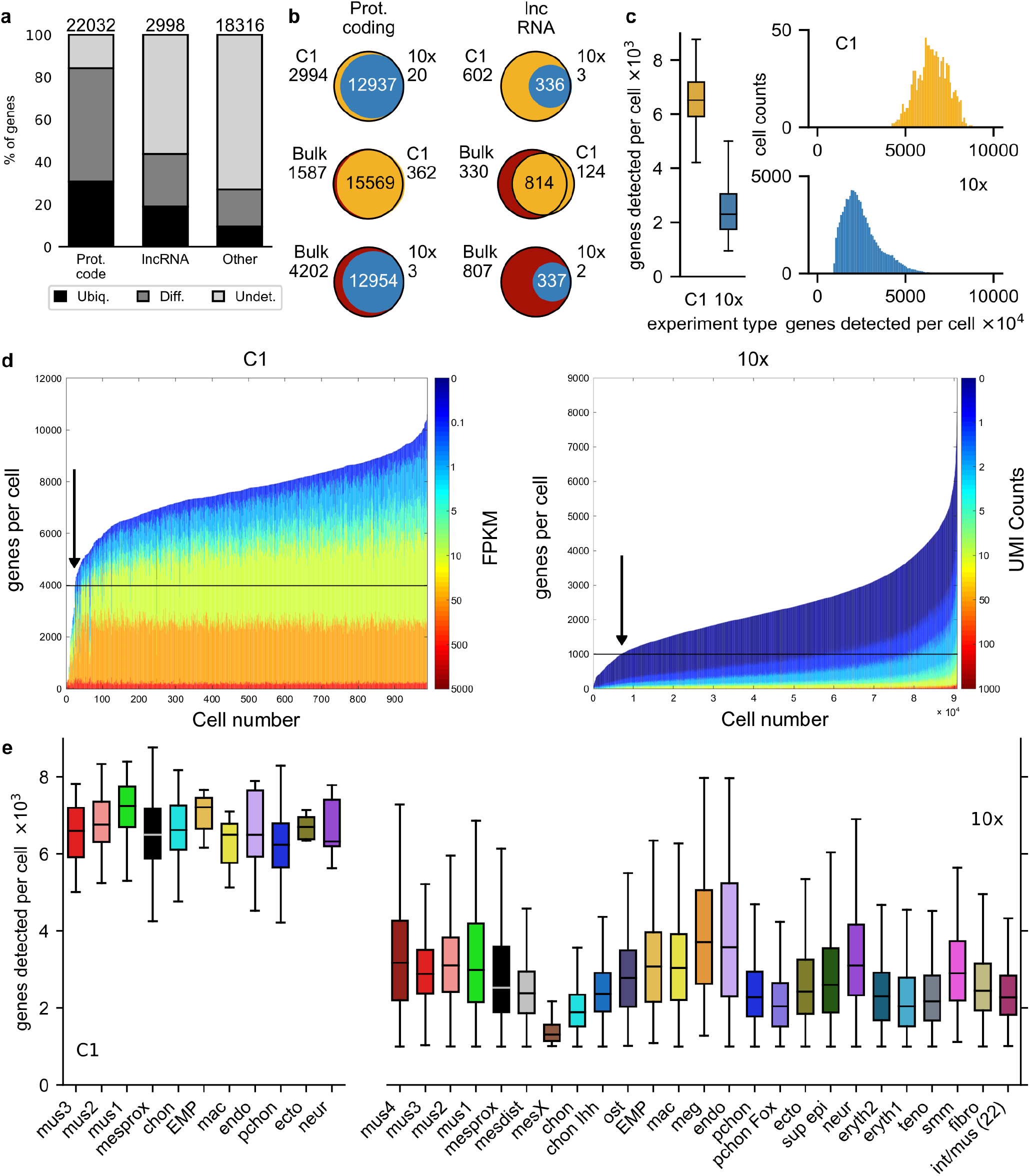
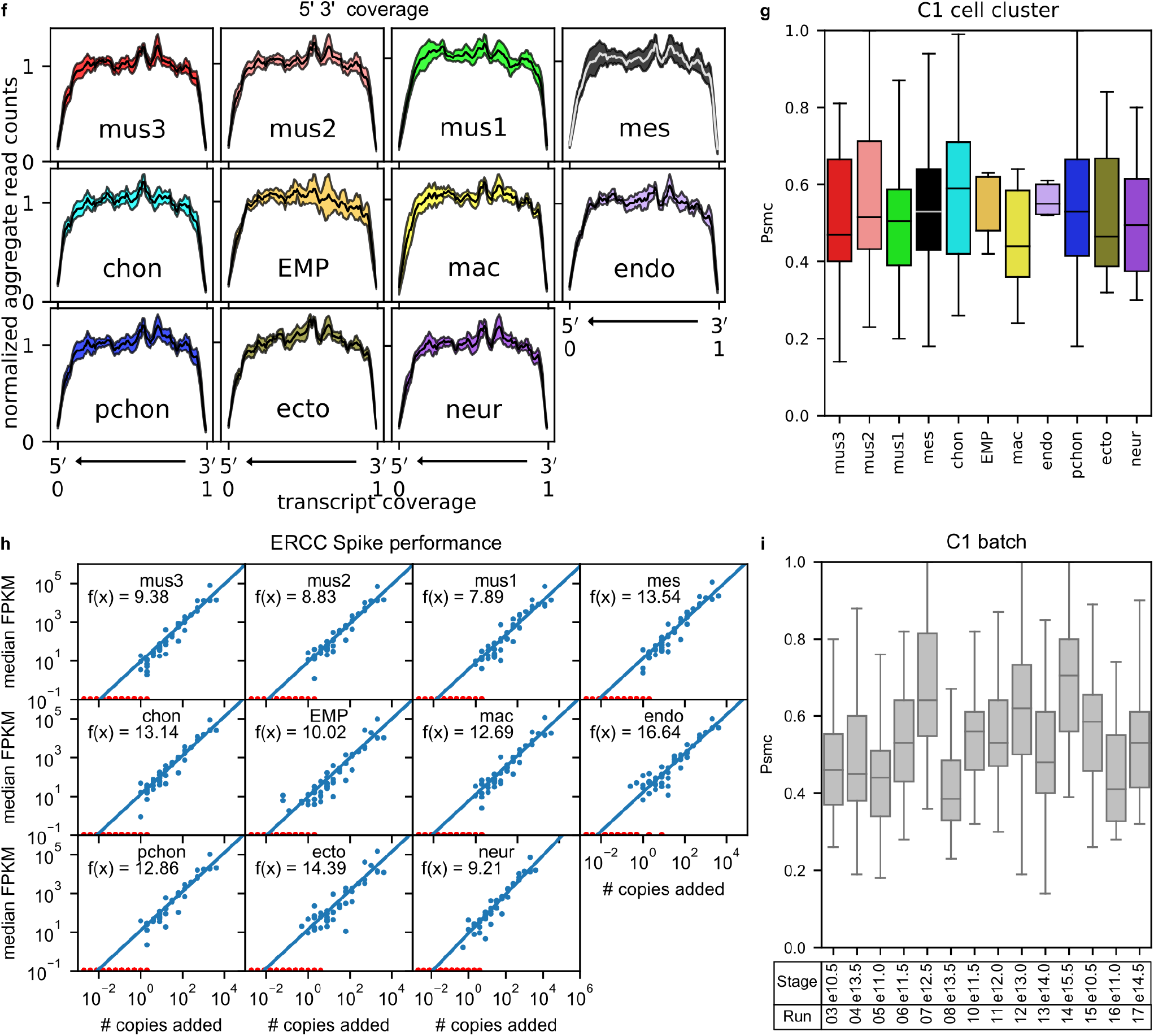
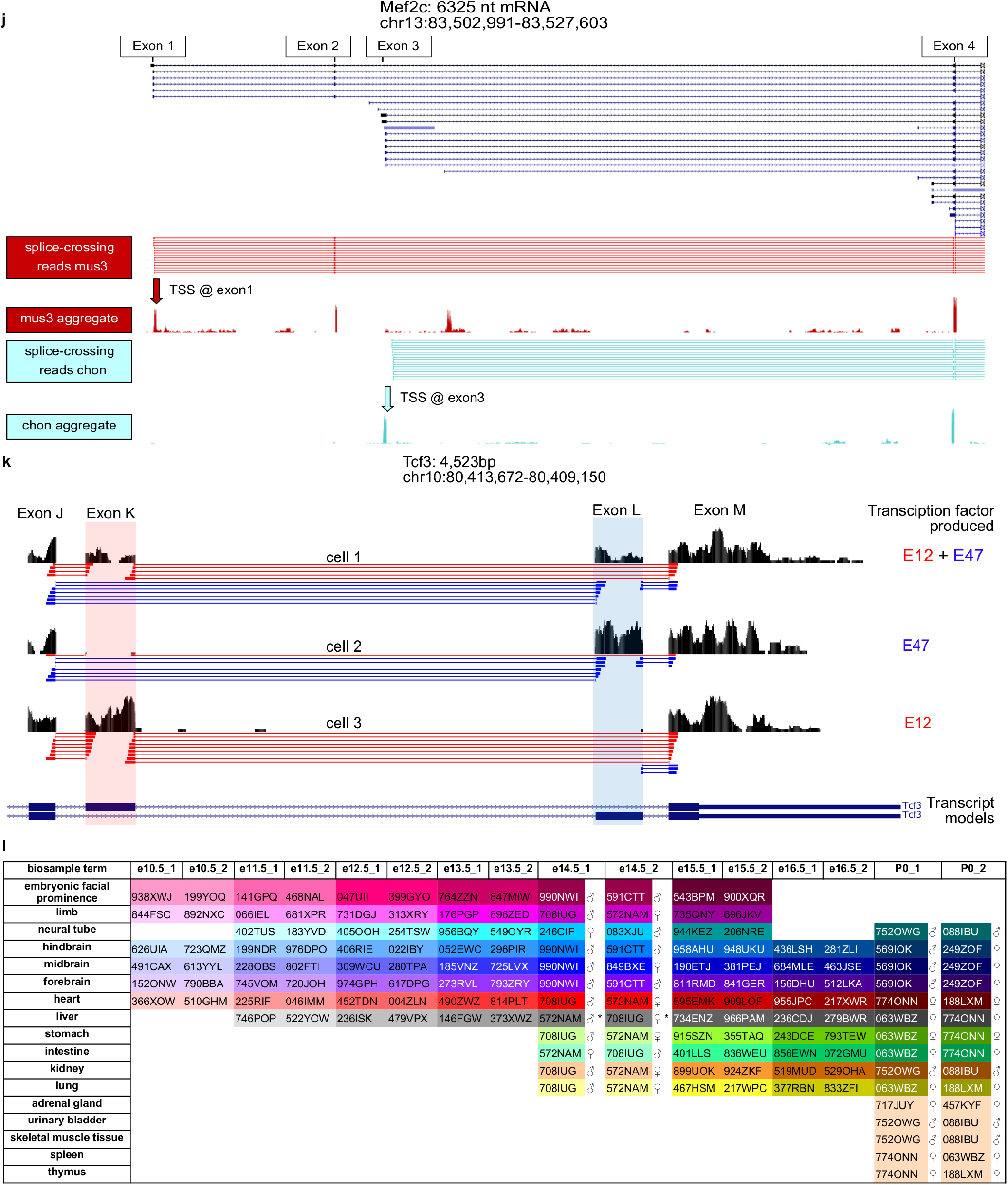
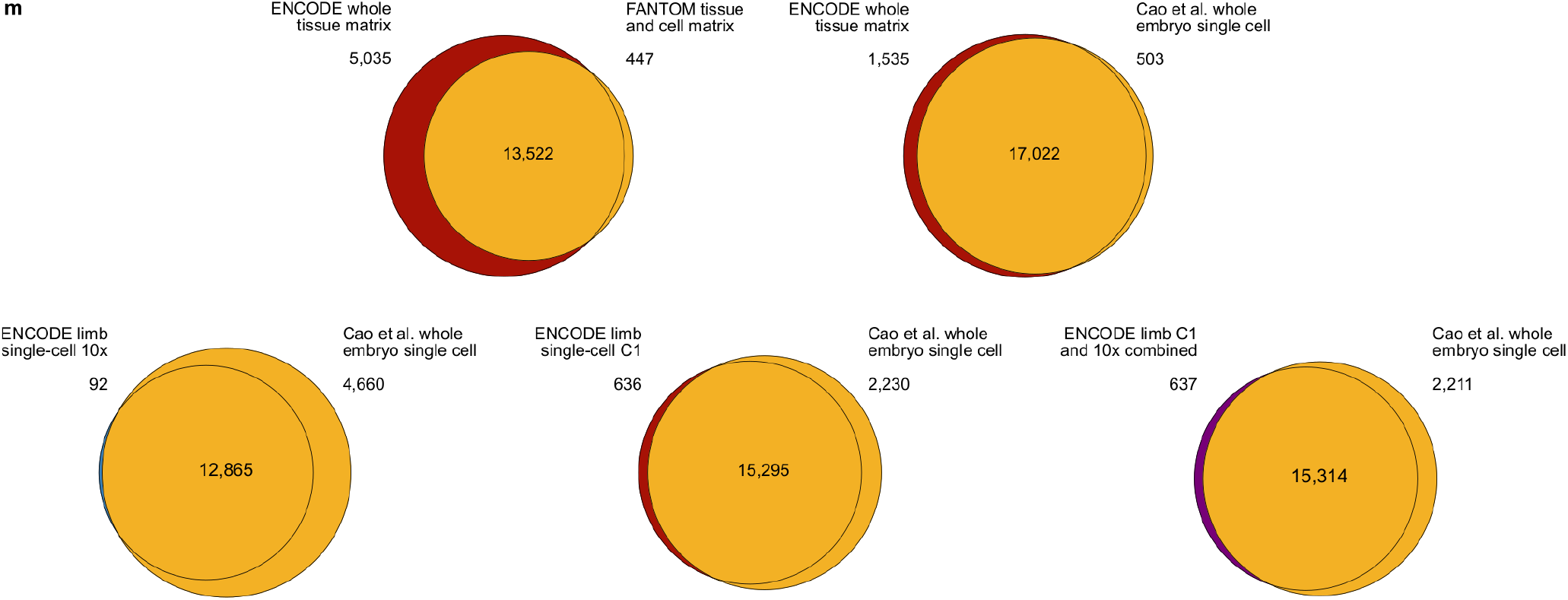
Quality metrics of bulk RNA-seq and single-cell RNA-seq. (a) Percentages of ubiquitous, differential and undetected genes in each of the three categories: Protein-coding genes (Prot. code), lncRNA (long intergenic noncoding RNA), and others. (b) Pairwise comparisons of detected protein-coding genes and lncRNAs among the three RNA-seq platforms. (c) Number of genes detected per cell by C1 and 10x platforms in boxplots (left panel), and full histogram distributions (right panel). *n* = 920 cells for C1; *n* = 90637 cells for 10x. (d) Genes per cell histogram distributions colored by abundance values. Cells are sorted in ascending order based on the number of genes detected per cell at the least stringent cutoff. Abundances are shown using the color scale on the right of the two plots. Arrows represent the “knee” cutoffs we picked for inclusion in the analysis (4000 genes/cell for C1; 1000 genes/cell for 10x). (e) Average numbers of genes detected among each cell type defined by the C1 platform (mus3, Muscle 3; mus2, Muscle 2; mus1; Muscle 1; mesprox, Mesenchymal; chon, chondrocyte; EMP, EMP; mac, Macrophage; endo, Endothelial; pchon, Perichondrial; sup epi, Epithelial; neur, Neural crest) and the 10x platform (mus4, Muscle 4; mus3, Muscle 3; mus2, Muscle 2; mus1, Muscle 1; mesprox, Mesenchymal 1; mesdist, Mesenchymal 2; mesX, Stressed mesenchymal; chon, Chondrocyte; chon Ihh, Ihh+ chondrocyte; ost, Osteoblast; EMP, EMP; mac, Macrophage; meg, Megakatyocyte; endo, Endothelial; pchon, Perichondrial; pchon Fox, Foxp1+ perichondrial; ecto, Epithelial 1; sup epi, Epithelial 2; neur, Neural crest; eryth2, Late erythrocyte; eryth1, Early erythrocyte; teno, Tenocyte; smm, Smooth muscle; fibro, Fibroblast; int/mus (22), Col1a1+ muscle 4). Boxes are 25th–75th percentiles; median centered; minimum and maximum 1.5×Interquartile. Left: *n* = 23 mus3 cells; *n* = 38 mus2 cells; *n* = 54 mus1 cells; *n* = 571 mesprox cells; *n* = 57 chon cells; *n* = 5 EMP cells; *n* = 10 mac cells; *n* = 7 endo cells; *n* = 139 pchon cells; *n* = 8 ecto cells; *n* = 8 neur cells. Right: *n* = 404 mus4 cells; *n* = 1,764 mus3 cells; *n* = 3,625 mus2 cells; *n* = 1,875 mus1 cells; *n* = 22,925 mesprox cells; *n* = 17,205 mesdist cells; *n* = 114 mesX cells; *n* = 10,536 chon cells; *n* = 494 chon Ihh cells; *n* = 86 ost cells; *n* = 238 EMP cells; *n* = 1,123 mac cells; *n* = 29 meg cells; *n* = 1,011 endo cells; *n* = 20,254 pchon cells; *n* = 912 pchon Fox cells; *n* = 2,719 ecto cells; *n* = 629 sup epi cells; *n* = 577 neur cells; *n* = 188 eryth2 cells; *n* = 425 eryth1 cells; *n* = 762 teno cells; *n* = 210 smm cells; *n* = 2,204 fibro cells; *n* = 328 int/mus(22) cells. (f) Transcript coverage from 5’ to 3’ (left to right on *x*-axis) in C1 single-cell libraries is uniform and consistent across the 11 different cell types. *Y*-axis is normalized, aggregate read counts. The center values are median values for each bin; the shading represents standard deviations for each bin. *n* = 23 mus3 cells; *n* = 38 mus2 cells; *n* = 54 mus1 cells; *n* = 571 mesprox cells; *n* = 57 chon cells; *n* = 5 EMP cells; *n* = 10 mac cells; *n* = 7 endo cells; *n* = 139 pchon cells; *n* = 8 ecto cells; *n* = 8 neur cells. (g) Probability of single-molecule capture (*p*_*smc*_) estimates for each of the 11 different C1 cell types. Boxes are defined by 25th and 75th percentiles; the center is median; minimum and maximum are 1.5×Interquartile. *n* = 23 mus3 cells; *n* = 38 mus2 cells; *n* = 54 mus1 cells; *n* = 571 mesprox cells; *n* = 57 chon cells; *n* = 5 EMP cells; *n* = 10 mac cells; *n* = 7 endo cells; *n* = 139 pchon cells; *n* = 8 ecto cells; *n* = 8 neur cells. (h) Estimated input (*x*-axis) and output (*y*-axis) amounts of ERCC spikes in each cell type. One cell is represented by one dot. The slopes of the fitted lines in *log* space have been labeled in each panel. (i) *P*_*smc*_ estimates for each C1 run. Error bars are standard error. Boxes are defined by 25th and 75th percentiles; the center is median; minimum and maximum are 1.5×Interquartile. *n* = 23 mus3 cells; *n* = 38 mus2 cells; *n* = 54 mus1 cells; *n* = 571 mesprox cells; *n* = 57 chon cells; *n* = 5 EMP cells; *n* = 10 mac cells; *n* = 7 endo cells; *n* = 139 pchon cells; *n* = 8 ecto cells; *n* = 8 neur cells. (j) Cell type specific TSS choice for *Mef2c* in the developing limb identified by short-read RNA-seq. UCSC genome browser tracks display Fluidigm C1 data from muscle3 (dark red) and chondrocyte (cyan) cells at *Mef2c* with Gencode VM20 gene and transcript models. Splice-crossing reads document exon1/2, 2/4 (red) and 3/4 junctions. Aggregate signal tracks for mus3 and chon show that the TSS at exon 1 is used in mus3, whereas chondrocytes select the TSS at exon 3. Median expressed level for *Mef2c* in muscle3 cells 53.4 FPKM; in chondrocytes 40.3. (k) Alternate splice choices in different single mesenchymal cells of the developing limb result in alternate forms of *Tcf3* (E12 and E47 bHLH TFs) with different DNA binding specificities. Individual splice-crossing reads are displayed beneath the read tracks for each of 3 separate exemplar cells. (l) Table representing all bulk RNA tissue/time samples in this study according to the color scheme in Figure 1, including ENCODE BioSample accession numbers. The individual embryo samples for E14.5 and P0 were characterized by sex-specific expression markers; embryo sex determinations are indicated. (m) Comparisons of whole tissue and single-cell transcriptome gene content with external whole tissue and single-cell resources ^10,11^. For all datasets, comparisons were restricted to only protein-coding genes that were detected.

**Extended Data Figure 2:**
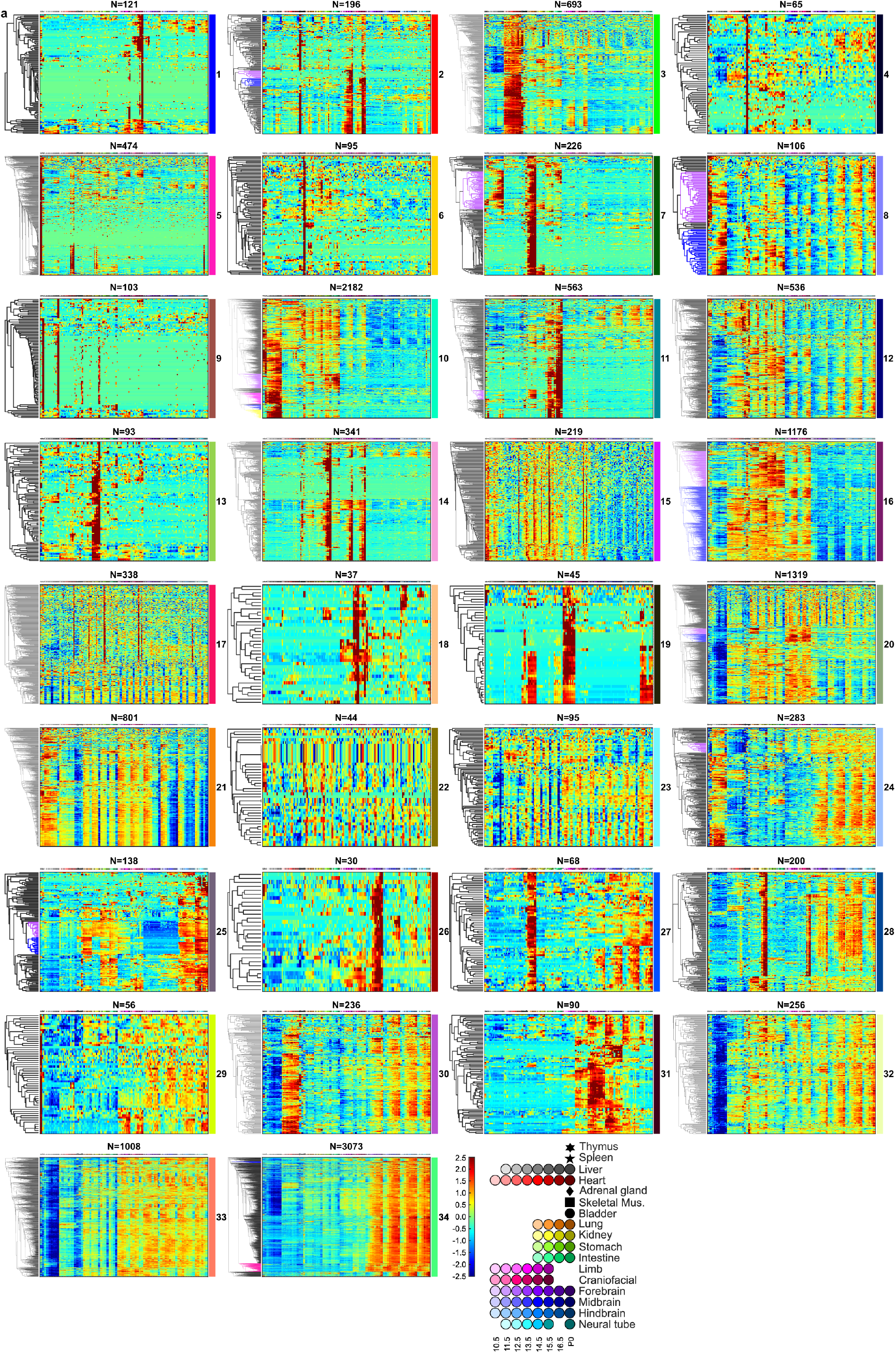
Individual clusters from hierarchical analysis. (a) Sample identities are labeled at the top with the code specified in Fig. 1. Cluster identities are labeled on the right, and the number of genes in each cluster (sample size *n*) is shown at the top of each panel. Normalized expression levels were mapped to the heatmap scale shown at the bottom right. Detailed descriptions of each cluster are given below. (b) Summary of expression cluster dynamics and dominant functional themes for clusters in (a). Rectangles represent major gene expression clusters from (a) with more than 30 members, labeled by the dominant features based on Gene Ontology, tissue specificity and gene class are labeled. Blue boxes indicate increase over time; pink decreases over time; green reflect relatively constant levels; lavender lacks coherent time course dynamics; yellow represent likely technical issues. The remainder are small clusters (≤30 genes), labeled as hexagons with the cluster size given. **Extended Data Fig. 2-1**. Cluster 1 from hierarchical clustering analysis of bulk RNA samples.

1. Cluster 1 has prominent increasing expression in limb and craniofacial prominence.
2. Over one third of the genes in this cluster are genes coding for keratin and keratin associated proteins. Top GO terms include “intermediate filament” (*p* = 3.7e-34) and “hair cycle” (*p* = 2.4e-7), pointing to development of skin and hair. **Extended Data Fig. 2-2**. Cluster 2 from hierarchical clustering analysis of bulk RNA samples.

1. Cluster 2 has prominent expression in skeletal muscle and an increasing trajectory in limb and craniofacial prominence.
2. Cluster 2 contains multiple muscle regulators like *Myod1* and *Myog*. Its top GO terms include “muscle system process” (*p* = 4.5e-18) and “contractile fiber part” (*p* = 9.9e-14). The increasing expression in limb and craniofacial prominence is likely due to differentiation of muscle precursors and to increasing relative muscle mass as a fraction of the total tissue.
3. In addition to the dominant muscle-limb-face feature, there are two clades with different patterns that illustrate the informational leverage that comes from a more pure P0 dissected tissue (here muscle).

a. The clade of 13 genes labeled in blue have increasing expression in limb and craniofacial prominence but not in the P0 pure skeletal muscle sample. Among the 13 genes, five (*Dcstamp*, *Mmp13*, *Bglap*, *Ifitm5* and *Ibsp*) are associated in prior work with osteogenesis.
b. The clade of 13 other genes labeled in purple is biased for limb alone, and not cranioface. It includes four major urinary protein (MUP) genes at low but detectable abundance. The mouse genome has 21 annotated Mup genes in a 2-Mbp cluster on Chromosome 4. Although none have human orthologs, members of the family have known functions in mouse chemical communication and nutrient metabolism. A recent study reported dramatic and unexpected upregulation of *Mup1* in mouse embryos when *Sox2*, a transcription factor regulating proximal bone formation in limbs, is mutated. This raises the possibility that the MUPs in this limb cluster play a role in limb development.
4. **Technical user note: Sporadic samples of adrenal gland, kidney, lung, stomach, hindbrain and neural tube from this mouse embryo bulk RNA ENCODE series show slight enrichments for genes from this cluster, implying variable minor tissue contamination during dissection. **Extended Data Fig. 2-3.** Cluster 3 from hierarchical clustering analysis of bulk RNA samples.

1. Most genes in this cluster have high and constant levels of expression in heart, and roughly half also have substantial expression in skeletal muscle-containing samples, which is expected due to contractile protein genes shared in both kinds of striated muscle.
2. GO terms are mainly about muscle, including “contractile fiber part” (*p* = 8.9e-47) and “regulation of heart contraction” (*p* = 4.3e-21)
3. *The clade in the upper half of the heatmap has narrow dark red bars, that indicate single replicate enrichment. This group of genes contains mostly pseudogenes (see also Cluster 15). **Extended Data Fig. 2-4**. Cluster 4 from hierarchical clustering analysis of bulk RNA samples.

1. Genes in Cluster 4 show differing degrees of bladder-specific expression, which may result from a bladder-specific cell type that has a unique transcriptome signature.
2. GO analysis produced no terms. Mouse bladder has not been extensively studied, and under-annotation may compromise the statistical power of GO in this case. **Extended Data Fig. 2-5.** Cluster 5 from hierarchical clustering analysis of bulk RNA samples.

1. Genes in Cluster 5 are very prominently expressed in thymus and most have minimal expression in other tissues.
2. Highly expressed genes also have positive signals in several non-thymus samples, with atypical irreproducibility between replicates. A candidate explanation is batch-specific contamination of thymus-proximate tissues with thymus during dissection. While this kind of contamination doesnt greatly alter global QC scores, it is readily detectable in this clustering analysis (see also CCA analysis).
3. GO analysis revealed enrichment in later stage maturing immune components, especially T-cell terms. Top terms include “immune system process” (*p* = 1.8e-18) and “regulation of T-cell activation” (*p* = 3.0e-13).
4. Roughly one quarter of the genes are T-cell receptor components (alpha chain, gamma chain and delta chain). Interestingly, the two recombinases *Rag1* and *Rag2* are also in this cluster, indicating a TCR VDJ theme for this cluster. **Extended Data Fig. 2-6.** Cluster 6 from hierarchical clustering analysis of bulk RNA samples.

1. The unifying theme of this cluster is high expression in the adrenal gland.
2. Top GO terms include “hormone biosynthetic process” (*p* = 1.5e-7) and “hormone metabolic process” (*p* = 7.3e-7). More specifically, *Cyp11b1*, *Cyp21a1* and *Cyp11b2* contribute to the term “mineralocorticoid biosynthetic process” (*p* = 2.8e-7). These cytochrome P450 genes are involved in biosynthesis of aldosterone which, unlike many other hormones, is produced only in the adrenal gland. However, these genes also have detectable expression signals in E15.5 and E16.5 samples of kidney. Their presence at E15.5 and E16.5 stages and absence in E14.5 and P0 may be due to contamination in E15.5 and E16.5 pooled samples, while E14.5 and P0 samples from individual embryos were more contamination-free. **Extended Data Fig. 2-7.** Cluster 7 from hierarchical clustering analysis of bulk RNA samples.

1. The central theme of this cluster is prominent expression in the developing kidney, where the RNA trajectories generally increase over time. Roughly 40% of these genes are also expressed in liver, again with increasing trajectories, plus some smaller subclades that are shared with gut or lung samples.
2. Top GO terms of this cluster include transporter-related categories such as “sodium ion transport” (*p* = 2.0e-14) and “anion transport” (*p* = 4.7e-9) and structural terms like “apical plasma membrane”. This cluster is dominated by genes responsible for transporter machinery and epithelial cell organization in the kidney. This cluster was also highlighted by CCA analysis. Further examination found that the increasing number of nephrons through time, and the up-regulation, in particular, of genes of the proximal tubule are explanatory.
3. The clade of 72 genes labeled in purple contains genes enriched in both liver and kidney. The top enriched GO terms for this group are for amino acid catabolic processes performed in both liver and kidney (“organic acid metabolic process” (*p* = 7.7e-12), “fatty acid metabolic process” (*p* = 1.2e-7) and “alpha-amino acid catabolic process” (*p* = 1.5e-6). 20 of these genes are enriched in kidney proximal tubule brush border cells while 7 are enriched in hepatocytes ^12^. **Extended Data Fig. 2-8.** Cluster 8 from hierarchical clustering analysis of bulk RNA samples.

1. The genes in Cluster 8 have increasing expression patterns in almost all tissues, although the kinetics of increase differ.
2. Top enriched GO terms include “inflammatory response” (*p* = 1.0e-6) and “extracellular exosome” (*p* = 1.5e-5).
3. There are two major clades. The clade labeled in purple is consistent with genes marking the immune system, whose levels are highest in thymus and spleen, but also include expression in the hematopoietic fetal liver. Subsets of these genes increase at later times in other tissues. GO analysis called terms including “regulation of T cell activation” (*p* = 8.8e-5) and “inflammatory response” (*p* = 2.2e-5). The second major clade, labeled in blue, is dominated by increasing expression in liver and gut tissues. Top GO terms included “extracellular exosome” (*p* = 3.3e-9). **Extended Data Fig. 2-9** Cluster 9 from hierarchical clustering analysis of bulk RNA samples.

1. The genes in Cluster 9 have highest enrichment by far in spleen and on the P0 liver, but not at earlier times. Moderate abundance is also seen in adrenal gland and lung. There is minimal but detectable expression in all other tissues at P0 but very little at all times before birth.
2. GO analysis did not yield significantly enriched terms, but more than half of the genes in this cluster are immunoglobulin components (kappa and lambda light chain variables, heavy chain variables and constant regions, consistent with B-cell maturation, appearing in liver, spleen and in lesser proportions in lung and other lymphatic-containing dissections. **Extended Data Fig. 2-10.** Cluster 10 from hierarchical clustering analysis of bulk RNA samples.

1. Over 60% of the genes in Cluster 10 are preferentially expressed in liver, much lower in CNS tissues and variously detected in other tissues. The RNA abundances mainly increase with time, but with differing kinetics.
2. Top GO terms of Cluster 10 include the immune system, such as “immune system process” (*p* = 4.8e-101) and “regulation of immune system process” (*p* = 2.0e-62). The additional prominence of many genes in the P0 thymus and/or spleen, along with other non-CNS tissues point to the lymphatic system.
3. In addition to the main immune theme, four clades with distinctions emerged. The one containing 267 genes labeled in purple are most enriched in liver, as well as stomach and intestine, increasing over time. Its top GO terms focus on lipids, including “lipid metabolic process” (*p* = 3.8e-13) and “lipid transport” (*p* = 4.1e-11), pointing to metabolic functions shared by hepatocytes and gut tissues.
4. The clade of 200 genes labeled in pink contains genes enriched in spleen and liver only and point to erythropoiesis. Its top GO terms are mainly related to maturing red blood cells, such as “tetrapyrrole biosynthetic process” (*p* = 1.2e-20) and “erythrocyte development” (*p* = 3.9e-10). DNA motif analysis of promoters in this clade revealed a significant enrichment of Tal1:Gata1, a known pair of regulators essential for hematopoiesis.
5. Members of the clade of 91 genes labeled in blue are mainly expressed in late stage liver and are hepatic functions, as well as in the adrenal gland. Top GO terms include “monooxygenase activity” (*p* = 1.1e-31), “steroid hydroxylase activity” (*p* = 1.7e-20) and “steroid metabolic process” (*p* = 5.3e-9). More than a quarter of these are proteincoding components of cytochrome P450, which are involved in steroid and drug metabolism. Additionally, six sulfotransferase genes are also in this group. Sulfotransferase plays an important role in the metabolism of drugs, hormones and bile acids.
6. Lastly, the clade of 155 genes labeled in yellow show more constant levels through time in liver, with additional expression detected in adrenal gland, kidney, stomach and intestine. Its top GO terms include “blood coagulation” (*p* = 1.7e-29) and “alpha-amino acid metabolic process” (*p* = 3.1e-12), with six coagulation factors, six complement factors, fibrinogens and regulators (protein C and serpins) are found in this clade. **Extended Data Fig. 2-11** Cluster 11 from hierarchical clustering analysis of bulk RNA samples.

1. The main theme of Cluster 11 is gut development and differentiation. Its genes are most highly expressed in intestine and are also enriched in stomach, with sharing of specific clades with either kidney (purple) or CNS (blue) tissues.. For the stomach, E14.5 and P0 timepoints show lower expression for multiple clades, which likely reflects systematic dissection differences at the boundaries between the two gut tissues.
2. Top GO terms are mainly about intestine structure, including “brush border” (*p* = 2.3e-11) and “brush border membrane” (*p* = 3.1e-9). Interestingly, out of 16 genes contributing to the term “brush border”, 8 are in the small clade of 43 genes labeled in purple. This clade also has prominent increasing expression in kidney, representing a shared program of brush border genes between kidney and intestine. Other terms include “sodium ion transport” (*p* = 1.0e-6), “digestive system process” (*p* = 1.0e-6) and “alpha-amylase activity” (*p* = 1.8e-6). Additionally, several gut hormones or peptides are found in this group such as cholecystokinin, gastrin, vasoactive intestinal polypeptide, ghrelin, glucagon and insulin genes. **Extended Data Fig. 2-12.** Cluster 12 from hierarchical clustering analysis of bulk RNA samples.

1. Most genes in Cluster 12 are expressed widely and with an increasing trend, except in the liver, where most of the cluster is depleted at all times.
2. The most prominent secondary theme is strong up-regulation at birth in multiple organs.
3. No GO terms were significantly enriched.
4. This cluster, and subclusters within, are candidates for novel DNA sequence motif-derivation or for correlated mir-signatures that could mediate the birth transition pattern and/or the liver suppression pattern **Extended Data Fig. 2-13.** Cluster 13 from hierarchical clustering analysis

1. Genes in Cluster 13 are mostly enriched in lung, especially at later stages.
2. Partly because of the small cluster size, Gene Ontology didnt provide highly significant terms. However, 4 surfactant-associated proteins contributing to the term “multivesicular body” (*p* = 4.3e-6) are included in this cluster, indicating a possible link to Type II alveolar cells in the lung. **Extended Data Fig. 2-14** Cluster 14 from hierarchical clustering analysis.

1. Cluster 14 contains genes that are highly expressed in stomach. Most are also highly expressed in limb and craniofacial prominence at very late stages. About a quarter of them are also expressed in the P0 bladder.
2. The top GO terms are “cornified envelope” (*p* = 6.7e-26), “keratinization” (*p* = 1.7e-27), “epidermis development” and “keratinocyte differentiation” (*p* = 1.6e-16). The cornified envelope is composed of a layer of dead cells found in skin epidermis and forestomach for protection against the environment. Its major components include loricrin, filaggrin, Involucrin, keratins and small proline-rich protein (SPR) genes that are all found in this cluster, together with the genes required for generating the cornified envelope, such as transglutaminase, cystatin and envoplakin. **Extended Data Fig. 2-15.** Cluster 15 from hierarchical clustering analysis of bulk RNA samples.

1. Genes in this group are coherently enriched in specific replicate samples, but they do not reproduce between replicates or among related tissues.
2. Almost all genes in this cluster are annotated as known pseudogenes or protein-coding genes with low mappability. These low-mappability genes top-abundance mappable counterparts (their corresponding protein-coding genes or paralogs) do not display similar variation (data not shown). **Extended Data Fig. 2-16.** Cluster 16 from hierarchical clustering analysis of bulk RNA samples.

1. The broad theme of this cluster is expression in most tissues and organs, with the exception of the CNS and liver, both of which show little expression.
2. Over the developmental time course, most members increase in limb and craniofacial prominence but are relatively less changing or decreasing in other tissues.
3. The top GO terms of Cluster 16 are dominated by extracellular matrix (ECM) components, such as “extracellular matrix” (*p* = 8.7e-58), “extracellular region part” (*p* = 6.0e-42) and “basement membrane” (*p* = 4.4e-34). Other significant terms include “regulation of cell migration” (*p* = 8.0e-26), “angiogenesis” (*p* = 1.1e-25) and “‘cell junction” (*p* = 2.3e-18).
4. This cluster contains two major clades, highlighted in purple and blue, that share expression in bladder, kidney, lung, stomach, intestine, limb and craniofacial prominence. The blue clade is distinct in also showing strong expression in heart. Their GO terms identify different biases. The purple clade features “occluding junction” (*p* = 7.6e-9) in addition to ECM terms, while “angiogenesis” is absent. The blue clade includes most cluster 16 thematic terms, but also emphasizes “anchoring junction” (*p* = 7.3e-19) and particularly “adherens junction” (*p* = 3.0e-18), consistent with epithelial/endothelial cell junction formation and tube morphogenesis. Thus the purple clade focuses on tight junctions that consist of an epithelial barrier and molecular gate between a cell mass and the environment, while the blue clade concentrates on angiogenesis and adherens junctions that link cells together and also carry cadherin receptors important for tissue morphogenesis. **Extended Data Fig. 2-17.** Cluster 17 from hierarchical clustering analysis of bulk RNA samples.

1. This cluster is divided into two major clades. The upper clade, similar to cluster 15, is coherently enriched in a few individual samples that do not replicate nor do they reproduce among related tissues. Apart from these individual samples, the pattern is noisy across developmental time. This pattern does not correspond to any known dissection or global QC issue. Also similar to cluster 15, no GO term enrichment was found.
2. The lower clade contains genes that are widely expressed among different tissues that are also systematically depleted in the E11.5 and E14.5 samples. This reflects known batch effects at tissue collection/dissection steps (see also CCA analysis) **Extended Data Fig. 2-18** Cluster 18 from hierarchical clustering analysis of bulk RNA samples.

1. Most genes in Cluster 18 are enriched in the craniofacial prominence at early stages but not later.
2. Its top GO terms mainly concern eye development, including “structural constituent of eye lens” (*p* = 1.6e-17) and “eye development” (*p* = 1.1e-15). Genes include crystallins, retinoic acid-metabolizing enzymes (*Cyp26a1* and *Cyp26c1*), lens membrane protein (*Lim2*), melanin regulators (*Tyrp1*, *Tyr* and *Pmel*) and one developmental regulator (*Vax2*).
3. The dissection plan for cranioface was to exclude the eyes, but at earlier stages it appears not to have been fully successful. The expression pattern and Gene Ontology, of Cluster 18 genes in the early craniofacial prominence samples (E10.5, E11.5 and E12.5), including sharp transitions between adjacent timepoints, are likely due to imperfect removal of early eyes. Sporadic enrichment of these genes in later stage craniofacial prominence samples (E16.5) is likely due to a few imperfect dissections in a large embryo pool. **Extended Data Fig. 2-19.** Cluster 19 from hierarchical clustering analysis of bulk RNA samples.

1. Genes in this cluster are mostly enriched in bladder, kidney, limb and neural tube. Within these tissues, expression levels are relatively constant over developmental time.
2. The lower half of this cluster contains 5’ Hox genes (*Hox9*-*Hox13*) and lincRNAs localized in the 5’ region of *Hox* clusters (*Hotair* and *Hottip*). Genes with names beginning with “Gm” that are clustered together with Hox genes are also localized in 5’ Hox gene regions, suggesting shared transcriptional regulatory elements or RNA precursors. Among these *Hox*-cluster genes, there are distinctions, with 5’ *Hox* expression being more abundant in posterior tissues (e.g. bladder, kidney and intestine), consistent with previous findings.
3. As the time course begins at E10.5, we could not follow the well-known upregulation sequence of Hox genes which displays “temporal co-linearity”, except for a gradual increase in *Hoxc12* and *Hoxc13* in the limb, which represent the distal ends of limbs and whose upregulation pattern is late enough to be captured in our time window.
4. In E14.5 neural tube samples the 5’ most *Hox* genes *Hox11-13* are missing, because that batch of embryo dissections did not include the posterior tip of the tube.
5. Four major urinary protein (MUP) genes are enriched in limb, similar to the MUP paralogs in Cluster 2. However, unlike those in Cluster 2, they are also enriched in early neural tube samples. **Extended Data Fig. 2-20.** Cluster 20 from hierarchical clustering analysis of bulk RNA samples.

1. Overall, genes in Cluster 20 are prominently absent from the liver at all stages and are absent or strongly reduced at P0 in most tissues having P0 data. The top ∼1/3 of the cluster contributes little to the two major expression and GO themes. It contains mainly pseudogenes and lncRNAs. In limb, craniofacial prominence and brain, depletion of some of these genes is evident at E11.5 and E14.5 similar to Cluster 17, and possibly related to the batch effect discussed before.
2. In the remaining bottom 2/3, there is considerable substructure among expressing tissues due to the two major biological themes: The first GO enrichment theme is tissue morphogenesis and development, such as “skeletal system morphogenesis” (*p* = 3.3e-13), “branching morphogenesis of an epithelial tube” (*p* = 5.5e-12), “sensory organ development” (*p* = 7.7e-11), “odontogenesis” (*p* = 1.3e-10), “gland development” (*p* = 4.5e-10), “ossification” (*p* = 9.1e-10), “limb morphogenesis” (*p* = 2.1e-9) and “kidney development” (*p* = 6.1e-9). The second GO theme is Wnt signaling, such as “regulation of Wnt signaling pathway” (*p* = 8.2e-12), “Wnt signaling pathway” (*p* = 2.0e-10) and “Wnt-protein binding” (*p* = 3.2e-10). Although this cluster called terms covering a variety of different aspects of development demonstrated in the first theme, the driving genes are often shared among multiple terms referring to different tissues. This likely reflects the broad usage of these signaling pathways in patterning and morphogenesis. Moreover, roughly a quarter of the genes contributing to any Theme 1 terms also contribute to the Wnt theme. Other Theme 1 genes that do not currently contribute to the Wnt GO terms, such as Irx3, Runx2, TWIST, Bmp4, Tbx1 and Tbx3 have been independently associated with this signaling system. This is consistent with the current appreciation that Wnt signaling plays an important and widely distributed role in different individual anlage, including stem cell renewal.
3. The clade of genes colored in purple are highly enriched in kidney and moderately enriched in limb and craniofacial prominence. The themes are suggested by Gene Ontology with terms “skeletal system morphogenesis” (*p* = 1.2e-6) and “branching morphogenesis of an epithelial tube” (*p* = 2.3e-6). These terms of this clade are consistent with the overall theme of this big cluster but the distinct gene expression pattern suggests intensive usage of this subprogram of genes in the kidney.
4. The clade of genes labeled in blue has prominent enrichment in limbs and craniofacial prominence, and is lower but still detectable and decreasing in other tissues. The top GO terms are similar to those called from the whole cluster, but with much enhanced significance for “embryonic skeletal system morphogenesis” (*p* = 1.8e-15) and “cartilage development” (*p* = 1.5e-9). **Extended Data Fig. 2-21.** Cluster 21 from hierarchical clustering analysis of bulk RNA samples.

1. Genes in Cluster 21 are expressed in all the fetal tissues and decrease over time in most. At P0 the majority are expressed in thymus and spleen but are notably depleted elsewhere.
2. Top GO terms are mainly cell division and nucleus components, such as “chromosomal part” (*p* = 1.2e-93) and “cell cycle process” (*p* = 1.6e-87) consistent with genes involved in executing the cell cycle, especially components of the chromosome and its associated proteins. The global decrease in RNA levels from these genes over time is consistent with shifting from fast growing proliferating cells to more differentiated ones.
3. Fetal hemoglobins are also found in this cluster such as *Hbb-bh1*, *Hbb-y* and *Hba-x*. **Extended Data Fig. 2-22.** Cluster 22 from hierarchical clustering analysis of bulk RNA samples.

1. Genes in Cluster 22 show distinct enrichment at E11.5 and E14.5 stages in some tissues, and they do so reproducibly among the replicates.
2. Most of these genes are pseudogenes and low-mappability protein-coding genes. They are similar to the batch-effect clades in Cluster 17 and Cluster 20 but display the inverse pattern trend. **Extended Data Fig. 2-23.** Cluster 23 from hierarchical clustering analysis of bulk RNA samples.

1. This small cluster contains genes most prominently expressed at early times in CNS tissues. They are also depleted preferentially at E16.5 in many other tissues. Unlike the candidate batch effects of clusters like 15, 17, and 22 that are heavily enriched in pseudogenes, this cluster is not explained by annotated pseudogenes. There was no significant GO enrichment. **Extended Data Fig. 2-24.** Cluster 24 from hierarchical clustering analysis of bulk RNA samples.

1. Genes in this cluster are widely expressed and are preferentially higher in the CNS regions and/or in the developing liver. Most, but not all, increase during development of these tissues.
2. Top Gene Ontology terms are dominated by lipid metabolism, such as “lipid metabolic process” (*p* = 2.7e-13) and “cholesterol biosynthetic process” (*p* = 1.7e-11). Interestingly, all of the nine genes contributing to the term “cholesterol biosynthetic process” are localized in a tiny clade of 23 genes labelled in purple. These 23 genes are all very abundant and highly correlated among themselves. **Extended Data Fig. 2-25.** Cluster 25 from hierarchical clustering analysis of bulk RNA samples.

1. More than half of the genes in Cluster 25 are consistently and highly enriched in the hindbrain and neural tube plus stomach, intestine and adrenal gland. Kidney and lung also express distinct subsets of these genes.
2. This cluster contains most of the 3’ *Hox* genes, almost all located in the two clades labeled in purple and blue. The purple clade consists of the 3’ most *Hox* genes and genes sitting in the 3’ end of *Hox* gene clusters while the blue clade is made of *Hox* genes and non-*Hox* genes in the center (less 3’ but not 5’) of *Hox* clusters. The purple-clade genes are expressed in lung while the blue ones are mostly not. This is probably because lung is relatively anterior to other endoderm tissues assayed, which corresponds to the 3’ end of the endoderm *Hox* A/P axis.
3. For the genes outside the *Hox* gene clades combined, Gene Ontology generated terms related to the neural system, such as “neuron differentiation” (*p* = 1.4e-7); focusing on “enteric nervous system development” (*p* = 2.4e-7).
4. In E14.5 neural tube samples some genes are more depleted compared to E13.5 and E15.5. We think this might result from a dissection protocol detail that produced shorter spinal cord and depleted the 3’-most *Hox* expressing tissue. **Extended Data Fig. 2-26.** Cluster 26 from hierarchical clustering analysis of bulk RNA samples.

1. Genes in Cluster 26 are mostly enriched in forebrain at late stages.
2. *Avp* and *Oxt* encode neuropeptides synthesized in the hypothalamus that regulate complex maternal and sexual behaviors. They are clustered together within 6 kbp on chromosome 2. **Extended Data Fig. 2-27.** Cluster 27 from hierarchical clustering analysis of bulk RNA samples.

1. Cluster 27 contains genes highly enriched in kidney. Most are also expressed in brain and neural tube at later stages but less abundantly than in kidney.
2. *Pax2*, *Pax8* and their target *Gata3* are found in this cluster which specify the nephric lineage and regulate branching morphogenesis in the developing kidney. Pax2 and Pax8 are also reported to specify GABAergic and glycinergic neuronal fates, partly explaining expression in the hindbrain and neural tube. It’s possible that this cluster concerns two independent cell fate specification and morphogenesis programs that use overlapping regulatory factor sets, such as Pax2, Pax8 and Gata3. **Extended Data Fig. 2-28.** Cluster 28 from hierarchical clustering analysis of bulk RNA samples.

1. Genes in Cluster 28 are expressed in many tissues, but with lung and craniofacial prominence being highest, followed by CNS regions. Almost all increase over time, but with differing kinetics in different tissues and brain regions.
2. Most of the significant GO terms are about ciliogenesis, such as “cilium movement” (*p* = 1.4e-19), “cilium” (*p* = 1.2e-17) and “outer dynein arm assembly” (*p* = 1.5e-14). The contributing genes include components of dynein arms and radial spokes, genes coding for assembly machinery such as dynein docking complex, tubulin modifying enzyme and the nexin-dynein regulatory complex. Two known cilium regulators, Foxj1 and Mcidas, are also in this cluster. The cilium is a fundamental structure, with primary cilia being ubiquitous while secondary and sensory cilia having more specialized distributions that correspond well with the pattern for the majority of genes in cluster 28. The pattern can be explained by the emergence of airway cilia in the lung, the airways of the craniofacial prominence, and the ependymal cilia of the CNS. **Extended Data Fig. 2-29.** Cluster 29 from hierarchical clustering analysis of bulk RNA samples.

1. Most genes in this relatively small cluster are distinguished by highest expression in the thymus, but more than half are also expressed substantially in brain or in face/limb, or in kidney/lung and gut.
2. Gene Ontology failed to identify a significantly enriched term for this group. **Extended Data Fig. 2-30.** Cluster 30 from hierarchical clustering analysis of bulk RNA samples.

1. Cluster 30 contains genes expressed most prominently in heart and/or CNS samples, with the admixture among the tissues varying across different clades.
2. Top enriched GO terms mainly identify transport of metal ions, such as “metal ion transport” (*p* = 4.1e-8), “metal ion transmembrane transporter activity” (*p* = 1.3e-7) and “potassium ion transmembrane transporter activity” (*p* = 2.0e-7). **Extended Data Fig. 2-31.** Cluster 31 from hierarchical clustering analysis of bulk RNA samples.

1. Genes in Cluster 31 are mainly enriched in brain and neural tube, with different regionalization for subclusters, plus the facial prominence (perhaps partly driven by cross-contamination of face with forebrain dissection at early times).
2. More than a third of the genes in this cluster are transcription factors (“sequence-specific DNA binding”, *p* = 1.0e-26), most of which also contribute to the GO term “neuron differentiation” (*p* = 4.4e-18). It is likely that this group of genes are involved in neuron maturation, such as *Dlx1*, *Dlx2* and *Helt* which specifies GABAergic neuron differentiation. Genes responsible for cerebral cortex GABAergic interneuron migration (*Lhx6*, *Arx* and *Fezf2*) are also found in this cluster. **Extended Data Fig. 2-32.** Cluster 32 from hierarchical clustering analysis of bulk RNA samples.

1. Genes in Cluster 32 are expressed in nearly all the tissues with increasing trajectories over time, with the notable exception of liver where they are expressed at very low levels and then decrease. This cluster contributes to the global separation of the CNS (where expression is strongest) from the developing liver.
2. Gene Ontology offered little specific annotation, except “positive regulation of adenylate cyclase activity” (*p* = 1.7e-5). **Extended Data Fig. 2-33.** Cluster 33 from hierarchical clustering analysis of bulk RNA samples.

1. Genes in the large cluster 33 are expressed in most tissues prior to P0, except liver. CNS and face/limb are by far the most prominent. Most of these genes are time-course variant. Timecourses in different tissues display distinctive trajectories, with decreasing courses being more common, unlike most other major clusters. Thus genes in this cluster vanish very early in liver; decrease monotonically in kidney, lung, stomach and intestine; remain constant early and slightly decrease at later stages in heart, craniofacial prominence, and limb.
2. Gene Ontology enrichment produced three major themes. First, 159 genes (16%) encode DNA binding proteins – especially transcription factors – contributing to “DNA binding” (*p* = 1.7e-17) and “RNA biosynthetic process” (*p* = 7.5e-14). As discussed in the Results section, zinc finger presumptive repressors are especially prominent (Ext. Data Fig. 5e). Second, this cluster contains genes regulating different aspects of morphogenetic processes, with enrichment in the term “embryonic morphogenesis” (*p* = 1.9e-12). This is similar to Cluster 20, which also has a broadly decreasing pattern, though it features an emphasis on the Wnt pathway that does not apply to Cluster 33. Finally, significant overlaps of Cluster 33 with cell projection-related genes are called by terms “cell projection organization” (*p* = 1.3e-15), “cilium assembly” (*p* = 1.9e-14), “neuron projection guidance” (*p* = 4.4e-11) and “regulation of nervous system development” (*p* = 4.2e-12). This cluster of genes is different from the cilium-related Cluster 28 in expression dynamics, showing opposite temporal trajectories that argue strongly for distinct regulation. **Extended Data Fig. 2-34.** Cluster 34 from hierarchical clustering analysis of bulk RNA samples.

1. The theme of this large cluster is expression in all four CNS tissues, with a dominant upward temporal pattern. While most increase over time, they do so with varying kinetics among subclades and between brain regions. Gene Ontology revealed enrichment for a large number of neuron-identity and structure terms associated with neuronal differentiation and maturation, with the most dominant ones being “synapse” (*p* = 1.0e-93), “neuron projection” (*p* = 6.9e-55), “behavior” (*p* = 2.3e-42) and “regulation of nervous system development” (*p* = 2.9e-34).
2. Apart from the central neuronal theme, subclades (colored purple, blue and pink) differ from each other and from the major neural cluster. All three are significantly enriched with transcription factors and neural development regulators, and they display diverse tissue patterns relative to each other. The small purple clade at the top features genes enriched caudally in neural tube and hindbrain. The blue clade below it is enriched in midbrain and significantly but less so in hindbrain and neural tube, with overall downward trajectories. The pink clade near the bottom features genes expressed earliest in all four CNS regions, diminishing in later stages.

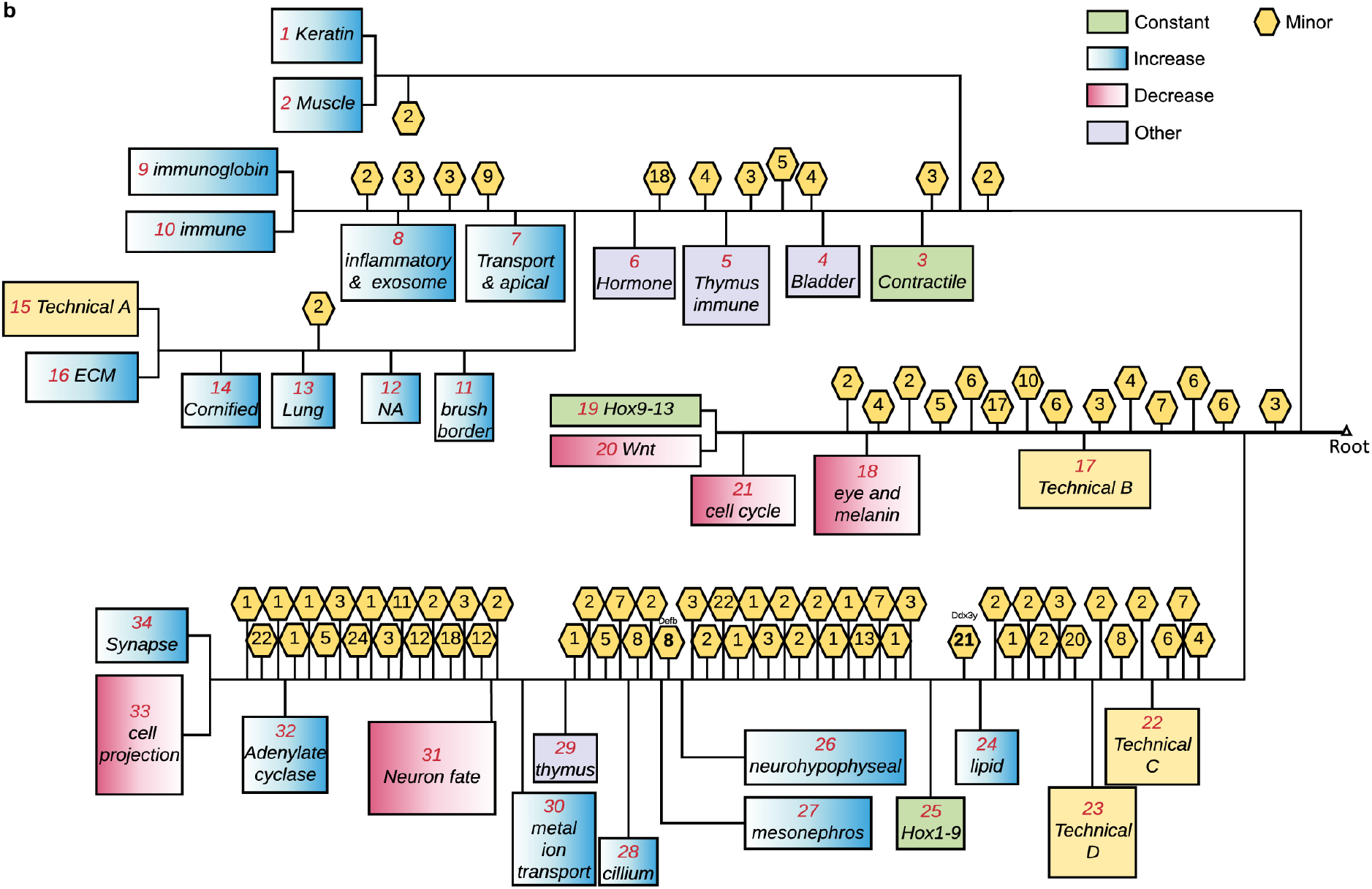

**Extended Data Figure 3:**
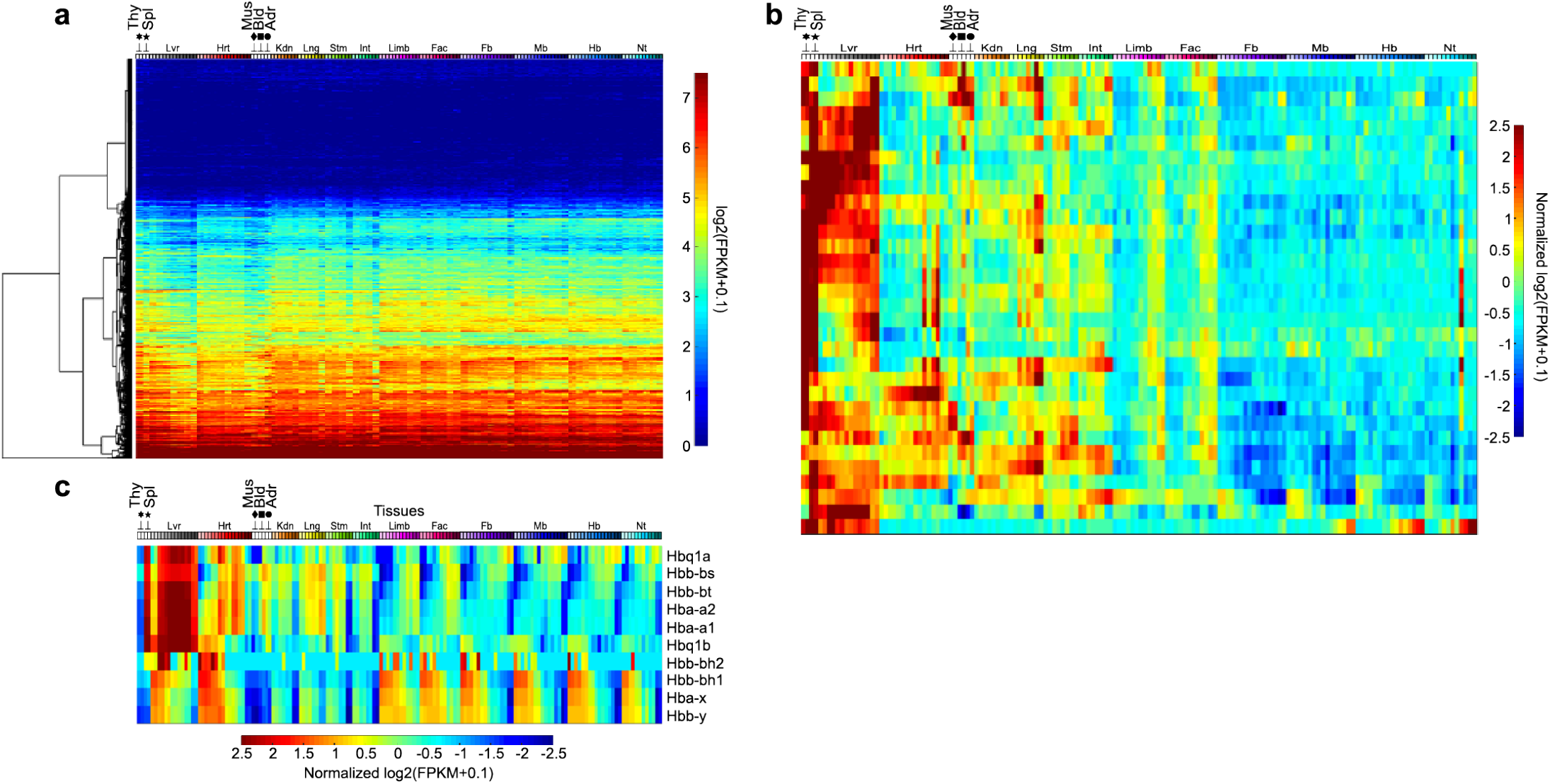
Additional groups of genes with diverse biological implications. In all plots, tissue identities are labeled on top (*x*-axis) matching Fig. 1, and genes are on the *y*-axis. (a) Expression levels of ubiquitous genes are shown in the heatmap according to the scale bar at right (b,c) Normalized expression levels of genes associated with B-cell activation in Cluster 10 (b) and hemoglobin genes (c).

**Extended Data Figure 4:**
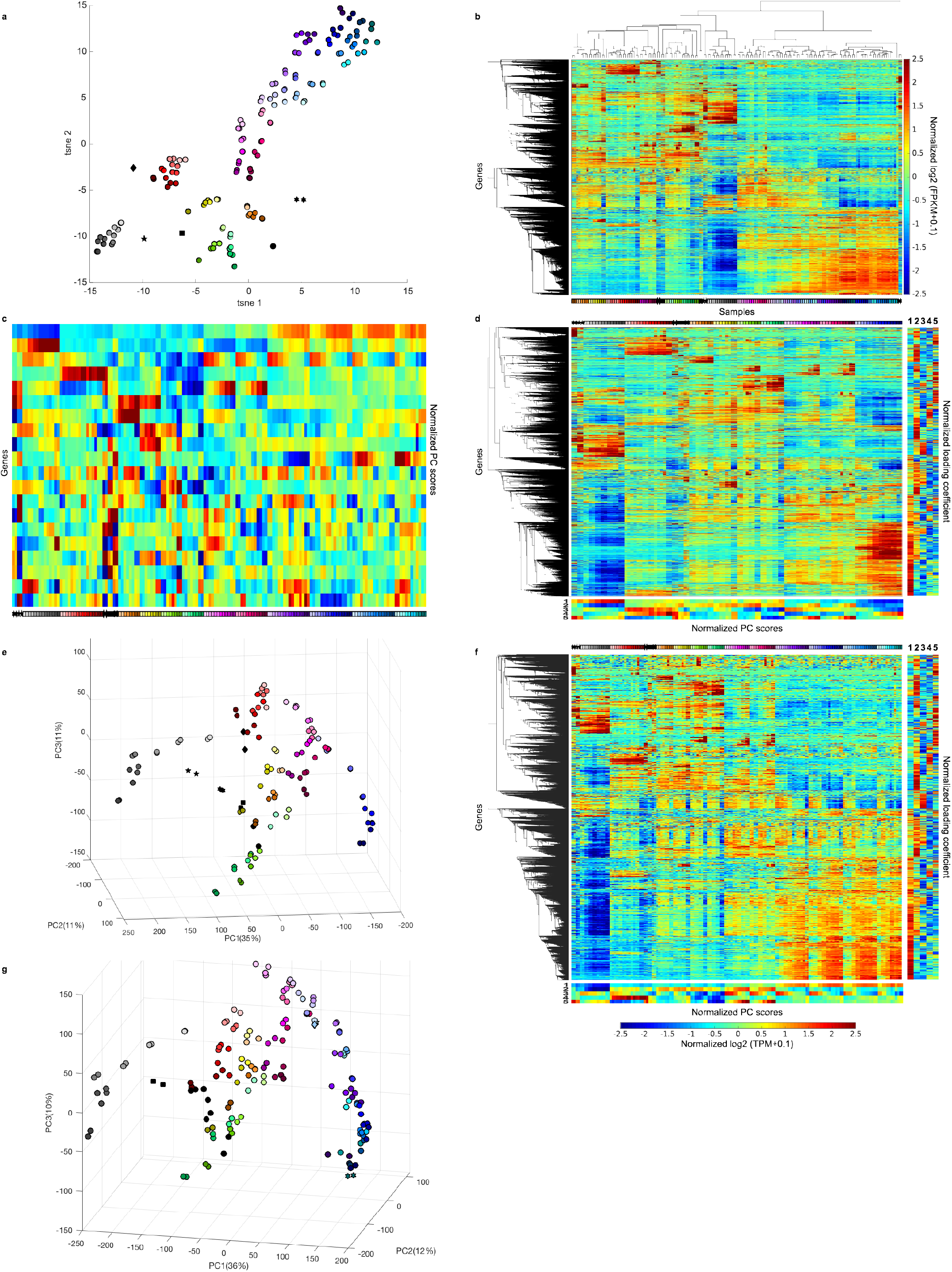
Alternative views of global bulk transcriptome. (a) Bulk tissue transcriptome is organized on a 2D t-SNE plane, with color code as in Figure 1. *n* = 156 bulk RNA-seq libraries (b) Two-way hierarchical clustering of differential genes in bulk data using Pearson correlation. (c) Normalized principal component scores of the top 20 components. Tissue identities and stages are labeled at the bottom following color codes in Fig. 1 (d,e) One-way hierarchical clustering (d) and PCA projection (PC scores are labeled at the bottom and loading coefficients are on the right) (e) of whole transcriptome with forebrains, hindbrains and neural tubes removed to test robustness. *n* = 112 bulk RNA-seq libraries. Color codes as in Figure 1. (f,g) One-way hierarchical clustering (f) and PCA projection (similar to (e)) (g) of whole transcriptome quantified by TPM instead of FPKM. *n* = 156 bulk RNA-seq libraries. Color codes as in Figure 1.

**Extended Data Figure 5:**
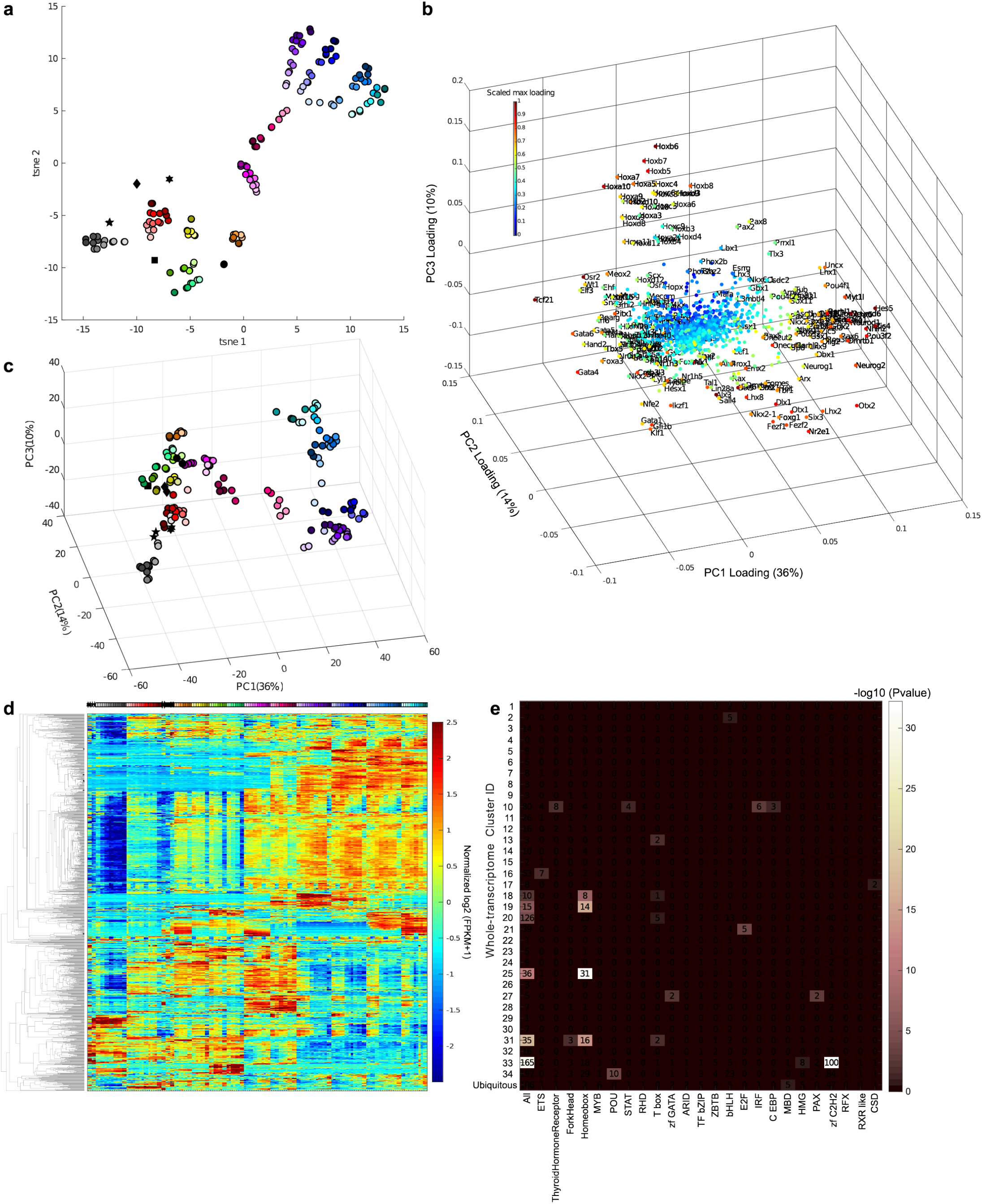
Transcription factor expressions in the bulk data. Color codes in (a–d) as in Figure 1. (a) *t*-SNE representation of transcription factor expression profiles; *n* = 156 bulk RNA-seq libraries. (b) 3-D projection of PC loading coefficients of transcription factors. (c) 3-D projection of PC loadings of transcription factor expression profiles; *n* = 156 bulk RNA-seq libraries. (d) one-way hierarchical clustering of transcription factor expressions in bulk data. Tissue identities are labeled following color codes in Fig. 1; *n* = 156 bulk RNA-seq libraries. (e) abundance representation of transcription factor families in individual bulk expression clusters. Colors indicate Bonferroni-corrected *p*-values from hypergeometric test.

**Extended Data Figure 6:**
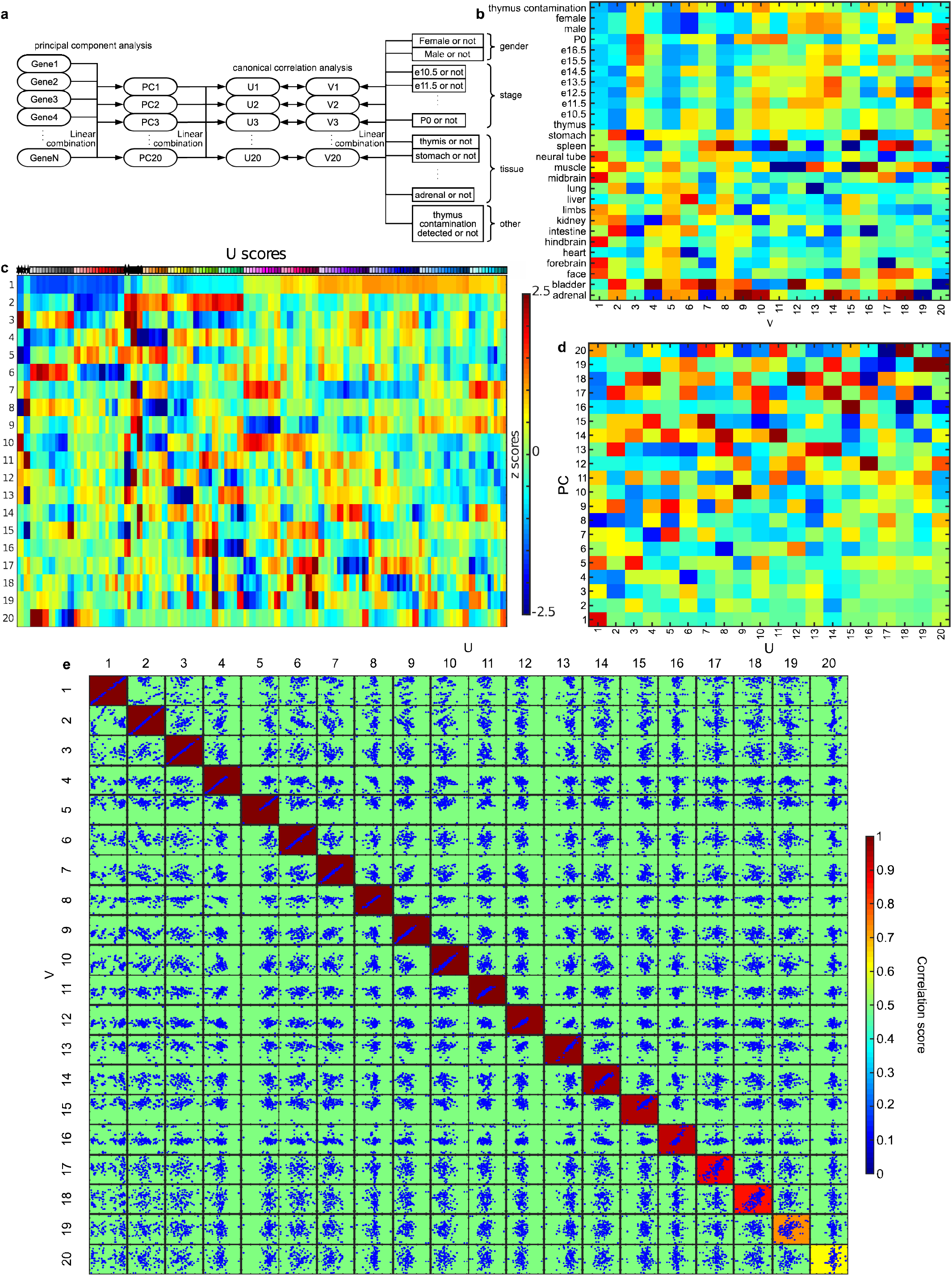
Canonical correlation analysis of the bulk data. (a) A diagram showing the setup of canonical correlation analysis (more details in Supplementary Data 3). (b) The vertically normalized loadings of the Boolean metadata variables. Tissue identities are labeled with color codes in Fig. 1. (c) The horizontally normalized scores of CCA variables across tissue samples. (d) The vertically normalized loadings of principal components. (e) The correlations between U and V variables. Pairwise relationships between U and V variables are shown by the corresponding scatter plots and heatmap representing the Pearson correlation coefficient; *n* = 156 bulk RNA-seq libraries.

**Extended Data Figure 7:**
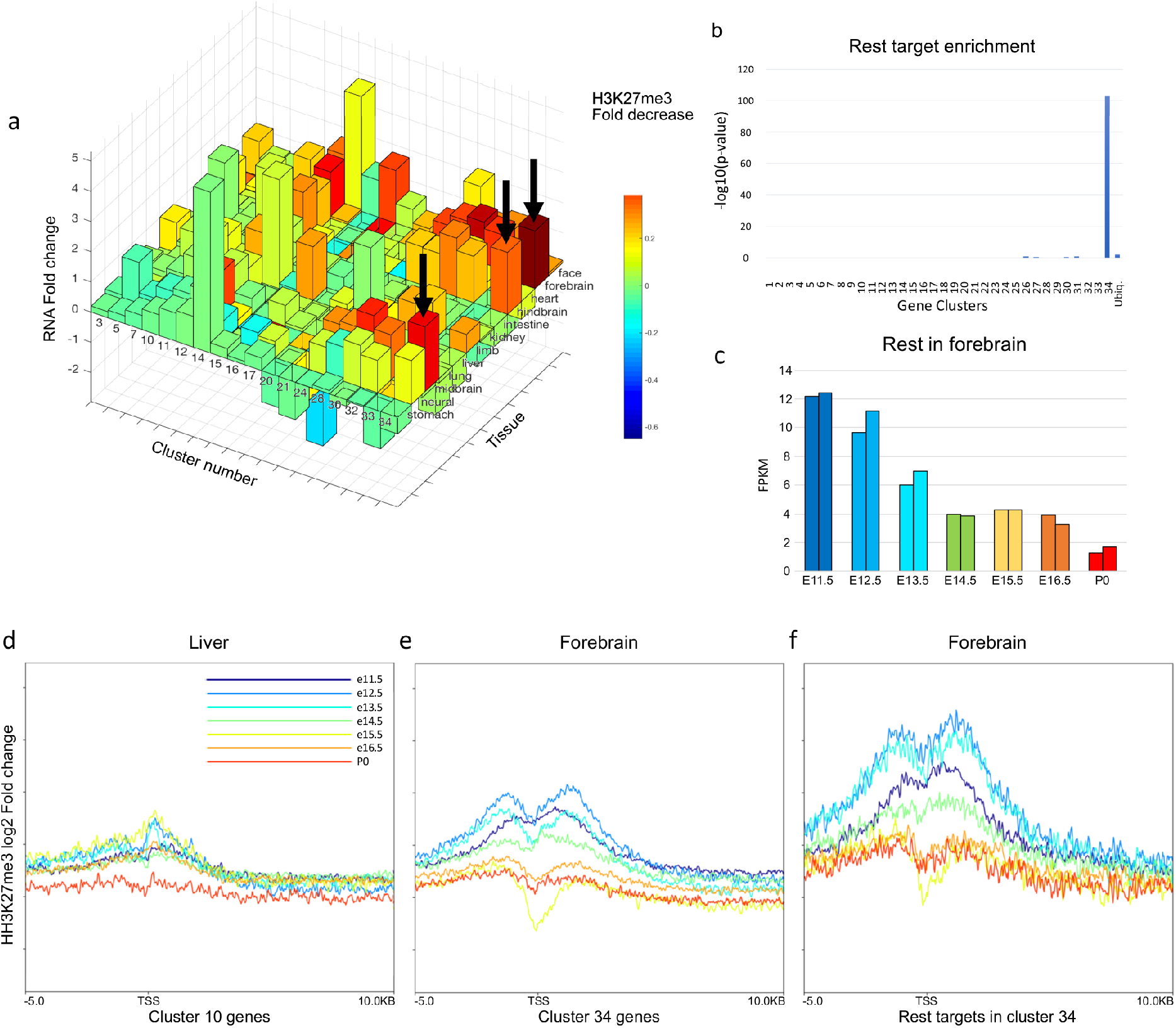
CNS-specific genes are associated with REST/NRSF binding and de-repression. (a) H3K27me3 fold-decrease and RNA fold-change. Each bar represents a cluster of genes in a tissue type. The height represents RNA fold-increase between the earliest and latest time points while the colors represent H3K27me3 ChIP signal fold decrease. The arrows point to the strongest decrease of H3K27me3 that happens in Cluster 34 in brain samples. (b) REST/NRSF target enrichment in individual clusters. Bonferroni-corrected *p*-values are calculated based on Hypergeometric tests. Sample size (equal to that of Ext Data Fig 2): Cluster 1 *n* = 21 genes, Cluster 2 *n* = 196 genes, Cluster 3 *n* = 693 genes, Cluster 4 *n* = 65 genes, Cluster 5 *n* = 474 genes, Cluster 6 *n* = 95 genes, Cluster 7 *n* = 226 genes, Cluster 8 *n* = 106 genes, Cluster 9 *n* = 103 genes, Cluster 10 *n* = 2,182 genes, Cluster 11 *n* = 563 genes, Cluster 12 *n* = 536 genes, Cluster 13 *n* = 93 genes, Cluster 14 *n* = 341 genes, Cluster 15 *n* = 219 genes, Cluster 16 *n* = 1,176 genes, Cluster 17 *n* = 338 genes, Cluster 18 *n* = 37 genes, Cluster 19 *n* = 45 genes, Cluster 20 *n* = 1,319 genes, Cluster 21 *n* = 801 genes, Cluster 22 *n* = 44 genes, Cluster 23 *n* = 95 genes, Cluster 24 *n* = 283 genes, Cluster 25 *n* = 138 genes, Cluster 26 *n* = 30 genes, Cluster 27 *n* = 68 genes, Cluster 28 *n* = 200 genes, Cluster 29 *n* = 56 genes, Cluster 30 *n* = 236 genes, Cluster 31 *n* = 90 genes, Cluster 32 *n* = 256 genes, Cluster 33 *n* = 1,008 genes, Cluster 34 *n* = 3,073 genes, Ubiquitous *n* = 3,000 genes. (c) Abundance of *Rest* mRNA in forebrain. The individual data points are shown as individual bars. (d–f), Averaged H3K27me3 profiles near promoter regions (*x*-axis) for liver ChIP-seq signals over Cluster 10 genes (d), forebrain ChIP-seq signals over Cluster 34 genes (e) and forebrain ChIP-seq signals over Rest-targeted genes in Cluster 34 (f).

**Extended Data Figure 8:**
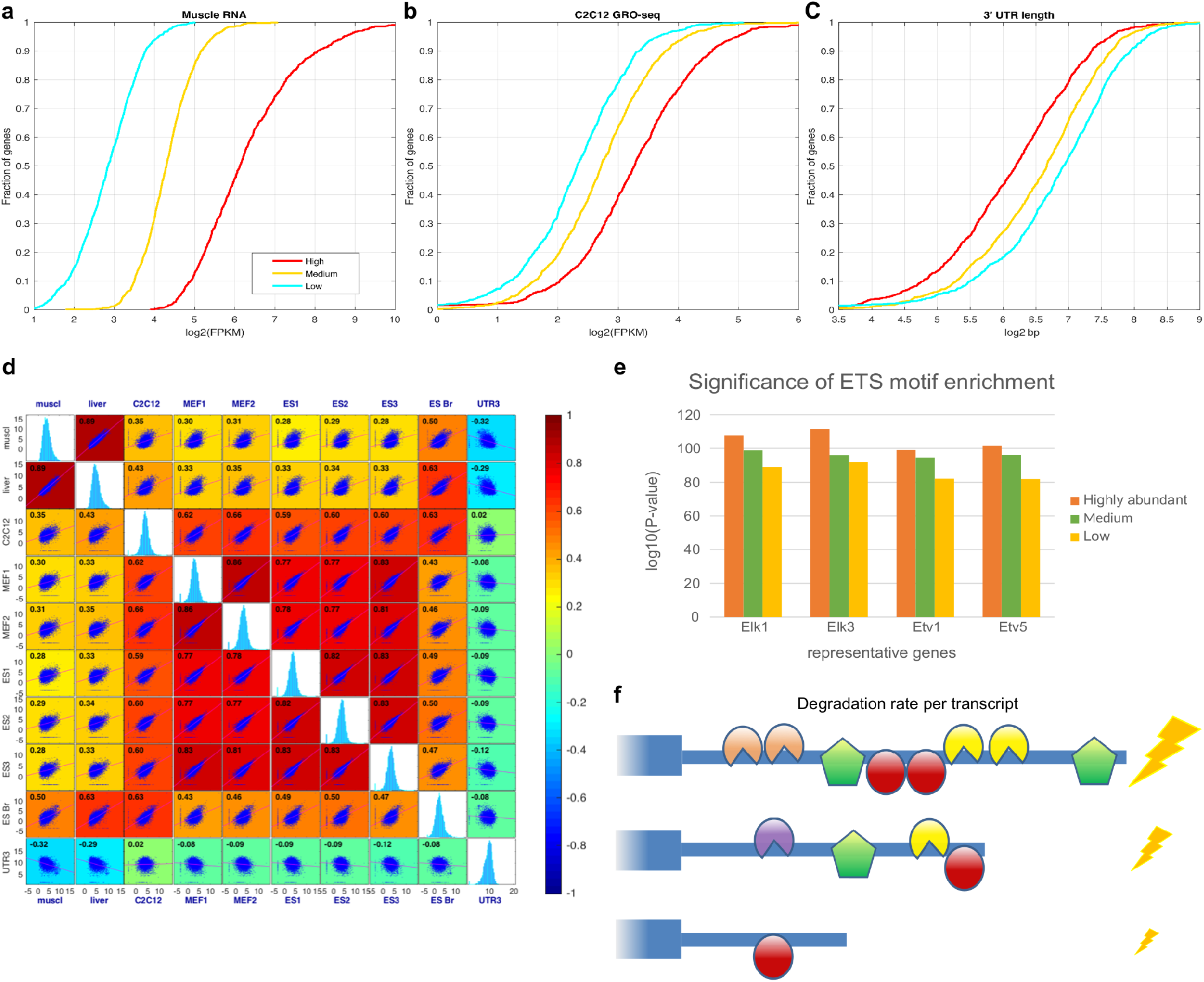
Regulatory mechanisms of ubiquitous genes. (a–c) Cumulative distribution function plots of polyA+ RNA-seq measurements from Skeletal muscle (a), C2C12 GRO-seq data (b) and average 3’ UTR length (c) are compared among three equal-sized groups of ubiquitous genes defined by their RNA-seq abundance. (d) Comparisons of 3’ UTR length, GRO-seq, Bru-seq and polyA+ RNA-seq assays among multiple different samples. Pearson correlation scores between each pair of measurements on the columns and rows are visualized using a heatmap. In the corresponding cell of the comparison, a scatter plot is provided. On the diagonal are histograms of each individual measurement; *n* = 24,832 detectable genes. (e) Significance of ETS motif enrichment in the promoters of ubiquitous genes determined using AME in MEME suite. *n* = 1,000 each for high, medium and low groups. (f) A model is proposed that longer 3’ UTR may harbor more binding sites for RNA-decay apparatus, leading to lower abundance at steady states.

**Extended Data Figure 9:**
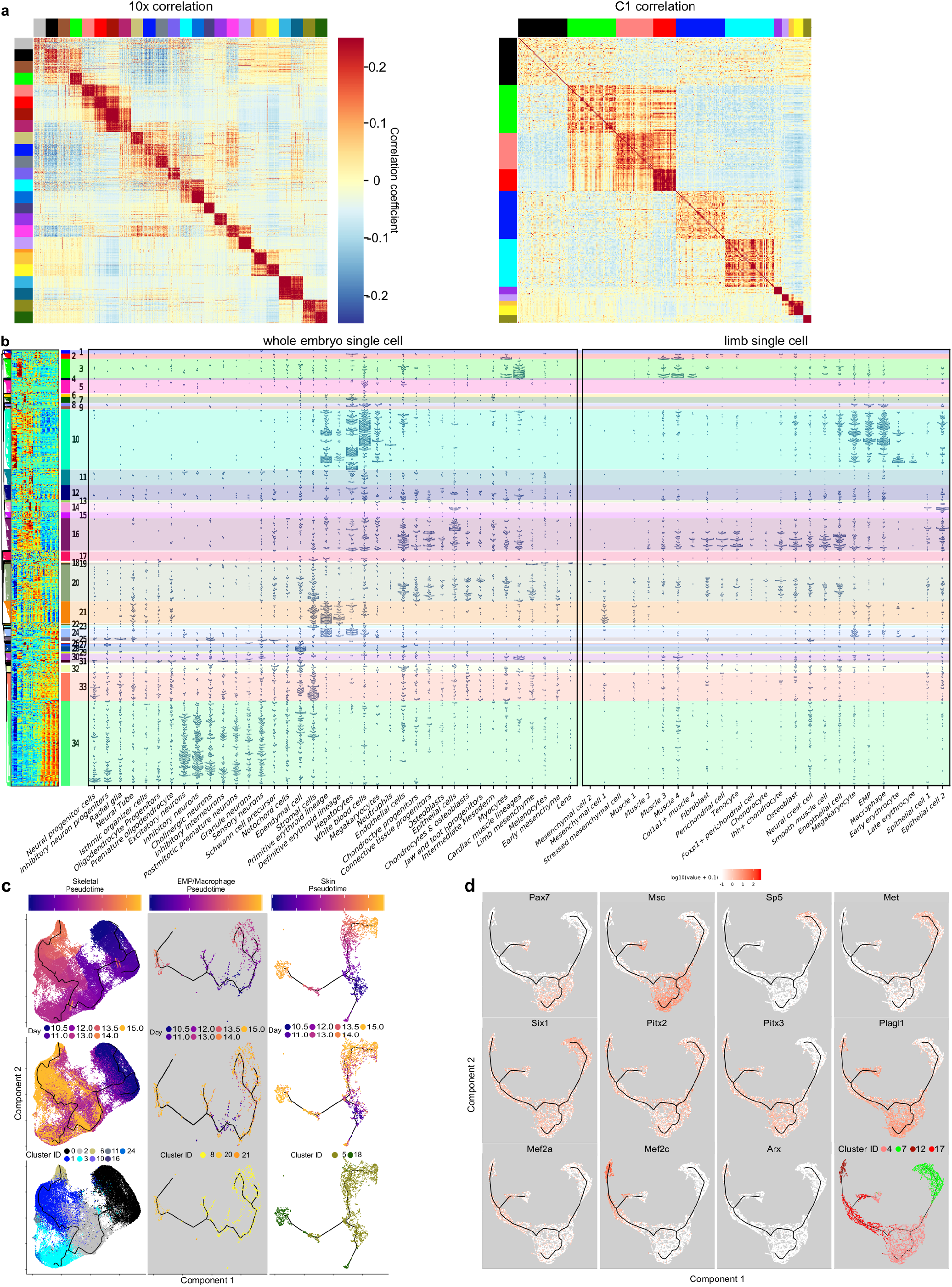
Cell-type relationships inferred from single-cell data. (a) Cell-cell correlations. Feature genes were used to calculate and visualize Pearson correlation coefficients between cells. Specific cell-type populations (indicated by color bands on the axes) were downsampled to 100 (10x) or 30 (C1) (b) Comparing single-cell data with bulk. As an extended version of Fig. 1d, this comparison added a panel for 10x single-cell RNA-seq limb data (far right). (c) Lineage inference. Skeletal (left), myeloid (middle) and skin (right) cell types were used for lineage inference, respectively. Pseudotime, developmental time and cell type are presented from top to the bottom. (d) Selected transcription factor expressions are displayed on the Monocle graphs produced from the 10x data for the 4 cell types comprising the myogenic lineage. (e) Feature gene expression profiles of C1 single cells. Normalized *log*-transformed FPKM values (*y*-axis) are used for hierarchical clustering using Spearman coefficients with complete linkage. Major cell types (*x*-axis) together with an *Lmo2* + mesenchyme subtype are highlighted using colors corresponding to Fig. 3b. The overall picture showed different numbers of marker genes across cell types. (f,g) CIBERSORT deconvolution of bulk data. CIBERSORT was used to deduce proportions of major cell types (*y*-axis) present in staged samples (*x*-axis) of independently produced forelimbs (f) and ENCODE mixed limb materials (g). The color codes match Fig. 3b. (h) Monocle lineage inference for four skeletal muscle clusters including Cluster 22. Pseudotime, developmental time and cell type are shown on the left, while marker gene expression is mapped on the right. (i) 20 micron sections of mouse E13.5 forelimb double-immunostained for Osr1 (green) and Myog (red) (left), DAPI blue counterstain (right). All images taken with 63× oil immersion objective. Images in upper panels are enlarged from boxed areas in lower panels. Arrowheads: green: Osr1(+) Myog(−) nucleus. Red: Myog(+) Osr1(−) nucleus. White: double (+/+) cells. Immunocytochemistry was repeated 3 times independently. (j) Heatmaps for all TFs that scored as differential genes for cell types in the myogenic lineage plus limb resident mesenchyme, at 0.2 *δ*pct cutoff, downsampled to 100 (10x) or 30 (C1). Color code per Fig. 3b, Extended Data Fig. 9e.

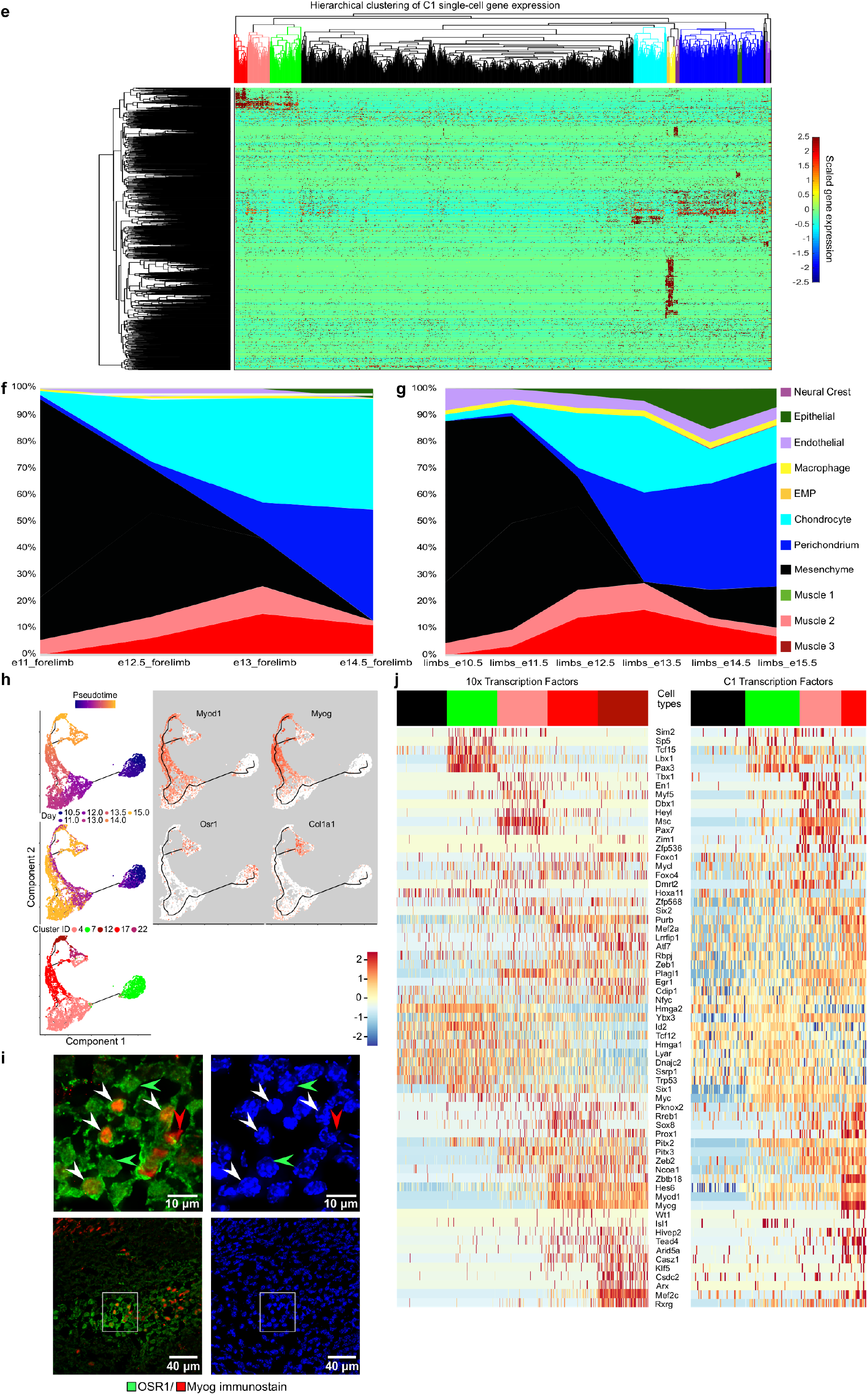

**Extended Data Figure 10:**
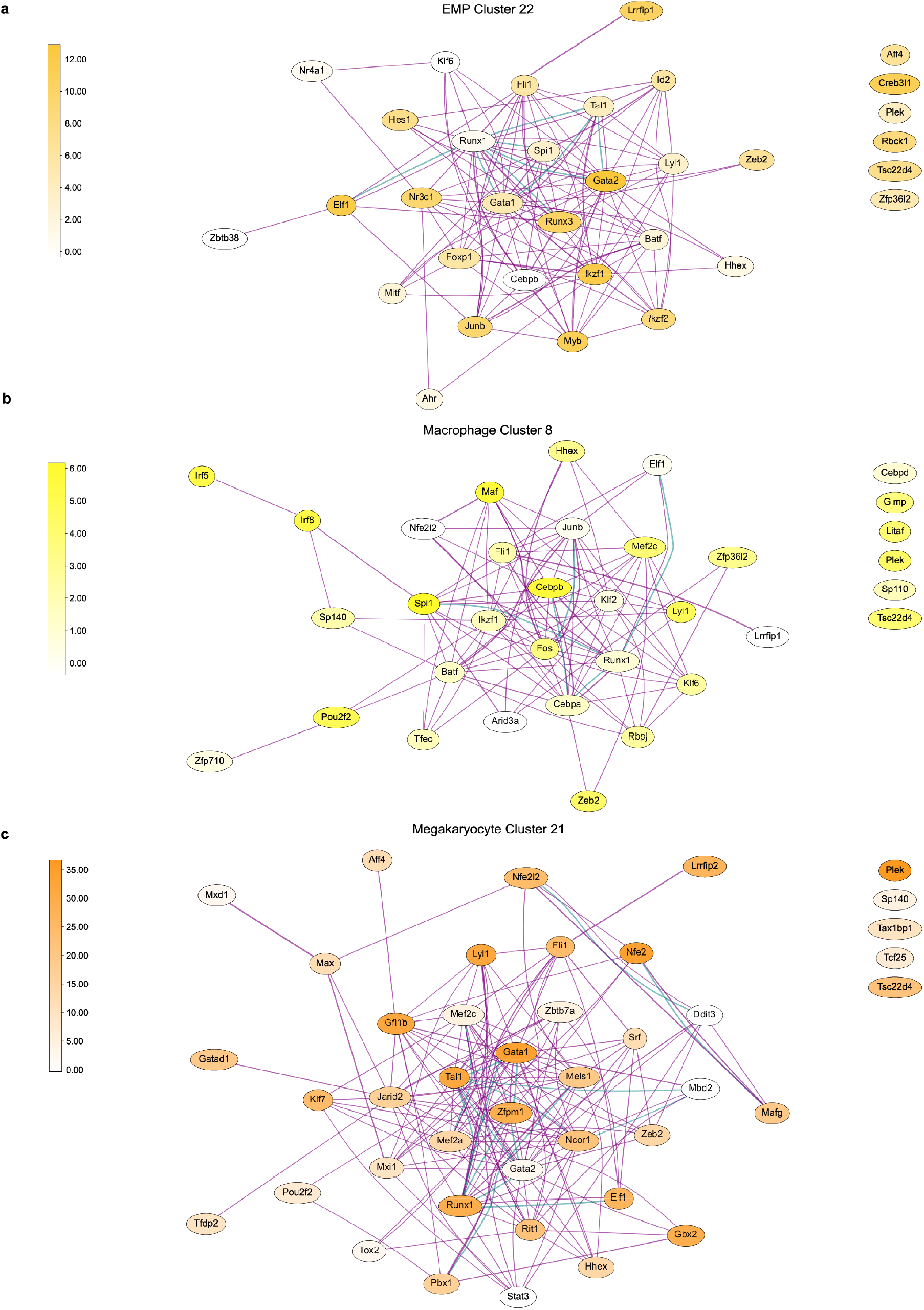
Cell type-specific transcription factor repertoire. (a-r) Transcription factors among the marker genes of each cell type (see Methods) were used for interaction analysis by StringDB, and the results were organized into graphs. Both database (teal) and experimental (magenta) evidence codes were used to produce edges. Node colors reflect the abundance of RNA-Seq expression measurements in the 10x data.

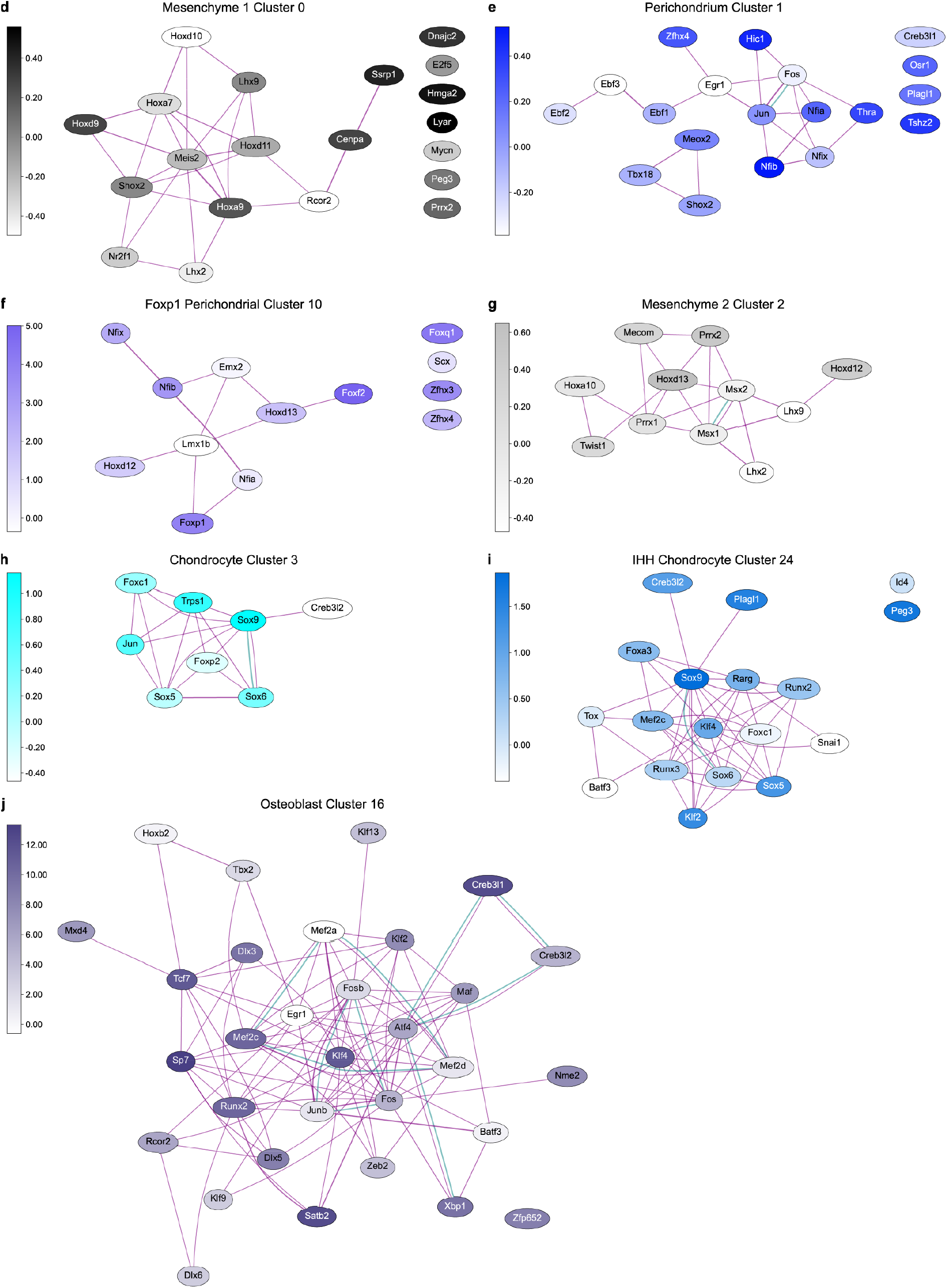

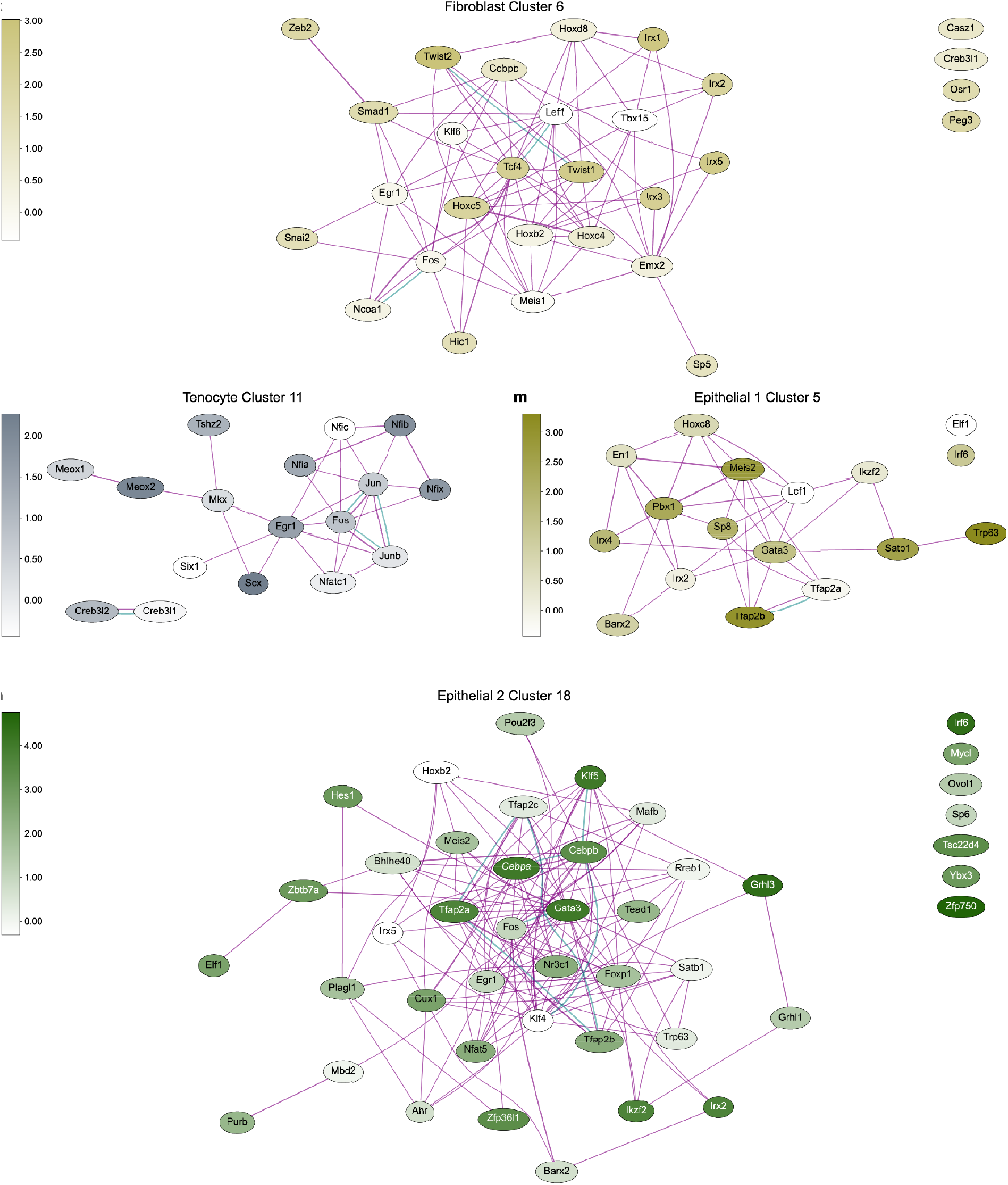

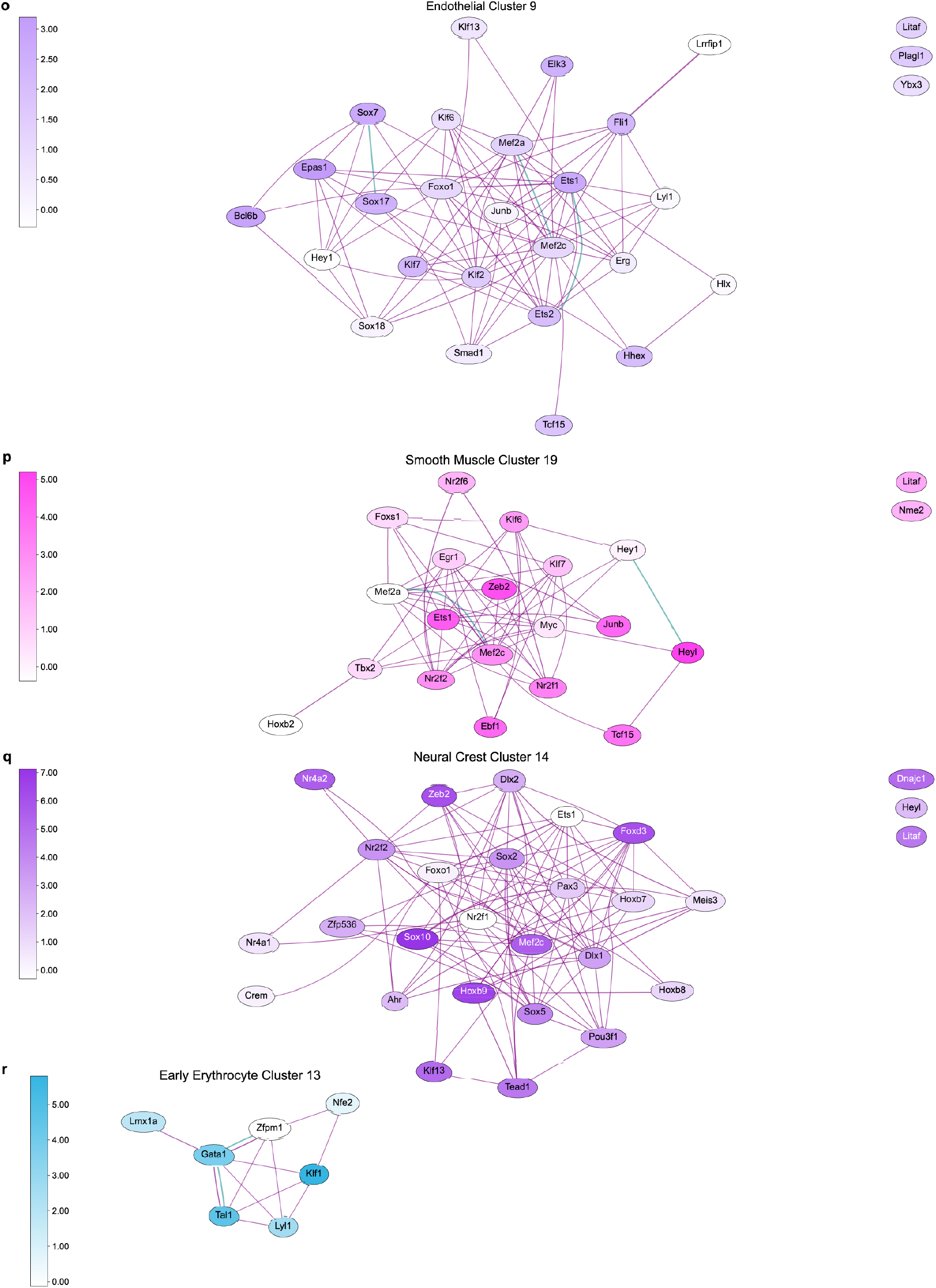

**Extended Data Figure 11:**
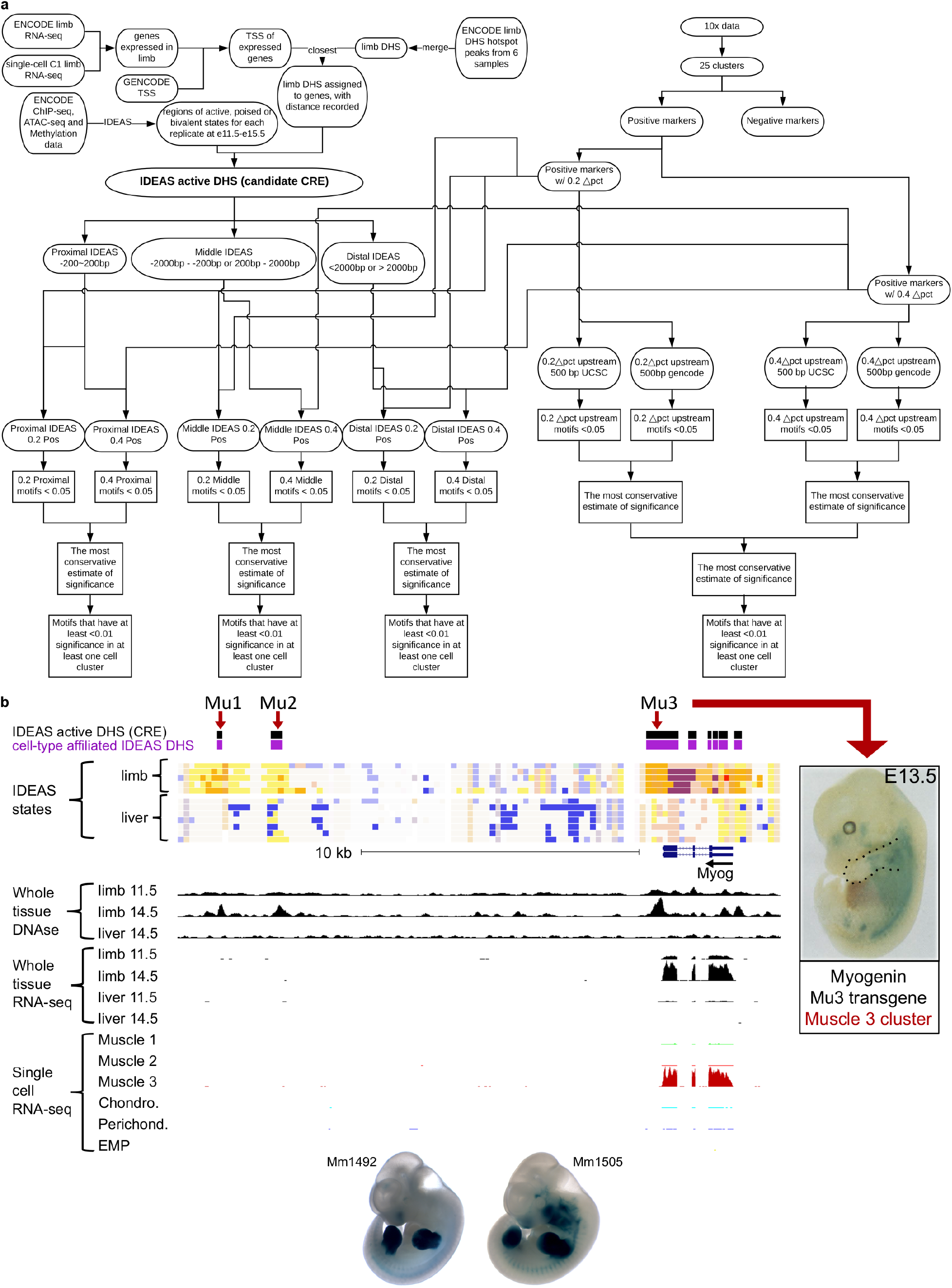
*Cis*-regulatory element analysis using ENCODE chromatin data and single-cell RNA-seq data. (a) A flowchart of the analysis (b) Computationally predicted regulatory elements at the *Myog* locus. From the top to the bottom are the tracks for limb IDEAS active DHS (black bars), cell-type affiliated ones among the former (purple bars), IDEAS scores of limb and liver samples from early to late timepoints, bulk DNAse-seq raw data, bulk RNA-seq raw data, and aggregated C1 single-cell RNA-seq data per cell type. Validation of the Mu3 element in mouse embryo by enhancer assay is also included at the bottom right. (Modified, with permission, from Yee and Rigby 1993, Cold Spring Harbor Laboratory Press) ^64^. Lower: examples of limb-positive enhancer results from the VISTA database that are not cell-type-specific. (c) UCSC genome browser visualization of the C1qb locus, which is in the limb macrophage cluster. Three cEnhs for limb-specific expression of this macrophage gene were identified (Lb1-3). (d) Enriched motifs over regulatory elements. Motifs enriched at the distal elements or promoters of positive and negative markers for each cell type found in the 10x data (see Methods) are visualized using a similar method as Fig. 2b. Colors and numbers of the round nodes correspond to 10x cell type identities (legend at lower right), while the grey and yellow ovals represent shared and unique motifs, respectively. (e) Comparison between FANTOM5 detected promoters and enhancers with promoters and enhancers detected in this study. Elements labeled as “other” are either active or poised in limb generally, but are not cell-type preferential. (f) UCSC Genome Browser shot at the MyoD1 locus. DHS locus accessibility data are shown, along with H3K4me2 and H3K4me3 histone ChIP-Seq data. Blue arrows indicate regions of early chromatin accessibility (see text and Fig. 4a).

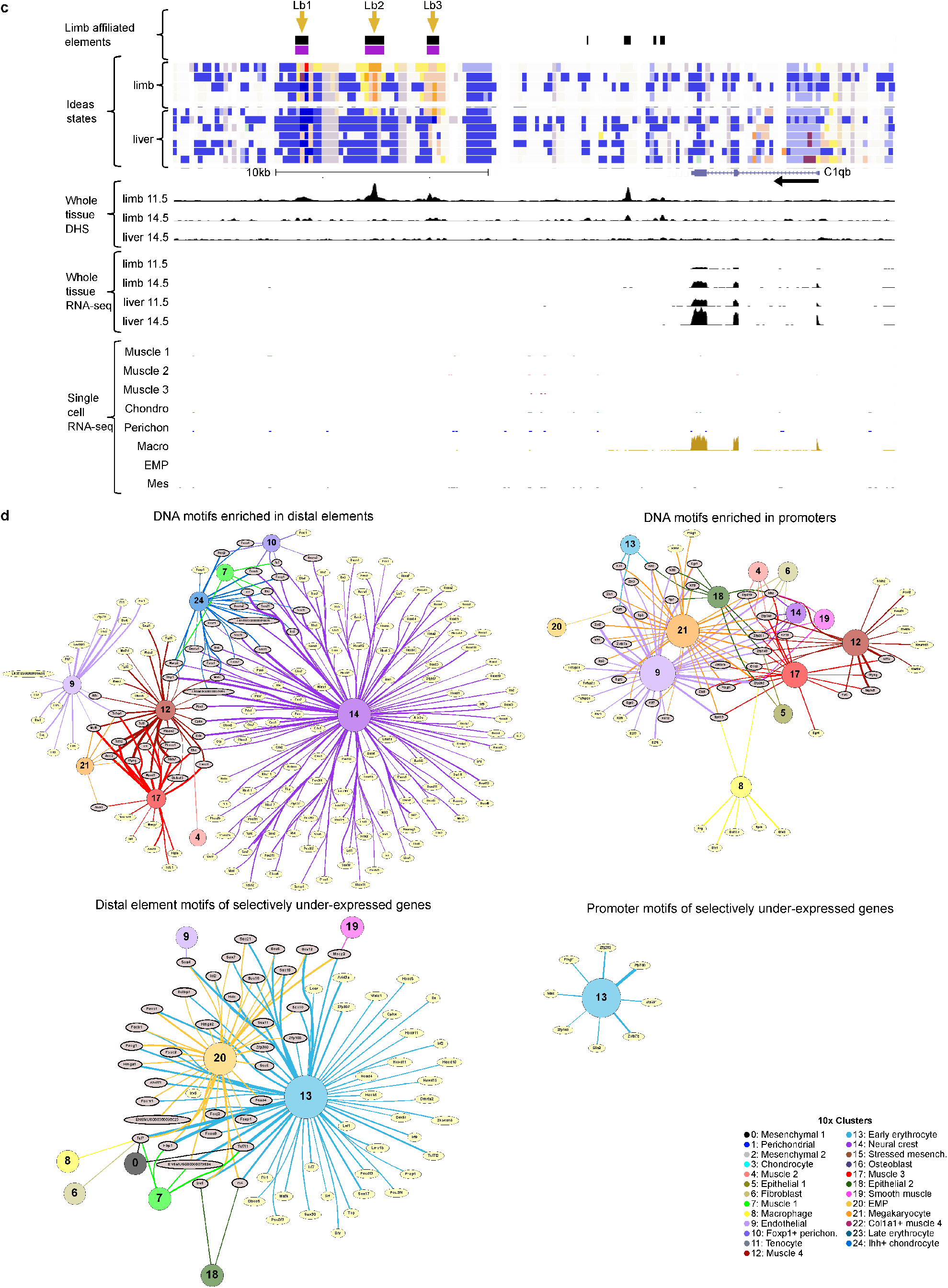

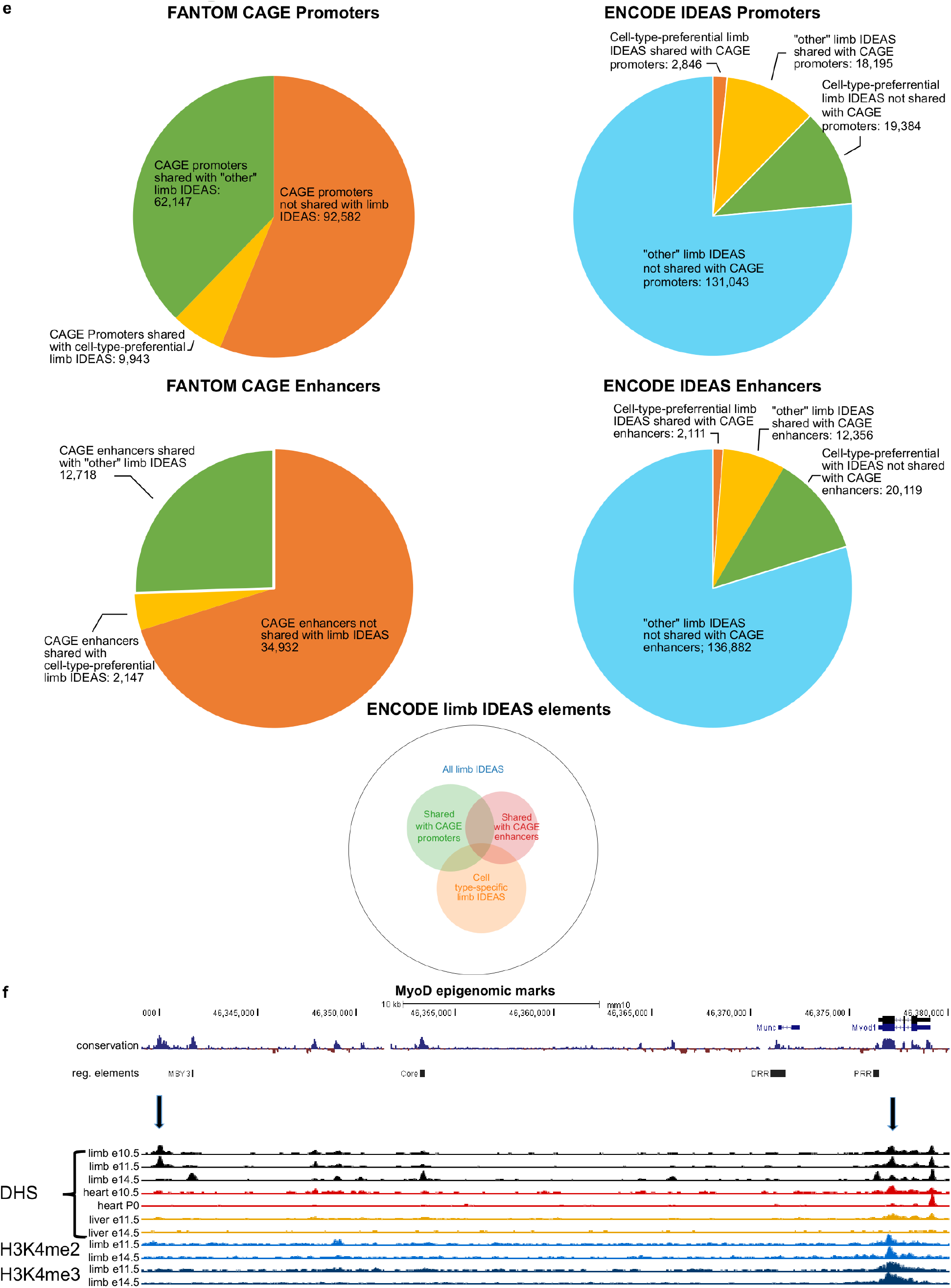

