## Supplementary Note for "The changing mouse embryo transcriptome at whole tissue and single-cell resolution"

### Supplementary Materials

#### Supplementary Note 1

##### Inferring cell types, states, and stages

Differentially expressed genes for each cell cluster were calculated separately for C1 and 10X datasets using Seurat's FindMarkers (`https://satijalab.org/seurat/seurat_clustering_tutorial_part2.html`) with `min.pct` = 0.25, applying its Wilcoxon rank sum test, and setting `min.diff.pct` to be 0.2 or 0.4.

The differential gene lists are provided in full in Supplementary Table 4, and representative top markers for each cluster are given in Tables SN1-1 and SN1-2. The top marker genes for each cell cluster are shown in Fig. 3c where each cell cluster was down-sampled to at most 100 cells for 10x data and at most 50 cells for C1 data.

To associate cell clusters with likely cell types/stages, the differential marker expression lists with `min.diff.pct` = 0.2 for each cluster were entered into FuncAssociate 3.0 (`http://llama.mshri.on.ca/funcassociate/`)[1], which returned GO attributes that were evaluated by LOD score and $p$ values, requiring that all terms used be $p_{adj} \leq 0.05$, corrected for multiple testing by empirical resampling. Attributes emphasizing terms that are explicitly or implicitly pertinent to developmental processes and cell types were selected by inspection and the appropriate attribute/entity tables were downloaded. The gene symbols from the attribute/entity tables were then used to search the literature to assemble curated marker genes and references (Tables SN1-1 and SN1-2). Key confirmatory literature markers for a candidate cell type were evaluated for significantly differential marker status (Supplementary Data 4) and included as literature markers. In cases where the literature provided a clear naming convention for a cell type association, we adopted that name in the tables. In other cases, the name is assigned from this work (e.g. *Ihh*+ chondrocyte). When marker ambiguity prevented an unequivocal cell type assignment, we chose a designation based on the weight of the available evidence that is consistent with known fate mapping, tracing and cell sorting studies.

Our cell type identification was performed independently for data from the two platforms. We then evaluated correlations between the clusters, labeled by their cell type assignments, by logistic regression using all the detected genes (Fig SN1). The core set of 11 major cell types and stages, displayed identity broadly across their entire transcriptomes (Fig 3). Cell types with multiple developmental stages or subdivisions show splitting of C1 state-specific genes across 2 or more 10x clusters (e.g. mus1, mus2, mus3 and mus4), as would be expected from differences in cell sampling depth and transcriptome-depth.

**Cell type sensitivity**. The greater cell sampling depth of the 10x Genomics data (~90,000 cells) detected 14 additional clusters with their own provisional cell-type assignments. Three of the 10x-only cell types were expected to be absent from C1 data due to platform-specific details of cell-size filtration (early and late erythrocytes) and sample preparation (prior removal of sticky epidermal cells). The remaining 10x only clusters include rarer cell types/states with no C1 equivalent, presumably due mainly to sparse cell sampling with C1 plus some additional subdivision of C1 types into multiple 10x types.

**Marker gene sensitivity**. For several of the best-studied cell types we found that the C1 data identified key known markers from the literature that were not in the corresponding 10x differential lists. Examples that are functionally important include *Pthlh*, a regulator of the proliferation/differentiation choice in early chondrocytes[2,3]; *Spry1*, which regulates quiescence in *Pax7* muscle precursor cells[4] that comprise the Mus2 cluster; and *Dmrt2*, which is directly regulated by Pax3 in muscle precursors (Mus1), and in turn directly regulates *Myf5*[5], which marks mus1 and mus2 in C1 data. None of these were significant markers in the corresponding 10x cell types, and all three were expressed at low levels in the corresponding C1 clusters (*Pthl*,: 6 copies per cell (cpc)); *Spry1*, 1 cpc; and *Dmrt2*, 3 cpc).

**Marker gene context**. Defining a marker gene or a multigene cell-type signature also depends heavily on the tissue sample context. This is a pertinent caution for future uses of the marker gene sets derived here, if they are applied to other studies in different biological settings. Conversely, this issue also informed our use of markers from the literature, where we favored the closest context available, although a strong match was often unavailable. Overall, our cell type identity assignments did prove consistent with classical and modern tracing studies and genetic knockouts for the better-studied cell types. We further observed that cell-type marker signatures increased in complexity as several lineages progressed (Fig. 3c and Extended Data Fig. 9 j), making the more mature types (e.g. Muscle 3 or Macrophage) easier to define with high confidence. By contrast, their progenitors (Muscle1, EMP and Mesenchyme1) displayed lower salience signatures, with few progenitor-unique genes compared with their more differentiated counterparts.

**Cluster structure sensitivity**. It is important to recognize that the relationship of a cell cluster with a dominant cell identity does not (and is not expected to) preclude the presence of additional cell types within the cluster. At this early level of single cell resolution, it is expected that much additional cell type structure is unresolved. For example, in the cell clusters identified here as predominantly EMP, close examination of candidate marker gene sets from the literature found some evidence for the presence of related cells (mast cells), as noted in Tables SN1-1 and SN1-2.

#### Supplementary Figures

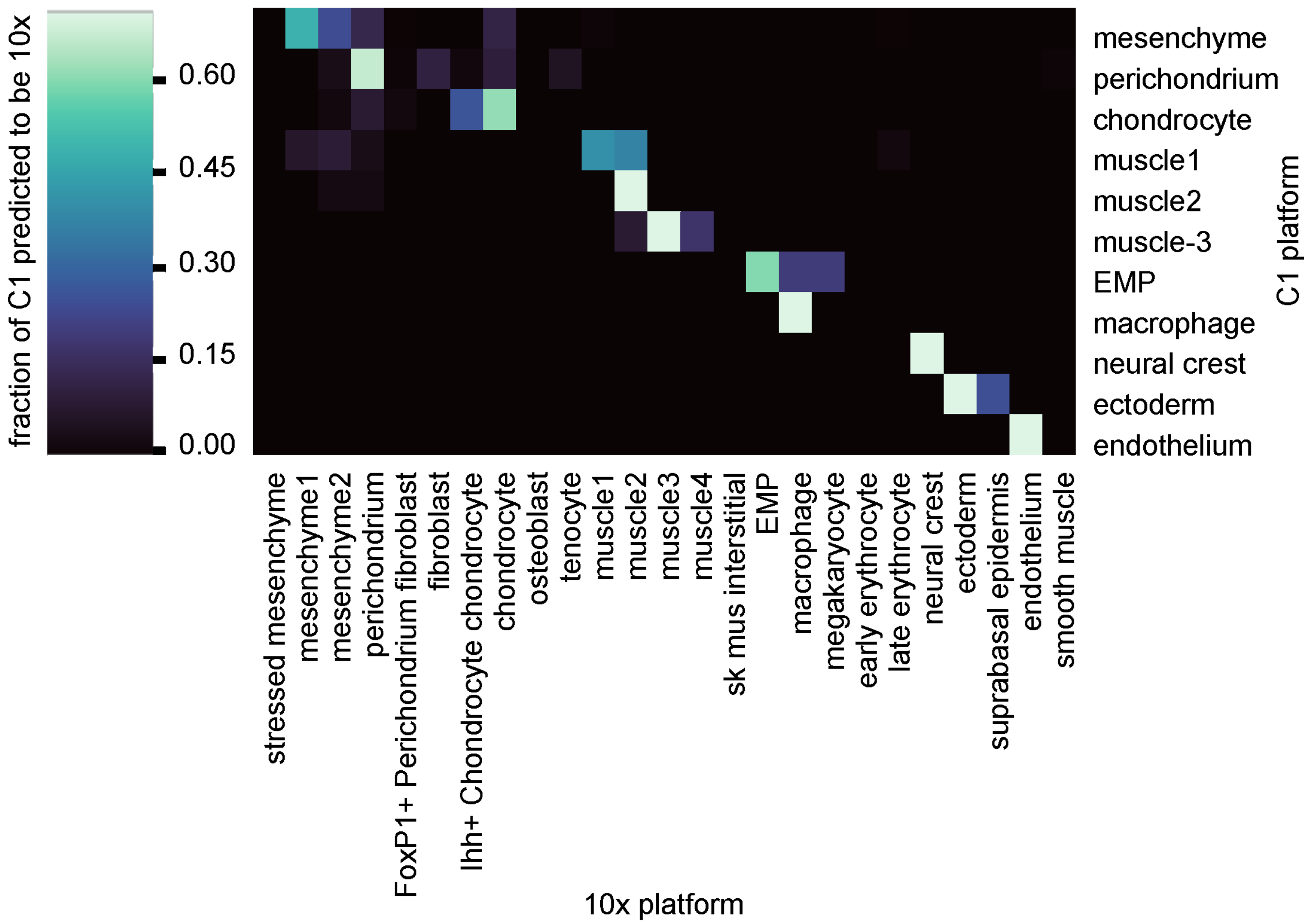


**Figure SN 1: Cross-comparison of cell type assignments across the two scRNA-seq platforms.**. Logistic regression results are plotted for all pairwise comparisons of cell type assignments across both scRNA-Seq platforms. Scale bar represents the fractional proportion of a C1 cell type predicted to be each 10x cell type. For cell types found in the 10x Genomics data that were not found in the C1 Fluidigm data, the scores are set at 0. Both 10x and C1 data were normalized, *log*-transformed and scaled before regression. To train a multinomial logistic regression model using the scikit-learn python package, the categorical information of the 25 cell types from 10x were used as the dependent variable. The trained model was then used to predict the C1 data. This model assigned each C1 single-cell into one of the 25 cell type categories defined by 10x data. The distribution (fraction) of the assignment per each C1 cell type into 10x categories was visualized as a row in the heatmap.

#### Supplementary Tables

**Table SN1 1: Fluidigm C1 marker genes and inferred cell identities, states and stages.** The statistical test for Gene Ontology analysis is a single-hypothesis $p$-value of the association between attribute and query (based on Fisher's Exact Test). $p_{adj}$: fraction (as a %) of 1000 null-hypothesis simulations having attributes with this single-hypothesis $p$ value or smaller. Sample sizes for C1 clusters are: mesenchyme, 1242 genes; perichondrium, 3108 genes; chondrocytes, 3756 genes, muscle 3, 2090 genes, muscle 1, 962 genes; neural crest, 2933 genes; EMP, 3463 genes; endothelial, 3145 genes; muscle 2, 2114; macrophage, 3022 genes; epithelial, 2625 genes. Sample sizes for 10x clusters are: Cluster 0, 398 genes; Cluster 1, 248 genes; Cluster 2, 70 genes; Cluster 3, 164 genes; Cluster 4, 146 genes; Cluster 5, 217 genes; Cluster 6, 262 genes; Cluster 7, 265 genes; Cluster 8, 734 genes; Cluster 9, 622 genes; Cluster 10, 160 genes; Cluster 11, 349 genes; Cluster 12, 556 genes; Cluster 13, 156 genes; Cluster 14, 334 genes; Cluster 15, 623 genes; Cluster 16, 542 genes; Cluster 17, 322 genes; Cluster 18, 431 genes; Cluster 19, 397 genes; Cluster 20, 592 genes; Cluster 21, 881 genes; Cluster 22, 117 genes; cluster 23, 42 genes; Cluster 24, 336 genes. $p$-values for GO term association are single-hypothesis $p$-values of the association between attribute and query (based on Fisher's Exact Test), and were adjusted by the fraction (as a %) of 1000 null-hypothesis simulations having attributes with this single-hypothesis P value or smaller. GO search sample sizes for C1 clusters are: mesenchymal, 90 genes; perichondrial, 369 genes; chondrocyte, 279 genes; muscle 3, 365 genes; muscle 1, 117 genes; neural crest, 165 genes; EMP, 393 genes; endothelial, 314 genes; muscle 2, 236 genes; macrophage, 632 genes; epithelial, 169 genes. GO search sample sizes for 10x clusters are: cluster 0, 177 genes; cluster 1, 116 genes; cluster 2, 16 genes; cluster 3, 30 genes, cluster 4, 100 genes, cluster 5, 158 genes, cluster 6, 173 genes, cluster 7, 153 genes, cluster 8, 679 genes; cluster 9, 561 genes; cluster 10, 50 genes; cluster 11, 170 genes; cluster 12, 516 genes; cluster 13, 149 genes; cluster 14, 305 genes; cluster 15, 2 genes; cluster 16, 450 genes, cluster 17, 263 genes, cluster 18, 380 genes, cluster 19, 323 genes; cluster 20, 520 genes; cluster 21, 833 genes; cluster 22, 60 genes; cluster 23, 38 genes; cluster 24, 205 genes.

| Cell Cluster # | cell cluster name | provisional cell type/state | GO ID | GO terms from Fluidigm data | LOD score | $p_{unadj}$ | $p_{adj}$. | top 5 genes from C1 clusters | selected literature marker genes |
|---|---|---|---|---|---|---|---|---|---|
| 0 | mesenchyme (mes) | mesenchyme | GO:0035115 | embryonic forelimb morphogenesis | 1.67 | 2.71E-07 | ≤0.001 | [*Mecom*, *Hsd11b2*, *Lix1*, *Car14*, *Hoxd10*] | [*Hoxd10* [6], *Hoxd11* [6], *Hoxd12* [6], *Msx1* [7], *Twi1* [8,9]] |
| | | | GO:0009954 | proximal/distal pattern formation | 1.67 | 4.69E-06 | 0.006 | | |
| | | | GO:0042733 | embryonic digit morphogenesis | 1.66 | 4.75E-12 | ≤0.001 | | |
| | | | GO:0009952 | anterior/posterior pattern specification | 1.15 | 3.95E-07 | ≤0.001 | | |
| 1 | perichondrium (pchon) | perichondrium | GO:0060351 | cartilage development involved in endochondral bone morphogenesis | 1.41 | 7.20E-06 | 0.032 | [*Col6a1*, *Egfl6*, *Creb3l1*, *Ogn*, *Col1a1*] | [*Col1a1* [10], *Dcn1* [1], *Dkk3* [12], *Ogn* [11], *Thbs2* [12]] |
| | | | GO:0030199 | collagen fibril organization | 1.27 | 1.33E-08 | ≤0.001 | | |
| | | | GO:0001649 | osteoblast differentiation | 0.98 | 7.79E-06 | 0.039 | | |

*Continued on next page*

| Cell Cluster # | cell cluster name | provisional cell type/state | GO ID | GO terms from Fluidigm data | LOD score | $p_{unadj}$ | $p_{adj}$. | top 5 genes from C1 clusters | selected literature marker genes |
|---|---|---|---|---|---|---|---|---|---|
| | | | GO:0001503 | ossification | 0.94 | 7.67E-09 | ≤0.001 | | |
| 2 | chondrocytes (chon) | immature chondrocytes | GO:0060351 | cartilage development involved in endochondral bone morphogenesis | 1.64 | 4.74E-08 | ≤0.001 | [*Susd5*, *Matn1*, *Foxa3*, *Ncmap*, *Acan*] | [*Acan* [3], *Col2a1* [3], *Dlx5* [13], Ihh3, *Pthlh* [3], *Runx2* [13], *Runx3* [14], *Sox5* [3], *Sox6* [3], *Sox9* [3], *Sp7* [13]] |
| | | | GO:0032331 | negative regulation of chondrocyte differentiation | 1.55 | 1.35E-07 | ≤0.001 | | |
| | | | GO:0001958 | endochondral ossification | 1.54 | 1.01E-09 | ≤0.001 | | |
| | | | GO:0001502 | cartilage condensation | 1.50 | 2.47E-06 | 0.006 | | |
| | | | GO:0030279 | negative regulation of ossification | 1.16 | 8.08E-11 | ≤0.001 | | |
| | | | GO:0001649 | osteoblast differentiation | 1.16 | 7.21E-08 | ≤0.001 | | |
| 3 | muscle3 (mus3) | myocyte | GO:0003009 | skeletal muscle contraction | 1.96 | 1.29E-18 | ≤0.001 | [*Gm7325*, *Ablim3*, *Kcnk13*, *Smyd1*, *Klhl41*] | [*Actc1* [15], *Act3* [15], *Fgfr4* [16], *Myod1* [16], *Myog* [16], *Myot* [15], *Ryr1* [15], *Tead4* [15]] |
| | | | GO:0048741 | skeletal muscle fiber development | 1.54 | 1.33E-10 | ≤0.001 | | |
| | | | GO:0007520 | myoblast fusion | 1.42 | 6.50E-07 | ≤0.001 | | |

*Continued on next page*

Table SN1 1 – *Continued from previous page*

| Cell Cluster # | cell cluster name | provisional cell type/state | GO ID | GO terms from Fluidigm data | LOD score | $p_{unadj}$ | $p_{adj.}$ | top 5 genes from C1 clusters | selected literature marker genes |
|---|---|---|---|---|---|---|---|---|---|
| 4 | muscle1 (mus1) | migratory limb muscle precursor cell | GO:0007517 | muscle organ development | 1.45 | 2.67E-10 | ≤0.001 | [*Pax3*, *Lbx1*, *Pitx2*, *Myod1*, *Eya1*] | [*Dmrt2*[5], *Eya1*[16], *Eya2*[16], *Lbx1*[16], *Met*[17], *Msc*[16], *Myf5*[16], *Myod1*[16], *Pax3*[18–20], *Pitx2*[16], *Pitx3*[16]] |
| | | | GO:0007519 | skeletal muscle tissue development | 1.50 | 1.32E-08 | ≤0.001 | | |
| | | | GO:0001756 | somitogenesis | 1.35 | 8.55E-06 | 0.018 | | |
| 5 | neural crest (neur) | neural crest | GO:0007422 | peripheral nervous system development | 1.62 | 8.84E-06 | 0.015 | [*Gpr17*, *Foxd3*, *St8sia5*, *Lgi4*, *Insc*] | [*Dlx1*[21], *Dlx2*[21], *Foxd3*[22], *Sox2*[22], *Sox10*[22], *Zeb2*[22]] |
| | | | GO:0042552 | myelination | 1.17 | 1.68E-06 | 0.003 | | |
| | | | GO:0045666 | positive regulation of neuron differentiation | 0.76 | 3.46E-07 | ≤0.001 | | |
| 6 | erythro-myeloid precursor (EMP)* | erythro-myeloid precursor | GO:0006909 | phagocytosis | 1.05 | 1.36E-07 | 0.001 | [*Slc22a3*, *1110028F11Rik*, *Ubash3a*, *Rab44*, *Gata1*] | [*Fcgr3*[23], *Gata1*[23], *Gata2*[23], *Gfi1b*[24], *Spi1*[23]] |
| | | | GO:0051707 | response to other organism | 0.53 | 3.08E-07 | 0.001 | | |
| | | | GO:0002478 | antigen processing and presentation of exogenous peptide antigen | 1.39 | 9.96E-07 | 0.003 | | |
| 7 | endothelium (endo) | endothelium | GO:0001945 | lymph vessel development | 1.81 | 9.26E-09 | ≤0.001 | [*Ccm2l*, *Myct1*, *Pcdh12*, *Sox17*, *Robo4*] | [*Aplnr*[25], *Flt4*[26], *Kdr*[26], *Lmo2*[27], *Sox17*[27]] |
| | | | GO:0002040 | sprouting angiogenesis | 1.43 | 5.46E-08 | ≤0.001 | | |
| | | | GO:0061028 | establishment of endothelial barrier | 1.39 | 8.34E-07 | ≤0.001 | | |

*Continued on next page*

Table SN1 1 – *Continued from previous page*

| Cell Cluster # | cell cluster name | provisional cell type/state | GO ID | GO terms from Fluidigm data | LOD score | $p_{unadj}$ | $p_{adj.}$ | top 5 genes from C1 clusters | selected literature marker genes |
|---|---|---|---|---|---|---|---|---|---|
| | | | GO:0001525 | angiogenesis | 1.17 | 1.67E-28 | ≤0.001 | | |
| 8 | muscle2 (mus2) | myoblast | GO:0007517 | muscle organ development | 1.31 | 1.57E-12 | ≤0.001 | [*Pax7*, *Ntrk1*, *Scn3b*, *Gm9947*, *Myf5*] | [*Dmrt2* [5], *En1* [28], *Eya1* [16], *Eya2* [16], *Hes6* [29], *Lbx1* [16], *Met* [17], *Msc* [16], *Myf5* [16], *Myod1* [16], *Notch3* [30], *Pax7* [31], *Pitx2* [16], *Pitx3* [16], *Six1* [16], *Six2* [16], *Sox8* [32], *Spry1* [4], *Vgll2* [33], *Vgll3* [34,35]] |
| | | | GO:0061061 | muscle structure development | 1.27 | 4.10E-12 | ≤0.001 | | |
| | | | GO:0051147 | regulation of muscle cell differentiation | 1.05 | 4.95E-12 | ≤0.001 | | |
| 9 | macrophage (mac) | tissue resident macrophage | GO:0098542 | defense response to other organism | 1.01 | 5.56E-44 | ≤0.001 | [*Lrrc25*, *Clec4a3*, *Cybb*, *Fcgr1*, *AI607873*] | [*Csf1r* [23], *Cx3cr1* [23], *Emr1* [23], *Fcer1g* [23], *Irf8* [23], *Runx1* [36], *Spi1* [23]] |
| | | | GO:0045087 | innate immune response | 1.03 | 1.00E-41 | ≤0.001 | | |
| | | | GO:0001819 | positive regulation of cytokine production | 0.95 | 2.31E-37 | ≤0.001 | | |
| | | | GO:0006935 | chemotaxis | 0.98 | 9.62E-28 | ≤0.001 | | |

*Continued on next page*

Table SN1 1 – *Continued from previous page*

| Cell Cluster # | cell cluster name | provisional cell type/state | GO ID | GO terms from Fluidigm data | LOD score | $p_{unadj}$ | $p_{adj}$. | top 5 genes from C1 clusters | selected literature marker genes |
|---|---|---|---|---|---|---|---|---|---|
| 10 | ectoderm (ecto) | ectoderm | GO:0005109 | frizzled binding | 1.55 | 1.13E-06 | 0.001 | [*4631405K08Rik*, *Ap1m2*, *Fermt1*, *Krt5*, *Krt14*] | [*Fzd6* [37], *Grhl2* [38], *Grhl3* [38], *Klf5* [39], *Krt1* [39], *Krt17* [39], *Wnt3* [37], *Wnt4* [37], *Wnt6* [37], *Wnt7a* [37], *Wnt7b* [37], *Wnt10a* [37]] |
| | | | GO:0035567 | non-canonical Wnt signaling pathway | 1.53 | 1.38E-06 | 0.002 | | |
| | | | GO:0030216 | keratinocyte differentiation | 1.41 | 5.53E-08 | ≤0.001 | | |
| | | | GO:0035136 | forelimb morphogenesis | 1.28 | 1.87E-05 | 0.038 | | |
| | | | GO:0016055 | Wnt signaling pathway | 0.89 | 2.29E-07 | ≤0.001 | | |
| | | | GO:0001228 | transcriptional activator activity, RNA polymerase II transcription regulatory region sequence-specific binding | 0.80 | 4.68E-08 | ≤0.001 | | |
| | | | GO:0007389 | pattern specification process | 0.68 | 1.45E-05 | 0.031 | | |
| | | | GO:0048646 | anatomical structure formation involved in morphogenesis | 0.61 | 9.81E-07 | 0.001 | | |

* 2 cells in this cluster expressed mast cell genes.

**Table SN1 2: 10x Genomics marker genes and inferred cell identities, states and stages.** The statistical test for Gene Ontology analysis is a single-hypothesis $p$-value of the association between attribute and query (based on Fisher's Exact Test). $p_{adj}$: fraction (as a %) of 1000 null-hypothesis simulations having attributes with this single-hypothesis $p$ value or smaller. Sample sizes for C1 clusters are: mesenchyme, 1242 genes; perichondrium, 3108 genes; chondrocytes, 3756 genes, muscle 3, 2090 genes, muscle 1, 962 genes; neural crest, 2933 genes; EMP, 3463 genes; endothelial, 3145 genes; muscle 2, 2114; macrophage, 3022 genes; epithelial, 2625 genes. Sample sizes for 10x clusters are: Cluster 0, 398 genes; Cluster 1, 248 genes; Cluster 2, 70 genes; Cluster 3, 164 genes; Cluster 4, 146 genes; Cluster 5, 217 genes; Cluster 6, 262 genes; Cluster 7, 265 genes; Cluster 8, 734 genes; Cluster 9, 622 genes; Cluster 10, 160 genes; Cluster 11, 349 genes; Cluster 12, 556 genes; Cluster 13, 156 genes; Cluster 14, 334 genes; Cluster 15, 623 genes; Cluster 16, 542 genes; Cluster 17, 322 genes; Cluster 18, 431 genes; Cluster 19, 397 genes; Cluster 20, 592 genes; Cluster 21, 881 genes; Cluster 22, 117 genes; cluster 23, 42 genes; Cluster 24, 336 genes. $p$-values for GO term association are single-hypothesis $p$-values of the association between attribute and query (based on Fisher's Exact Test), and were adjusted by the fraction (as a %) of 1000 null-hypothesis simulations having attributes with this single-hypothesis P value or smaller. GO search sample sizes for C1 clusters are: mesenchymal, 90 genes; perichondrial, 369 genes; chondrocyte, 279 genes; muscle 3, 365 genes; muscle 1, 117 genes; neural crest, 165 genes; EMP, 393 genes; endothelial, 314 genes; muscle 2, 236 genes; macrophage, 632 genes; epithelial, 169 genes. GO search sample sizes for 10x clusters are: cluster 0, 177 genes; cluster 1, 116 genes; cluster 2, 16 genes; cluster 3, 30 genes, cluster 4, 100 genes, cluster 5, 158 genes, cluster 6, 173 genes, cluster 7, 153 genes, cluster 8, 679 genes; cluster 9, 561 genes; cluster 10, 50 genes; cluster 11, 170 genes; cluster 12, 516 genes; cluster 13, 149 genes; cluster 14, 305 genes; cluster 15, 2 genes; cluster 16, 450 genes, cluster 17, 263 genes, cluster 18, 380 genes, cluster 19, 323 genes; cluster 20, 520 genes; cluster 21, 833 genes; cluster 22, 60 genes; cluster 23, 38 genes; cluster 24, 205 genes.

| Cell Cluster # | cell cluster name | provisional cell type/state | GO ID | GO terms from 10x Genomics data | LOD score | $p_{unadj}$ | $p_{adj}$. | top 5 genes from 10x clusters | selected literature marker genes |
|---|---|---|---|---|---|---|---|---|---|
| 0 | proximal mesenchyme (mesprox) | proximal mesenchyme | GO:0030326 | embryonic limb morphogenesis | 1.22 | 3.72E-09 | ≤0.001 | [*Lix1*, *Asb4*, *Igdcc3*, *Hmga2*, *Rrm2*] | [*Hoxa9*[6], *Hoxd9*[6], *Hoxd10*[6], *Hoxd11*[6], *Shox2*[40]] |
| | | | GO:0048704 | embryonic skeletal system morphogenesis | 1.06 | 7.90E-06 | 0.018 | | |
| | | | GO:0022402 | cell cycle process | 0.70 | 1.06E-08 | ≤0.001 | | |
| 1 | perichondrium (pchon) | perichondrium | GO:0005583 | fibrillar collagen trimer | 2.48 | 2.86E-12 | ≤0.001 | [*Dlk1*, *Meg3*, *Col3a1*, *Igf1*, *Col1a1*] | [*Col1a1*[10], *Col5a1*[41], *Igfbp3*[42], *Shox0*[40]] |
| | | | GO:0001649 | osteoblast differentiation | 1.44 | 3.22E-08 | 0.001 | | |
| | | | GO:0009653 | anatomical structure morphogenesis | 0.59 | 3.33E-07 | 0.002 | | |
| 2 | distal mesenchyme (mesdist) | distal mesenchyme | GO:0030326 | embryonic limb morphogenesis | 2.63 | 8.06E-21 | ≤0.001 | [*Msx1*, *Hoxd13*, *Prrx2*, *Hoxa11os*, *Hoxd12*] | [*Hoxa10*[6], *Hoxd12*[6], *Hoxd13*[6], *Msx1*[7], *Msx2*[7]] |

*Continued on next page*

| Cell Cluster # | cell cluster name | provisional cell type/state | GO ID | GO terms from 10x Genomics data | LOD score | $p_{unadj}$ | $p_{adj}$. | top 5 genes from 10x clusters | selected literature marker genes |
|---|---|---|---|---|---|---|---|---|---|
| | | | GO:0042733 | embryonic digit morphogenesis | 2.48 | 1.37E-14 | ≤0.001 | | |
| | | | GO:0007389 | pattern specification process | 1.74 | 1.31E-10 | ≤0.001 | | |
| | | | GO:0009952 | anterior/posterior pattern specification | 1.93 | 1.24E-09 | ≤0.001 | | |
| | | | GO:0051216 | cartilage development | 2.12 | 2.98E-09 | ≤0.001 | | |
| 3 | chondrocytes (chon) | immature chondrocytes | GO:0060174 | limb bud formation | 2.54 | 3.21E-07 | ≤0.001 | [*Sox9*, *Col2a1*, *Wwp2*, *Col9a1*, *Gdf5*] | [*Acan* [3,14], *Col2a1* [3,14], *Sox5* [3,14], *Sox6* [3,14], *Sox9* [3,14]] |
| | | | GO:0002063 | chondrocyte development | 2.38 | 1.09E-08 | ≤0.001 | | |
| | | | GO:0001503 | ossification | 1.63 | 4.41E-07 | 0.001 | | |
| 4 | muscle2 (mus2) | early myoblast | GO:0007519 | skeletal muscle tissue development | 1.57 | 4.40E-09 | ≤0.001 | [*Msc*, *Kcne1l*, *Itm2a*, *Pdgfa*, *Vgll2*] | [*Msc1* [6], *Myf5* [16], *Myod1* [16], *Pax7* [31], *Pitx2* [16], *Six1* [16], *Vgll2* [33]] |
| | | | GO:0014706 | striated muscle tissue development | 1.39 | 7.43E-09 | ≤0.001 | | |
| | | | GO:0007389 | pattern specification process | 0.93 | 3.52E-08 | ≤0.001 | | |
| 5 | ectoderm (ecto) | ectoderm | GO:0010482 | regulation of epidermal cell division | 3.02 | 3.11E-07 | ≤0.001 | [*Cxcl14*, *Pdgfa*, *Wnt6*, *Krt14*, *Gjb2*] | [*Cxcl14* [43], *En1* [44], *Fzd6* [37], *Fzd10* [37], *Wnt4* [37], *Wnt6* [37], *Wnt7b* [37], *Wnt10a* [37]] |
| | | | GO:0003334 | keratinocyte development | 1.85 | 1.42E-06 | 0.001 | | |
| | | | GO:0043588 | skin development | 1.67 | 1.39E-15 | ≤0.001 | | |
| | | | GO:0009954 | proximal/distal pattern formation | 1.53 | 1.44E-06 | 0.001 | | |

*Continued on next page*

Table SN1 2 – *Continued from previous page*

| Cell Cluster # | cell cluster name | provisional cell type/state | GO ID | GO terms from 10x Genomics data | LOD score | $p_{unadj}$ | $p_{adj}$. | top 5 genes from 10x clusters | selected literature marker genes |
|---|---|---|---|---|---|---|---|---|---|
| 6 | fibroblast (fibro) | fibroblast | GO:0048407 | platelet-derived growth factor binding | 2.14 | 1.41E-10 | ≤0.001 | [*Crabp1*, *Crip1*, *Twist2*, *Lum*, *Rspo1*] | [*Irx1* [45], *Irx2* [45], *Irx3* [45], *Irx5* [45], *Tbx15* [46], *Twist2* [47]] |
| | | | GO:0045667 | regulation of osteoblast differentiation | 1.11 | 5.88E-09 | ≤0.001 | | |
| | | | GO:0030278 | regulation of ossification | 0.97 | 1.06E-08 | ≤0.001 | | |
| | | | GO:0048705 | skeletal system morphogenesis | 0.97 | 7.70E-06 | 0.011 | | |
| 7 | muscle1 (mus1) | migratory limb muscle precursor cells | GO:0014706 | striated muscle tissue development | 1.13 | 3.02E-06 | 0.007 | [*Pax3*, *Lbx1*, *Tcf15*, *Ppp1r14b*, *Slc25a5*] | [*Lbx1* [16], *Met* [17], *Pax3* [18–20]] |
| | | | GO:0048562 | embryonic organ morphogenesis | 1.06 | 3.72E-07 | ≤0.001 | | |
| | | | GO:0060537 | muscle tissue development | 1.05 | 9.13E-06 | 0.021 | | |
| 8 | macrophage (mac) | macrophage | GO:0048246 | macrophage chemotaxis | 1.41 | 1.52E-06 | 0.002 | [*Apoe*, *Fcer1g*, *Tyrobp*, *C1qb*, *C1qc*] | [*Aif1* [23], *Cx3cr1* [23], *Emr1* [23], *Csf1r* [23], *Fcgr1* [23], *Fcgr3* [23], *Grn* [23], *Irf8* [23], *Maf* [23], *Spi1* [23], *Zeb2* [23]] |
| | | | GO:0019884 | antigen processing and presentation of exogenous antigen | 1.40 | 6.53E-11 | ≤0.001 | | |
| | | | GO:0006911 | phagocytosis, engulfment | 1.19 | 1.57E-08 | ≤0.001 | | |
| 9 | endothelium (endo) | endothelium | GO:0001945 | lymph vessel development | 1.55 | 2.91E-07 | ≤0.001 | [*Gng11*, *S100a16*, *Egfl7*, *Cdh5*, *Crip2*] | [*Aplnr* [25], *Efnb2* [48], *Ets1* [49], *Ets2* [49], *Flt4* [26], *Kdr* [26], *Sox17* [27]] |

*Continued on next page*

Table SN1 2 – *Continued from previous page*

| Cell Cluster # | cell cluster name | provisional cell type/state | GO ID | GO terms from 10x Genomics data | LOD score | $p_{unadj}$ | $p_{adj.}$ | top 5 genes from 10x clusters | selected literature marker genes |
|---|---|---|---|---|---|---|---|---|---|
| | | | GO:0045446 | endothelial cell differentiation | 1.35 | 2.77E-07 | ≤0.001 | | |
| | | | GO:1903672 | positive regulation of sprouting angiogenesis | 1.24 | 1.14E-06 | ≤0.001 | | |
| 10 | FoxP + perichondrium (pchon-FoxP+) | perichondrial cells expressing FoxP1 | GO:0030199 | collagen fibril organization | 1.98 | 3.15E-10 | ≤0.001 | [*Foxp1*, *Foxf2*, *Foxq1*, *Rprm*, *Pid1*] | [*Col1a1* [10], *Emx2* [50], *FoxP1* [51], *Lmx1b* [50,52]] |
| | | | GO:0001501 | skeletal system development | 1.32 | 1.15E-05 | 0.021 | | |
| | | | GO:0009653 | anatomical structure morphogenesis | 0.84 | 5.30E-08 | ≤0.001 | | |
| 11 | tenocytes (teno) | tenocytes | GO:0035989 | tendon development | 2.98 | 3.88E-07 | 0.002 | [*Col1a1*, *Tnmd*, *Col1a2*, *Scx*, *Ogn*] | [*Col5a1* [53], *Col5a2* [53], *Col11a1* [53], *Mkx* [54], *Scx* [55], *Selm* [56], *Thbs4* [55], *Tnmd* [55]] |
| | | | GO:0005583 | fibrillar collagen trimer | 2.31 | 2.93E-11 | ≤0.001 | | |
| | | | GO:0061448 | connective tissue development | 1.24 | 4.21E-06 | 0.007 | | |
| 12 | muscle4 (mus4) | myocyte | GO:0035995 | detection of muscle stretch | 2.60 | 2.43E-07 | 0.001 | [*Mylpf*, *Myl1*, *Actc1*, *Tnnc1*, *Myl4*] | [*Actc1* [15], *Chrng* [15], *Jsrp1* [15], *Myl1* [15], *Mylpf* [15], *Ryr1* [15], *Sln* [15], *Tnnc1* [15], *Tnnt1* [15]] |
| | | | GO:0070296 | sarcoplasmic reticulum calcium ion transport | 1.98 | 2.95E-10 | ≤0.001 | | |
| | | | GO:0003009 | skeletal muscle contraction | 1.81 | 1.18E-16 | ≤0.001 | | |

*Continued on next page*

Table SN1 2 – *Continued from previous page*

| Cell Cluster # | cell cluster name | provisional cell type/state | GO ID | GO terms from 10x Genomics data | LOD score | $p_{unadj}$ | $p_{adj}$. | top 5 genes from 10x clusters | selected literature marker genes |
|---|---|---|---|---|---|---|---|---|---|
| 13 | early erythrocyte (eryth1) | early erythrocyte | GO:0005833 | hemoglobin complex | 3.38 | 3.95E-16 | ≤0.001 | [*Hba-x*, *Hbb-bh1*, *Hba-a1*, *Hba-a2*, *Hbb-bt*] | [*Gata1*[57], *Hbb-y*[57], *Klf1*[57], *Lyl1*[57], *Spta1*[57], *Sptb*[57], *Tal1*[57], *Zfpm1*[57]] |
| | | | GO:0048821 | erythrocyte development | 1.98 | 7.18E-13 | ≤0.001 | | |
| | | | GO:0030097 | hemopoiesis | 1.13 | 5.32E-07 | 0.002 | | |
| 14 | neural crest (neur) | neural crest | GO:0014044 | Schwann cell development | 1.72 | 5.81E-06 | 0.017 | [*Ednrb*, *Arpc1b*, *Plp1*, *Fabp7*, *Phactr1*] | [*Ets1*[22], *Foxd3*[22], *Lmo4*[58], *Metrn*[59], *Pax3*[22], *Sox5*[22], *Sox10*[22], *Zeb2*[22]] |
| | | | GO:0001755 | neural crest cell migration | 1.25 | 6.86E-07 | 0.002 | | |
| | | | GO:0010001 | glial cell differentiation | 1.19 | 4.66E-11 | ≤0.001 | | |
| 15 | (mesX) | mesenchymal cells expressing cell stress genes | GO:0034663 | endoplasmic reticulum chaperone complex | 1.84 | 1.17E-10 | ≤0.001 | | [*Dnajc10*[60], *Hspa5*[61], *Hsp90*[62]] |
| | stressed mesenchyme | GO:0098803 | respiratory chain complex | 1.19 | 6.90E-13 | ≤0.001 | | | |
| | | | GO:0045454 | cell redox homeostasis | 1.10 | 1.82E-10 | ≤0.001 | | |
| 16 | osteoblast (ost) | osteoblast | GO:0001958 | endochondral ossification | 1.40 | 1.80E-09 | ≤0.001 | [*Ibsp*, *Ifitm5*, *Smpd3*, *Sgms2*, *Sp7*] | [*Col1a1*[10], *Dlx5*[13], *Mef2c*[63], *Pth1r*[64], *Runx2*[13], *Sp7*[13]] |
| | | | GO:0002062 | chondrocyte differentiation | 1.21 | 7.45E-09 | ≤0.001 | | |
| | | | GO:0001649 | osteoblast differentiation | 1.04 | 3.86E-08 | 0.001 | | |

*Continued on next page*

| Cell Cluster # | cell cluster name | provisional cell type/state | GO ID | GO terms from 10x Genomics data | LOD score | $p_{unadj}$ | $p_{adj}$. | top 5 genes from 10x clusters | selected literature marker genes |
|---|---|---|---|---|---|---|---|---|---|
| 17 | muscle3 (mus3) | late myoblast | GO:0003009 | skeletal muscle contraction | 1.88 | 1.71E-14 | ≤0.001 | [*Myog*, *Actc1*, *Tnnt1*, *Acta2*, *Vgll2*] | [*Dll1* [65], *Mef2c* [16], *Myod1* [16], *Myog* [16], *Pitx2* [16], *Six1* [16], *Sox8* [32], *Tnnc1* [15], *Tnnt1* [15], *Vgll2* [33], *Zbtb18* [66]] |
| | | | GO:0007519 | skeletal muscle tissue development | 1.36 | 3.09E-11 | ≤0.001 | | |
| | | | GO:0045661 | regulation of myoblast differentiation | 1.24 | 1.78E-08 | ≤0.001 | | |
| 18 | suprabasal epithelium (sup epi) | suprabasal epithelium | GO:0061436 | establishment of skin barrier | 1.75 | 3.75E-13 | ≤0.001 | [*Krtdap*, *Perp*, *Hspb1*, *Sfn*, *Krt5*] | [*Krt1* [39], *Krt10* [39], *Krt14* [39], *Krtdap* [39], *Notch1* [39], *Sbsn* [39], *Trp63* [39]] |
| | | | GO:0003334 | keratinocyte development | 1.72 | 2.91E-08 | ≤0.001 | | |
| | | | GO:0008544 | epidermis development | 1.31 | 2.35E-12 | ≤0.001 | | |
| 19 | smooth muscle (smm) | smooth muscle | GO:0014910 | regulation of smooth muscle cell migration | 1.04 | 1.49E-07 | ≤0.001 | [*Acta2*, *Rgs5*, *Tagln*, *Ndufa4l2*, *Rasgrp2*] | [*Acta2* [67], *Cav1* [68], *Coro1b* [69], *Cspg4* [68], *Cyr61* [68], *Egr1* [68], *Jag1* [68], *Pten* [67], *Tagln* [67], *Tagln2* [67]] |
| | | | GO:0001525 | angiogenesis | 0.93 | 2.08E-14 | ≤0.001 | | |
| | | | GO:0048514 | blood vessel morphogenesis | 0.91 | 1.98E-06 | 0.006 | | |
| | | | GO:0060537 | muscle tissue development | 0.88 | 3.59E-06 | 0.012 | | |

*Continued on next page*

Table SN1 2 – *Continued from previous page*

| Cell Cluster # | cell cluster name | provisional cell type/state | GO ID | GO terms from 10x Genomics data | LOD score | $p_{unadj}$ | $p_{adj}$. | top 5 genes from 10x clusters | selected literature marker genes |
|---|---|---|---|---|---|---|---|---|---|
| 20 | erythro-myeloid precursor (EMP) * | erythro-myeloid precursor | GO:0030099 | myeloid cell differentiation | 0.90 | 9.59E-14 | ≤0.001 | [*Cma1*, *Srgn*, *Cpa3*, *Rac2*, *Tyrobp*] | [*Cd34*[70], *Fcgr3*[23], *Gata1*[23], *Gata2*[23], *Kit*[23], *Spi1*[23]] |
| | | | GO:0050764 | regulation of phagocytosis | 0.85 | 2.17E-06 | 0.009 | | |
| | | | GO:0002821 | positive regulation of adaptive immune response | 0.79 | 2.80E-06 | 0.011 | | |
| | | | GO:0002819 | regulation of adaptive immune response | 0.79 | 9.95E-09 | ≤0.001 | | |
| 21 | megakaryocyte (meg) | megakaryocyte | GO:0030220 | platelet formation | 1.57 | 7.03E-12 | ≤0.001 | [*Pf4*, *Ppbp*, *Ctla2a*, *Gp1bb*, *Gp9*] | [*Fli1*[71], *Gata1*[71], *Nfe2*[71], *Mef2c*[71], *Meis1*[72], *Runx1*[71]] |
| | | | GO:0070527 | platelet aggregation | 1.29 | 4.23E-09 | ≤0.001 | | |
| | | | GO:0035855 | megakaryocyte development | 1.25 | 1.70E-06 | 0.006 | | |
| 22 | interstitial fibroblast (int/mus) | interstitial fibroblast | GO:0005861 | troponin complex | 3.33 | 4.35E-18 | ≤0.001 | [*Mylpf*, *Acta1*, *Myl1*, *Myl4*, *Sln*] | [*Col3a1*[73], *Col6a1*[73], *Col6a3*[73], *Cxcl12*[73], *Lum*[73], *Osr1*[73], *Vim*[73]] |
| | | | GO:0005583 | fibrillar collagen trimer | 2.45 | 8.44E-09 | ≤0.001 | | |
| | | | GO:0003009 | skeletal muscle contraction | 2.12 | 2.50E-09 | ≤0.001 | | |
| 23 | late erythrocyte (eryth2) | late erythrocyte | GO:0031722 | hemoglobin beta binding | 3.50 | 2.62E-06 | 0.008 | [*Snca*, *Alas2*, *Car2*, *Gypa*, *Ube2l6*] | [*Bpgm*[57], *Hbb-bt*[57], *Hbb-bs*[57], *Hbb-bh1*[57], *Hba-a2*[57], *Hba-a1*[57]] |

*Continued on next page*

Table SN1 2 – *Continued from previous page*

| Cell Cluster # | cell cluster name | provisional cell type/state | GO ID | GO terms from 10x Genomics data | LOD score | $p_{unadj}$ | $p_{adj}$. | top 5 genes from 10x clusters | selected literature marker genes |
|---|---|---|---|---|---|---|---|---|---|
| | | | GO:0031721 | hemoglobin alpha binding | 3.18 | 1.63E-08 | ≤0.001 | | |
| | | | GO:0048821 | erythrocyte development | 2.07 | 6.14E-06 | 0.01 | | |
| 24 | Ihh + chondrocyte (chonIhh+) | chondrocyte expressing Ihh | GO:0005594 | collagen type IX trimer | 2.90 | 6.82E-07 | 0.003 | [*Col2a1*, *Col11a1*, *Hapln1*, *Col9a3*, *Col9a1*] | [*Acan*[3], *Col2a1*[3], *Ihh*[3], *Mef2c*[3], *Pth1r*[74], *Runx2*[74], *Sox9*[14]] |
| | | | GO:0002063 | chondrocyte development | 1.81 | 2.69E-10 | ≤0.001 | | |
| | | | GO:0060351 | cartilage development involved in endochondral bone morphogenesis | 1.78 | 7.57E-09 | ¡0.001 | | |
| | | | GO:0045667 | regulation of osteoblast differentiation | 1.08 | 2.97E-09 | ≤0.001 | | |

* Two markers of mast cells that are not expressed in macrophages are seen in this cluster.
